# Long-Read Genome Assembly and Gene Model Annotations for the Rodent Malaria Parasite *Plasmodium yoelii* 17XNL

**DOI:** 10.1101/2023.01.06.523040

**Authors:** Mitchell J. Godin, Aswathy Sebastian, Istvan Albert, Scott E. Lindner

## Abstract

Malaria causes over 200 million infections and over 600 thousand fatalities each year, with most cases attributed to a human-infectious *Plasmodium* species, *Plasmodium falciparum*. Many rodent-infectious *Plasmodium* species, like *Plasmodium berghei, Plasmodium chabaudi*, and *Plasmodium yoelii*, have been used as genetically tractable model species that can expedite studies of this pathogen. In particular, *P. yoelii* is an especially good model for investigating the mosquito and liver stages of parasite development because key attributes closely resemble those of *P. falciparum*. Because of its importance to malaria research, in 2002 the 17XNL strain of *P. yoelii* was the first rodent malaria parasite to be sequenced. While sequencing and assembling this genome was a breakthrough effort, the final assembly consisted of >5000 contiguous sequences that impacted the creation of annotated gene models. While other important rodent malaria parasite genomes have been sequenced and annotated since then, including the related *P. yoelii* 17X strain, the 17XNL strain has not. As a result, genomic data for 17X has become the *de facto* reference genome for the 17XNL strain while leaving open questions surrounding possible differences between the 17XNL and 17X genomes. In this work, we present a high-quality genome assembly for *P. yoelii* 17XNL using HiFi PacBio long-read DNA sequencing. In addition, we use Nanopore long-read direct RNA-seq and Illumina short-read sequencing of mixed blood stages to create complete gene models that include not only coding sequences but also alternate transcript isoforms, and 5’ and 3’ UTR designations. A comparison of the 17X and this new 17XNL assembly revealed biologically meaningful differences between the strains due to the presence of coding sequence variants. Taken together, our work provides a new genomic and gene expression framework for studies with this commonly used rodent malaria model species.

## Introduction

Malaria remains a major global health burden (WHO Malaria Report 2022, (1)), with most of the 600,000 fatalities resulting from infection by human-infectious *Plasmodium falciparum*. The use of rodent-infectious model species has been instrumental to better understand those species that cause human disease due to high levels of genetic and physiological conservation across species (2). Researchers have routinely used these rodent model species, such as *P. yoelii, P. berghei*, and *P. chabaudi*, to investigate the entire *Plasmodium* life cycle, as genetic manipulations have long been rapid and rigorous in these species (2). We and others study *P. yoelii*, which is an especially good model for the mosquito and liver stages of *P. falciparum* parasite development (2). This is partly because *P. yoelii* mosquito stage parasites develop at a similar pace as do those of *P. falciparum*, and their sporozoites are less promiscuous than *P. berghei* sporozoites (2). Because of this, many studies of genetically attenuated parasite (GAP) vaccine candidates based upon sporozoites have recently included the use of *P. yoelii* as a pre-clinical model system (3). In support of this, large-scale analyses of gene expression of *P. yoelii* now match those available for *P. berghei* in many ways (4-9). For these reasons, *P. yoelii* has been an important malaria parasite used as a proxy for *P. falciparum* in pre-clinical and discovery phase studies.

Intuitively, genetic studies of any species are best conducted with accurate genome assemblies and gene models. Therefore, several species of *Plasmodium* parasites were the subject of early whole-genome sequencing efforts in the late 1990s and early 2000s (10, 11). This work provided a genome assembly of the human-infectious *P. falciparum* parasite with 14 nuclear chromosomes and the two organellar genomes of its mitochondrion and apicoplast (11). In addition, gene models for *P. falciparum* were annotated with introns/exons, with further improvements establishing 5’/3’ untranslated regions (UTRs) and transcript isoforms (12, 13). Similarly, the rodent-infectious *Plasmodium berghei* ANKA parasite was originally sequenced in 2005, resulting in a genome assembly with 7,497 contiguous sequences (contigs) that were later reduced to 16 contigs with a hybrid Illumina and 454 sequencing approach in 2014, and then further refined using PacBio sequencing in 2016 (14-16). Prior to this, the non-lethal *P. yoelii* 17XNL strain was the first rodent malaria parasite sequenced in 2002, which used ABI3700 sequencers and yielded a genome assembly of over 5,000 contigs (10). The *P. yoelii* 17X strain, from which 17XNL was derived, was sequenced in 2014 alongside PbANKA using the same Illumina and 454 sequencing approach to similarly establish a 16 contig genome assembly (15). With the advent of more accurate long-read sequencing technologies, there has been a renewed interest in sequencing the *Plasmodium* genomes and transcriptomes, including those of another *P. yoelii* strain, PyN67, which has been used to study genetic polymorphisms and drug responses (17). In addition, the genomes of other apicomplexan parasites, such as *Cryptosporidium* and *Babesia* species, have now been established using a combination of long-read Nanopore sequencing and short-read Illumina sequencing (18-20).

Although their genomes have been updated and are conveniently provided on PlasmoDB.org, Py17X and PbANKA have gene models that largely reflect the coding sequences, but not their UTRs despite the availability of RNA-seq data that could be used to approximate them (4-9, 15, 21-25). Finally, while the 17XNL strain of *P. yoelii* remains a highly used laboratory strain worldwide, its reference genome and gene models have not been revisited since 2002, and thus its genome assembly and gene models remain highly fragmented and incomplete. As a result, most researchers use the genome assembly and gene models of the related *P. yoelii* 17X strain as a proxy when working with the 17XNL strain and must operate under the assumption that the genomes of the two strains are effectively the same. However, this prompts a few important questions. How similar are the 17X and 17XNL strains? In what ways are they truly suitable proxies for one another? Given the state of the 17XNL genome assembly and the limited gene models available for both strains, these questions could not be accurately addressed. However, these kinds of questions can now be more rigorously addressed with the inclusion of long-read DNA sequencing. The long sequence reads produced by PacBio and Nanopore approaches better facilitate the scaffolding of long, contiguous sequences in a *de novo* assembly, even for complex genomes that have extreme AT-content and/or high degrees of repetitiveness, such as found with *Plasmodium* species (26, 27). Additionally, by combining long-read and short-read sequencing, the strengths of each can be used to polish the assembly to reduce systematic errors introduced by each of the different methodologies.

Therefore, here we have created a high-quality reference genome and gene model annotation for the *P. yoelii* 17XNL strain that we have used to address these outstanding questions. We utilized HiFi PacBio DNA-seq to create a Py17XNL reference genome with 16 high confidence/high accuracy contigs. Even without any polishing efforts, this approach outperformed a parallel effort using a hybrid Nanopore long-read DNA-seq/Illumina short-read DNA-seq method by several key metrics, including its assembly quality and the reduction of gaps. Furthermore, we created gene annotations for genes transcribed in asexual and sexual blood stages using a combination of Nanopore direct RNA-seq and our pre-existing Illumina RNA-seq datasets. These annotations include definitions of introns, exons, 5’, and 3’ UTRs, and transcript isoforms expressed in asexual and sexual blood stages. Using these data, we compared the genomic variance between the Py17XNL and Py17X strains to gain insight into the differences between the two strains and identified that most sequence variants reside in intergenic regions, whilst variation in the coding sequence of a select few genes could result in meaningful changes in Py17XNL parasite biology.

## Results

### A Comparison of Genome Assembly Approaches: PacBio HiFi vs. Nanopore/Illumina Hybrid Sequencing

*P. yoelii* 17XNL remains a commonly studied rodent malaria strain. Yet, its genome assembly remains highly fragmented and consists of over 5000 contigs as generated in 2002 (10). Consequently, most researchers use the reference genome of the related Py17X strain as a substitute for Py17XNL without knowing how appropriate it is to use it as a genomic proxy (15). To resolve these questions, we created a high-quality genome assembly of *P. yoelii* 17XNL Clone 1.1 obtained from BEI Resources, which is the common origin of this strain of parasites for many laboratories. Because several sequencing methodologies are now commonly used to assemble whole genomes, we used Nanopore, PacBio, and Illumina sequencing with DNA-or RNA-based libraries to determine an optimal approach to create a genome assembly with associated gene models for *P. yoelii* 17XNL. Nanopore ligation-based long-read DNA sequencing is currently favored by many researchers as it can provide extremely long sequence reads, resolve long stretches of repetitive regions, and assemble long structural variants in the genome (26, 27). HiFi PacBio DNA sequencing provides very high accuracy due to the sequencing of circular consensus sequences (ccs) of ∼10kb DNA fragments, providing a middle ground between the sequencing sizes provided by Illumina and Nanopore sequencing (27). We explored several data analysis protocols for combining data from different platforms to optimize this genome assembly. As widely available Nanopore sequencing chemistries (Q10) yield a systematic error, Nanopore data are often paired with Illumina data to provide error correction. The sequencing error rates from Illumina are typically above 99.9% and can be used with polishing algorithms to identify errors in assemblies that were produced with long, noisy reads (18, 19). A detailed outline of our experimental methods is included in Supplemental Figure 1. Briefly, swiss webster outbred mice were infected with Py17XNL Clone 1.1 parasites that had been passaged only once following receipt from BEI Resources to create a genome assembly reflective of the current stocks available in the depository. Upon reaching 1-3% parasitemia, mice were euthanized, white cells were depleted by cellulose, and red blood cells (RBCs) were lysed by saponin. The parasite pellets were used to produce high molecular weight genomic DNA using the NEB Monarch HMW DNA Extraction Kit for Cells and Blood as previously described (28). DNA purity, quantity, and fragment lengths were determined to all be high quality by NanoDrop, Qubit, and TapeStation measurements, respectively (Supplementary Table 1, Supplementary Figure 2)). This approach yielded DNA fragments of higher quality and higher molecular weight than the Qiagen QIAamp DNA Blood Mini Kit that is routinely used in our laboratory and in others. Matched gDNA samples were sequenced on a Nanopore MinION R9.4.1 flow cell using the ligation sequencing kit, as well as on an Illumina NextSeq 550 using the Illumina DNA PCR-Free kit. In parallel, Py17XNL HMW gDNA was sequenced on a PacBio Sequel using a PacBio SMRT cell. The raw reads from the PacBio sequencing run were converted into circular consensus sequences using the CCS algorithm.

To assess the quality of both Nanopore sequencing runs, we utilized Nanoplot, a quality control plotting suite specifically for long-read sequencing data (Supplemental Figure 3, Supplemental Table 2)) (29). The Nanopore sequencing runs for both replicate one and two resulted in an overall average read length of 16,706 bases, with an average Qscore of 11.3 (Supplemental Table 2). During this study, an improved high-accuracy base calling algorithm from Nanopore was released, which we also tested to see if it could improve our read quality. Upon re-basecalling the fast5 files, we saw a considerable increase in Qscore, from 11.3 to 14.1, even without the quality score filter that is imposed with the default, fast basecalling algorithm (Supplemental Figure 3, Supplemental Table 2). Despite this improvement in quality, there were no significant differences in the mean read length or the throughput (Supplemental Table 2). Using PacBio ccs reads, we saw an improvement in accuracy to an average Qscore of 36.3 (Figure 1A). The biggest difference came with throughput, which increased to 1,660,222,360 bases from 707,945,539 bases whilst still maintaining an average read length of 5,712.7 bases (Figure 1B, Supplemental Table 2).

**Figure 1:**
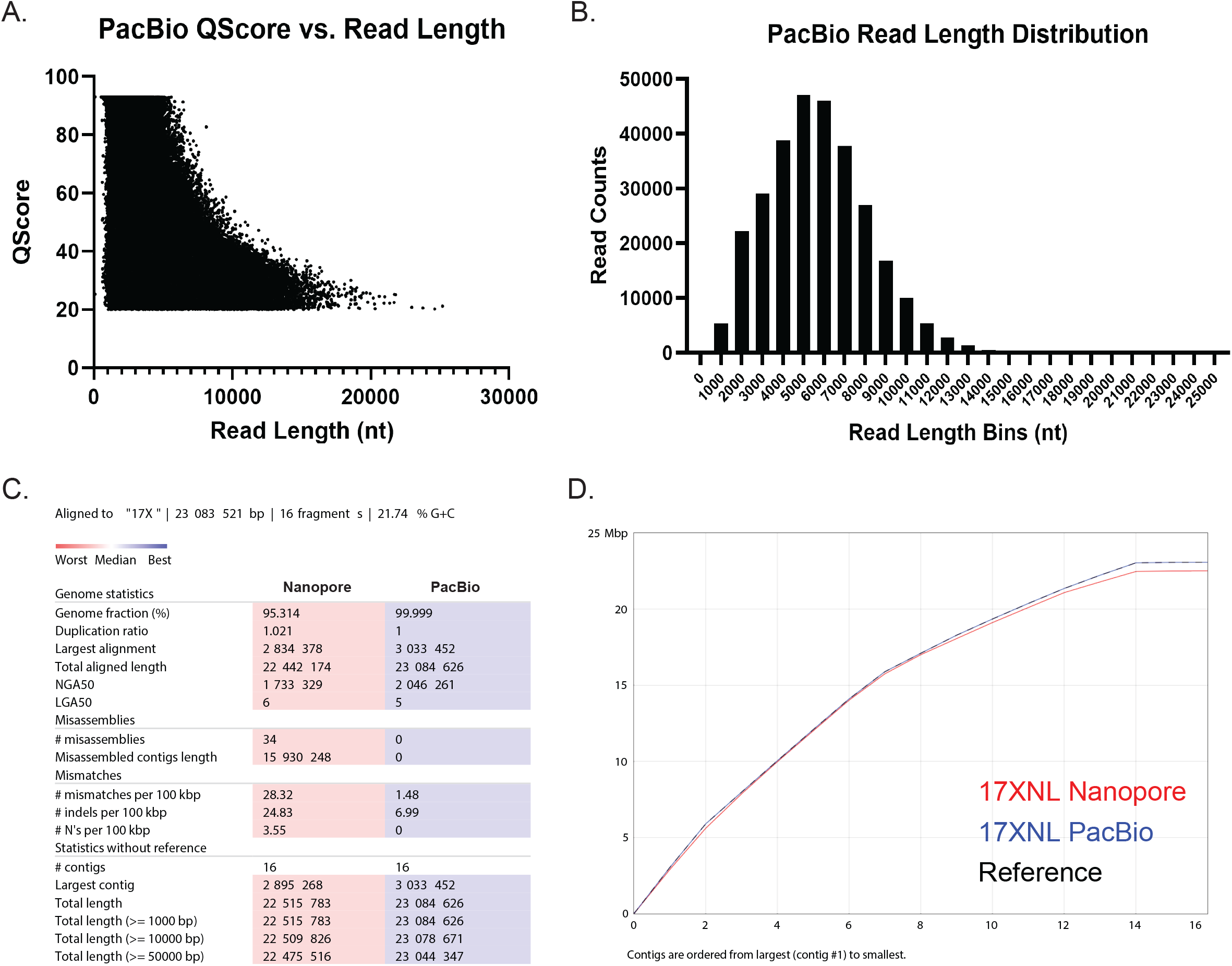
PacBio HIFi high-quality long reads improve upon the pre-existing Py17XNL genome and outperform a hybrid assembly approach with Nanopore and Illumina sequencing. (A) QScore vs. read length distribution for a PacBio sequencing run that was used to construct the final Py17XNL_2 genome assembly is presented. Note: HiFi PacBio sequencing has a minimum QScore threshold of 20, and a maximum QScore threshold of 93. (B) A histogram is plotted to illustrate the distribution of PacBio read lengths. (C) A comparison of assembly statistics between Nanopore and PacBio sequencing runs is provided. All statistics are based on contigs of size >=500 bp. (D) The cumulative length of contigs is plotted from largest to smallest.

We generated genome assemblies from both long-read datasets with the bioinformatic workflows described in Figure 2. To create the Nanopore/Illumina hybrid genome assembly, we assembled the Py17XNL Nanopore data using Flye (30) and scaffolded the contigs using the Py17X genome as a guide in conjunction with the RagTag scaffolding program (30, 31). Finally, we layered error correction onto it in a multi-step approach, first using nextpolish, followed by multiple rounds of consensus generation based on Illumina data alignment and variant calling (Figure 2) (32). Through this process, we were able to reduce the number of contigs down to 16, but at the cost of covering less of the genome (95%) and introducing 34 misassembles (Figure 1C) as defined by the assembly evaluator program Quast (33). The PacBio-based genome assembly was generated with the HiCanu program (34), which produced a *de novo* genome assembly with 132 contigs (Figure 2). The resulting contigs were filtered to contain only the target species by aligning them against the Py17X genome using minimap2 (35). Contigs that had a primary alignment length of >2% of the 17X reference chromosome were assigned the matching chromosome names. A consensus genome was then created by aligning these contigs with the 17X reference genome and filling in the missing genomic regions, mainly chromosomal ends. This resulted in a final assembly of 23.08 Mb with 16 contigs (Figure 1C, Table 1). We have adopted the higher quality PacBio-based genome assembly for the *P. yoelii* 17XNL strain for the rest of our analyses and for provision to the community on PlasmoDB.org, which we term Py17XNL_2 to distinguish it from the original genome assembly (Py17XNL_1) (10, 36). However, as both the hybrid Nanopore/Illumina and PacBio assemblies are potentially valuable to our research community, both assemblies have been publicly deposited in NCBI.

**Figure 2:**
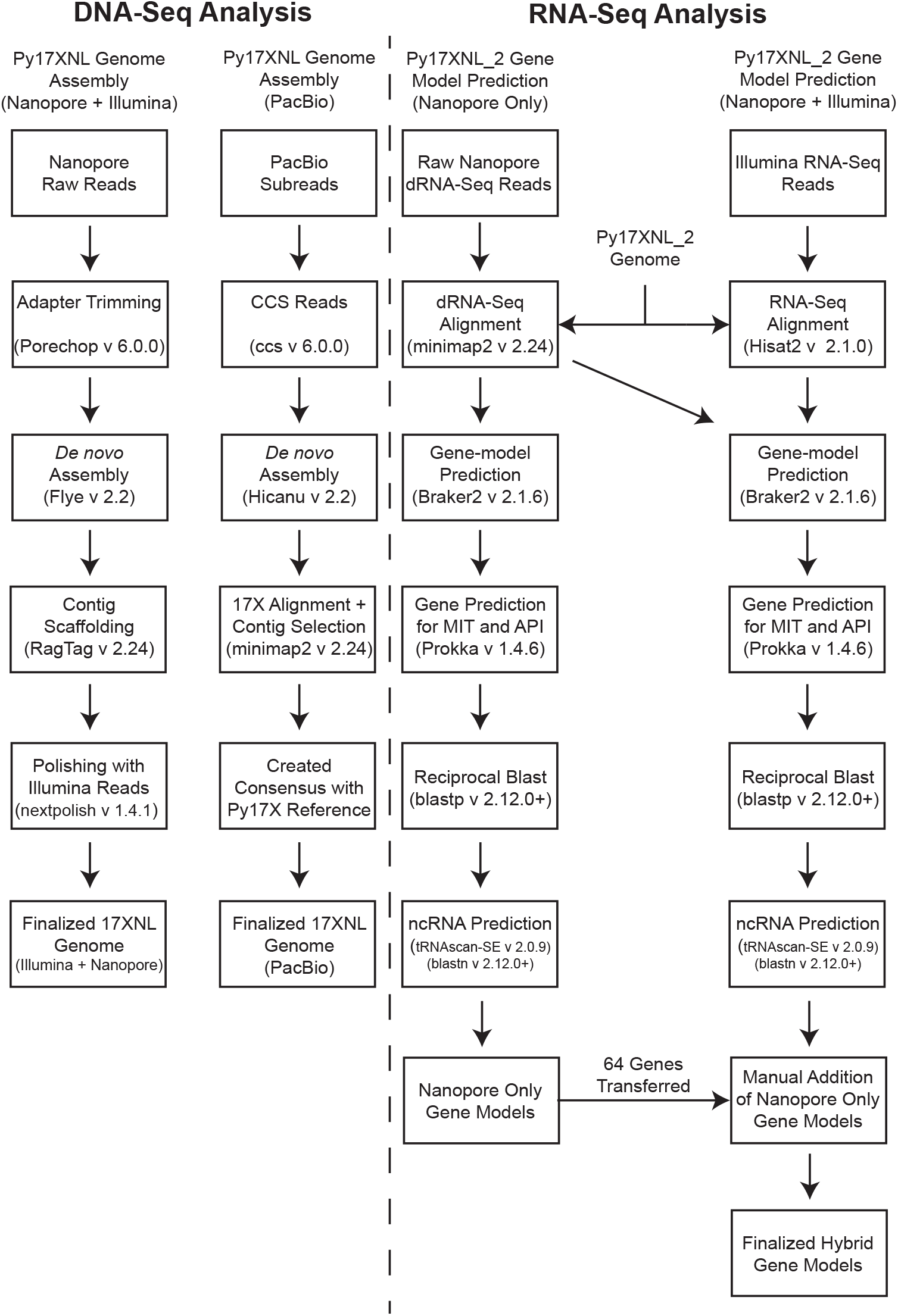
Bioinformatics workflow used for genome assembly and annotation. (Left) Genome Assembly: High-accuracy ccs reads that were generated from PacBio subreads and trimmed Nanopore reads were *de novo* assembled to create draft genomes. Contigs were selected, and chromosome names were assigned based on the *P. yoelii* 17X reference genome alignment. Further processing of the Nanopore + Illumina hybrid assembly involved implementing scaffolding and iterative polishing. (Right) Gene-model prediction: A Nanopore dRNA-seq-based gene model and a hybrid gene model combining both Nanopore dRNA-seq and Illumina RNA-seq data were generated using Braker2. The predicted genes were annotated using reciprocal BLAST against *P. yoelii* 17X proteins. Illumina RNA-seq reads were previously reported (37).

**Table 1.** Summary of finalized genome assembly and gene model creation statistics. * Determined using the Quast program by alignment to the 17X reference for this study. ** Determined using a BUSCO analysis for this study. *** As reported in Carlton et al Nature 2002. N/D: Not determined.

|  | 17X (Otto *et al.* 2014) | 17XNL_1 (Carlton *et al.* 2002) | 17XNL_2 (This Study) |
| --- | --- | --- | --- |
| **Quast summary** |  |  |  |
| Number of contigs | 16 | 5617 (5687<sup>***</sup>) | 16 |
| Assembled genome size | 23.08Mb | 23.1 Mb | 23.08 Mb |
| Genome Fraction (%) | - | 88.68% * | 99.90% |
| Largest Alignment | - | 51.5 Kb | 3,033,452 |
| NGA50 | - | 7,668 * | 2,046,261 |
| LGA50 | - | 851 * | 5 |
| Mismatches per 100 kb | - | 72.72 * | 1.48 |
| Indels per 100 kb | - | 44.07 * | 6.99 |
| N's per 100 kb | - | 50 * | 0 |
| **BUSCO summary** |  |  |  |
| BUSCO genome completeness | 98% ** | 83.9% ** | 97.60% |
| BUSCO protein completeness | 99% ** | 64.7% ** | 90.00% |
| **Annotation summary** |  |  |  |
| Number of genes | 6,263 | 7774 (5878<sup>***</sup>) | 6,086 |
| Number of mRNAs | 6,041 | 7,724 | 7,052 |
| Number of tRNAs | 79 | 50 (39<sup>***</sup>) | 66 |
| Number of rRNAs | 40 | 0 (7<sup>***</sup>) | 40 |
| **Variant summary** |  |  |  |
| Variants with respect to 17X | - | N/D | 1,955 |
| Number of substitutions | - | N/D | 360 |
| Number of insertions | - | N/D | 833 |
| Number of deletions | - | N/D | 762 |
| Number of variants in CDS | - | N/D | 334 |

\* Determined using the Quast program for this study.
\*\* Determined using a BUSCO analysis for this study.
\*\*\* As reported in Carlton *et al* Nature 2002
N/D Not Determined

### Nanopore Direct RNA sequencing provides new information to pre-existing gene models

We also set out to create more comprehensive gene models to increase the utility of the new Py17XNL_2 genome assembly for the P. yoelii 17XNL strain. In the currently available gene models for *P. berghei* (ANKA) and *P. yoelii* (17X, 17XNL) on PlasmoDB, only the coding sequences of genes are provided with no designation of untranslated regions (UTRs), and little information is provided about alternatively spliced transcripts. We generated gene models that provide complete transcript information, including start and stop codons, transcription start and stop sites, and UTRs. Experimentally, we performed Nanopore direct RNA-seq in biological duplicate to generate long sequence reads of asexual and sexual blood stage transcripts. Briefly, total RNA was extracted from parasitized mouse blood to create an RNA-seq library that was sequenced with the Nanopore Direct RNA-Sequencing Kit (Supplemental Figure 1, Supplemental Figure 4, Supplemental Table 1). These direct RNA-sequencing reads were quality controlled using Nanoplot with the same parameters described for Nanopore ligation DNA-sequencing for both “Fast” and “High Accuracy” basecalling approaches (Supplemental Figure 5) (29). In total, 429,888,068 bases were sequenced after combining the replicates, with an average Qscore of 12 after high accuracy basecalling, which again outperformed the fast basecalling approach (Supplemental Figure 5, Supplemental Table 2)). The mean read length across replicates was 858 bases, with the longest read being 8,789 bases (Supplemental Figure 5, Supplemental Table 2).

Gene models were created with two alternative methodologies using Nanopore direct RNA-seq long reads alone or in combination with our previously published Illumina short-read RNA-seq of mixed asexual and sexual blood stage parasites (Figure 2) (37). These parallel approaches are both informative, given the strengths and limitations of both sequencing techniques. Nanopore direct RNA sequencing provides information that allows us to identify long/full-length sequencing reads that initiate at the 3’ end of mRNAs (38). However, when the full-length mRNA is not sequenced, less information is provided for the 5’ end (38). This limitation is remedied by the strong depth and breadth of sequencing coverage provided via Illumina sequencing. For both approaches, Nanopore RNA-seq reads were aligned to our Py17XNL_2 genome using minimap2 (35). For gene models created with both Nanopore and Illumina RNA-seq data, in parallel, the Illumina short reads were aligned to the Py17XNL_2 genome using Hisat2 (31375807). To create the gene models and assign gene names/descriptions, we used Braker2, as well as reciprocal blast searches using the blastp program of the BLAST suite (39, 40). The Nanopore-only approach helped us identify 5,683 genes, 5,828 mRNAs, 66 tRNAs, and 40 rRNAs. Using the Nanopore/Illumina hybrid approach, we found 6,077 genes, 7,047 mRNAs, 66 tRNAs, and 40 rRNAs (Table 1). Gene models that were generated using both Nanopore and Illumina reads more closely matched the anticipated UTR length that was defined in recent *Plasmodium falciparum* transcriptomics data (13). A representative example of this more comprehensive gene model is illustrated in Figure 3A.

**Figure 3:**
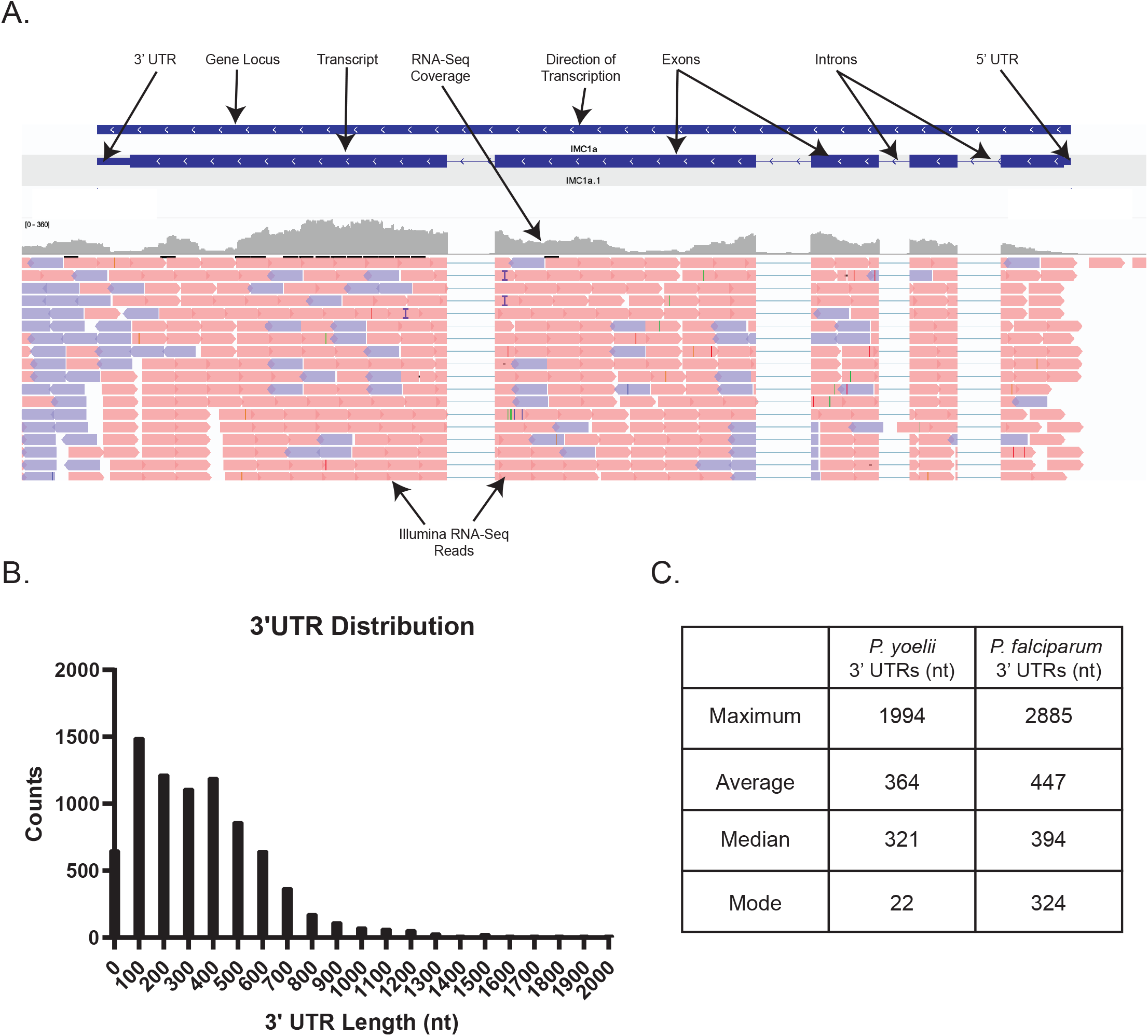
Expanded *Plasmodium yoelii* 17XNL gene models leveraging RNA-seq data. (A) An example gene model depicting IMC1a and its respective sequence features is provided. (B) The 3’UTR length distribution of all detected mRNAs is plotted as a histogram for chromosomal and mitochondrial genes. Transcripts encoded by the apicoplast are not polyadenylated and were not detected by Nanopore dRNA-seq. (C) The maximum, average, median, and mode of the 3’ UTR lengths from all chromosomal and mitochondrial transcripts are compared to those from a *Plasmodium falciparum* dataset (13).

Nanopore direct RNA-seq initiates at the 3’ end due to the use of poly(dT) sequencing primers. As a result, significantly higher coverage was obtained for the 3’ UTRs than for 5’ UTRs (Supplemental Table 3). The higher coverage allows us to further analyze the 3’ UTR length distribution for *P. yoelii* 17XNL, which is of interest as *cis-*regulatory elements are often found in this portion of eukaryotic mRNAs (41). The majority of reads have a UTR length between 100 and 200 bp, with a mean length of 364 bp. The largest UTR reported for the H2B.Z histone variant mRNA, with 1994 nt (Figure 3B, Supplemental Table 3). Compared to the most up-to-date *P. falciparum* transcriptome, which used DAFT-Seq to resolve UTRs, P. yoelii 17XNL’s 3’ UTRs appear slightly shorter on average (Figure 3C) (13).

### Comparison between Py17XNL_2 and reference genomes demonstrates the completeness of the assembly

Using the Py17XNL_2 genome assembly and associated gene models, we compared our results to the original Py17XNL genome (Py17XNL_1) and the Py17X reference genome. As anticipated, there was a substantial reduction in the number of gaps/misassembles and greater genome coverage when comparing Py17XNL_2 vs. Py17XNL_1 (Table 1). Although our Nanopore/Illumina-based genome assembly (Py17XNL Nanopore) contained the same number of contigs as the PacBio-based Py17XNL_2 assembly, the significantly fewer misassemblies generated in the PacBio-based assembly provided a more accurate reference genome for future research uses. Additionally, our final PacBio-based assembly (Py17XNL_2) closely resembles that of Py17X, which was also created using recently developed sequencing technologies (15). However, when compared to Py17X, the Py17XNL_2 assembly has lower coverage of repetitive sequences at the sub-telomeric regions, which precluded us from robustly assembling these regions for some chromosomes, requiring consensus generation based on alignments to the Py17X reference genome. Similarly, the 6,086 new gene annotations more accurately represent the anticipated number of genes for Py17XNL and more closely match those annotated in other *Plasmodium* species (Table 1). Moreover, these new gene models include both coding sequences, UTRs, and transcript isoforms, which are lacking in the provided gene models currently available on PlasmoDB for this specific species. In addition to these assessment metrics, we determined the completeness of the reference genome based on marker genes. To quantify this, we used a Benchmarking Universal Single-Copy Orthologs (BUSCO) analysis that detects whether a predefined set of single-copy marker genes in the *Plasmodium* lineage are present in these data (Figure 4) (42). This BUSCO dataset contains 3642 BUSCO groups from 23 different species, including *P. falciparum* 3D7, *P. yoelii* 17XNL, *P. vivax, P. berghei* ANKA, *P. chabaudi*, and others. From this search, 3556/3642 (97.6%) of complete and single copy BUSCOs were found to be present, indicating that this genome assembly and gene annotation has a high level of completeness (Figure 4).

**Figure 4:**
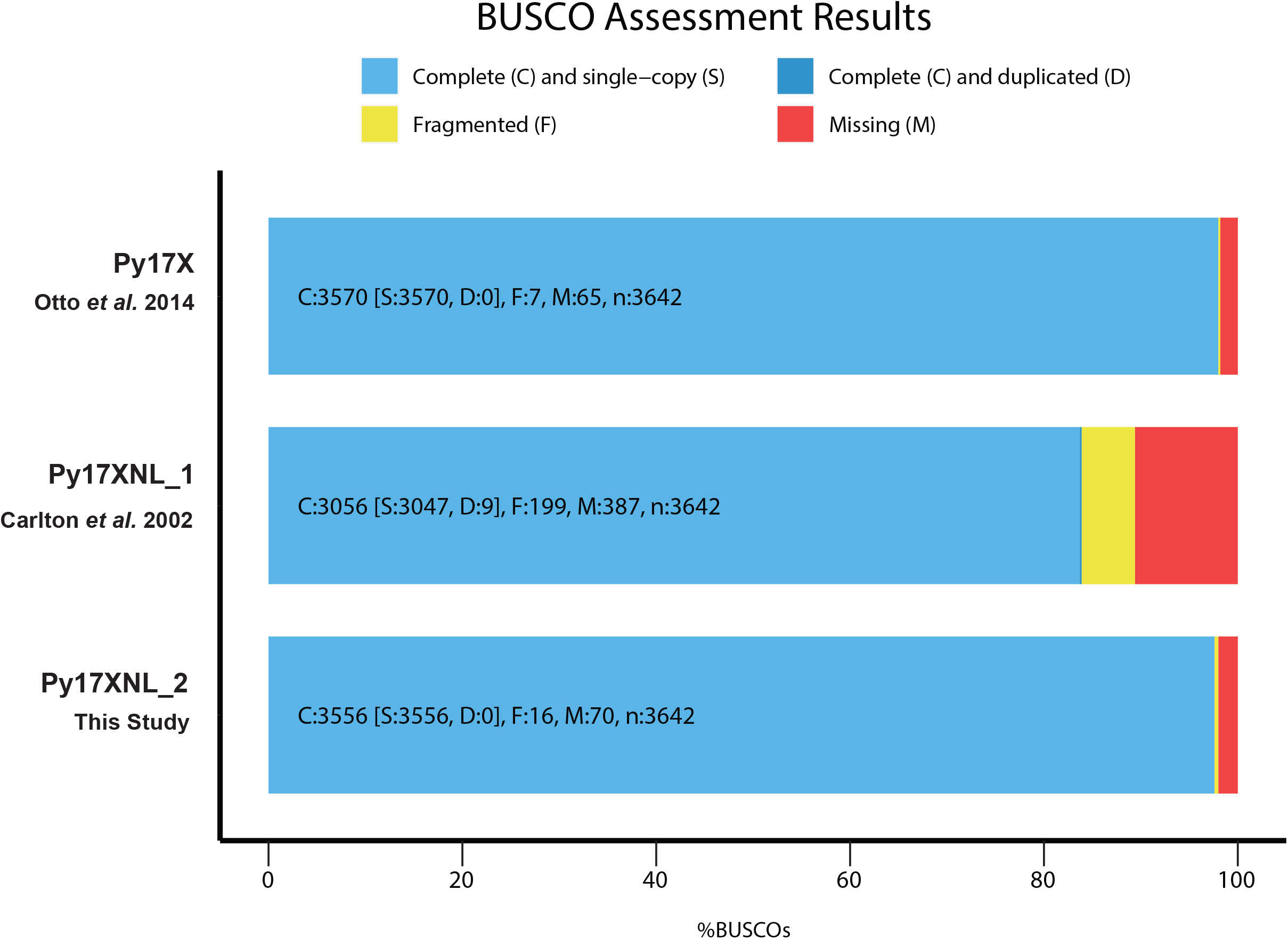
BUSCO analysis demonstrates genome assembly completeness. Of the 3,642 BUSCO groups that were searched, 3,556 single-copy BUSCOs were found to be present in the 17XNL_2 assembly resulting in a completeness score of 97.6%. The BUSCO results for Py17XNL_1 (83.9%) and Py17X (98.0%) reference genomes are also shown for comparison.

### Variation between the Py17XNL and Py17X reference genomes primarily resides in the intergenic regions and the ends of chromosomes

As Py17X is commonly used as an interchangeable proxy genome for Py17XNL, we sought to determine what similarities and differences exist between the strains and how the differences may impact genetic studies. We performed chromosome-wide alignments between the Py17X and Py17XNL_2 genomic builds using the minimap2 (35) program and assessed the genome-wide variants with paftools. We observed extensive linear agreement between the two strains, with 99.9% of the Py17X genome matching with the Py17XNL_2 genomic build (Figure 5A). We did not detect any large structural variation between the strains, a finding also supported through our alternative Py17XNL_2 Nanopore/Illumina genome build. At the same time, we also identified a total of 1,955 potential single nucleotide/short variants across the two strains, the majority (62%) of which were found in intergenic regions (Figure 5B). We found that the apicoplast genome was identical between strains, whereas the Py17XNL_2 mitochondrial genome has a 127 bp deletion in the middle of its sequence. Compared to Py17X, the deletion is located in the intergenic region between *cox1* (PY17X_MIT00800) and a ribosomal RNA fragment annotated as PY17X_MIT00700. Together, we conclude that while these two strains are highly similar, there are sequence differences that may be functionally relevant.

**Figure 5:**
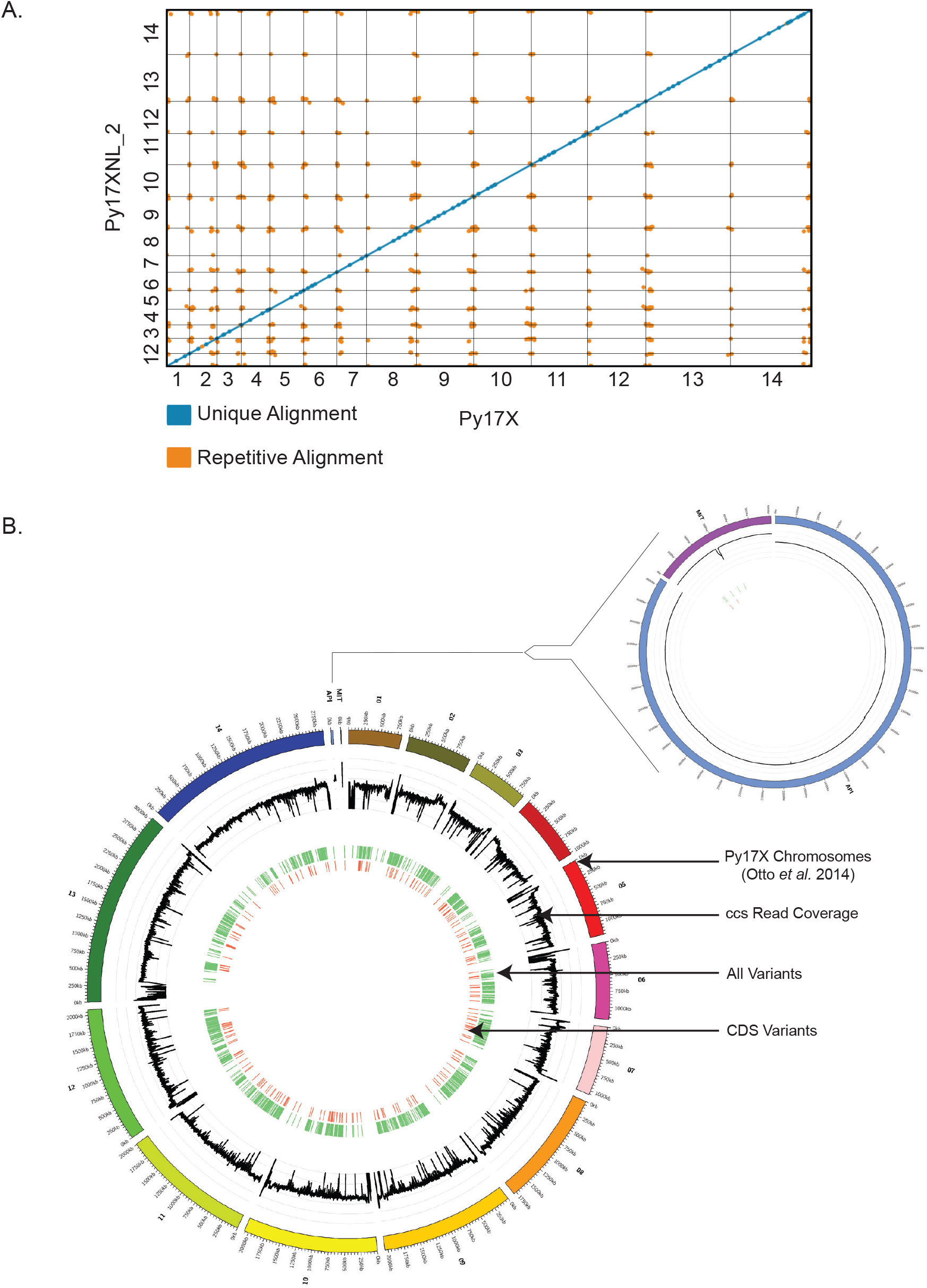
Differences between the *P. yoelii* 17X and 17XNL_2 assemblies. (A) The Py17XNL_2 reference genome was mapped to Py17X to determine their degree of similarity. A dot plot depicting this agreement is shown, with blue lines denoting unique alignments and orange lines depicting repeat regions. (B) A circos plot is presented with the following tracks listed from outside to inside: 1) Py17X reference genome, 2) Py17XNL_2 ccs read coverage in the natural log scale (minimum value of 0 and maximum value of 8), 3) SNPs and indels between the two genomes are shown in light green, 4) SNPs and indels in the coding sequence of genes are shown in orange. An expanded view that includes the apicoplast and mitochondria is shown separately.

To further determine the potential impacts of these genome variants, we characterized the position of variants with respect to nearby genes and, when applicable, determined the specific DNA and amino acid changes that would result from the change. Most variants were found to be located in intergenic regions and were characterized as single base pair indels (Figure 6, Figure 7A,B). Of the 334 variants that fell within coding regions, we characterized the changes in nucleotide and amino acid protein composition of the encoded proteins (representative examples are provided in Figure 6, nucleotide and amino acid level changes are provided in Supplemental Table 4). Due to the over representation of single bp indels, the majority of amino acid changes lead to frameshifts (Figure 7C). Most of these frameshifts took place in genes that were unnamed with an unknown function, requiring further investigation to determine the biological impacts of these differences.

**Figure 6:**
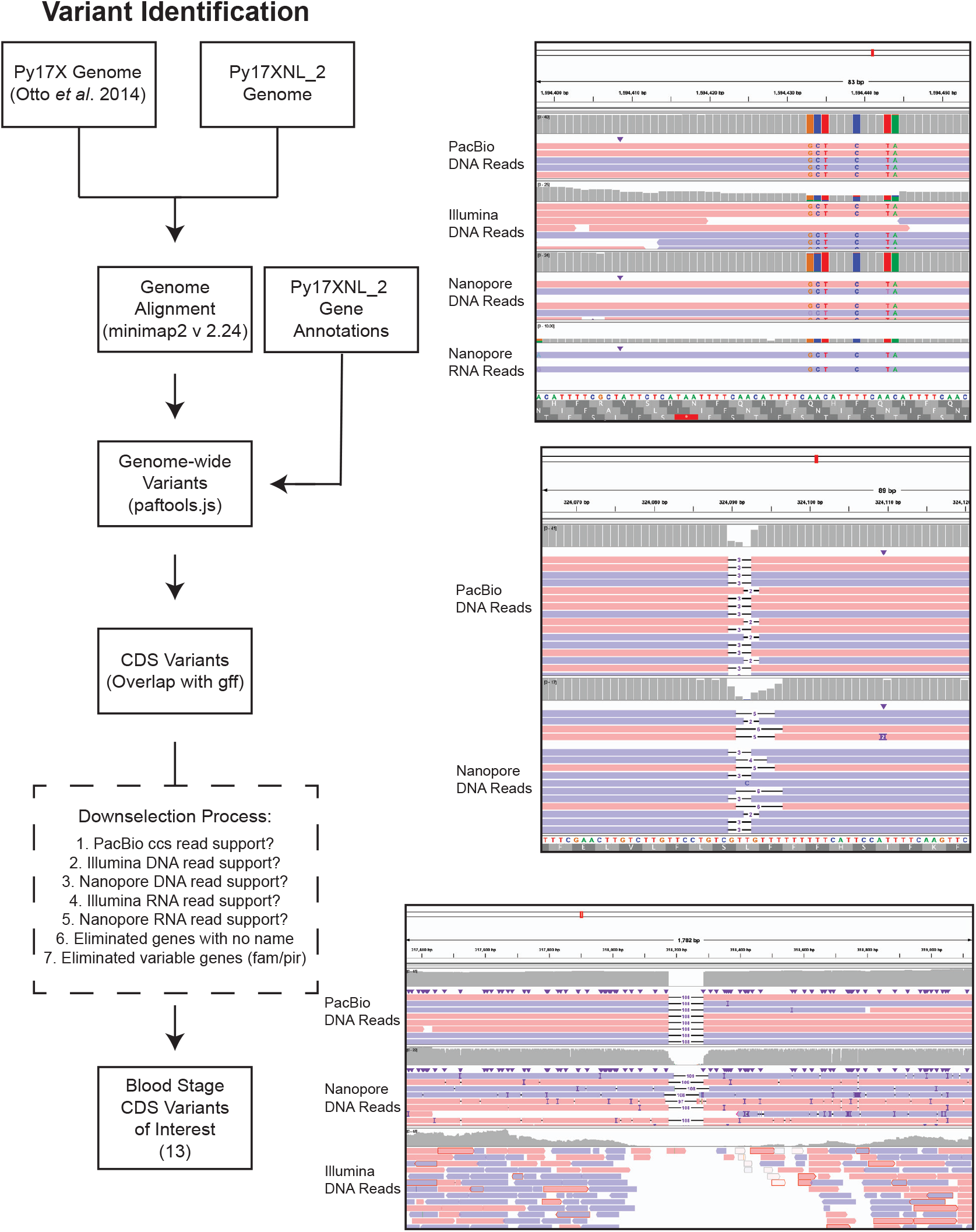
Identification of blood stage-expressed variants between 17X and 17XNL_2. (Left) Variants of interest that are expressed in blood-stage parasites were chosen based on the presence of the variant sequence within the coding sequence, the extent to which the variant calls are supported by sequencing data, and if the gene has been named. Downselected genes are further described in Supplemental Table 4. To be considered, at least two sequencing methods needed to support the variant call, with at least 80% of the reads in agreement and a minimum of five reads at the position (three read minimum for Nanopore). (Right) IGV snapshots with representative examples of different variants found in AP2-SP (PY17XNL_1303202), RAD50 (PY17XNL_0104722), or CSP (PY17XNL_0404050) are presented top to bottom.

**Figure 7:**
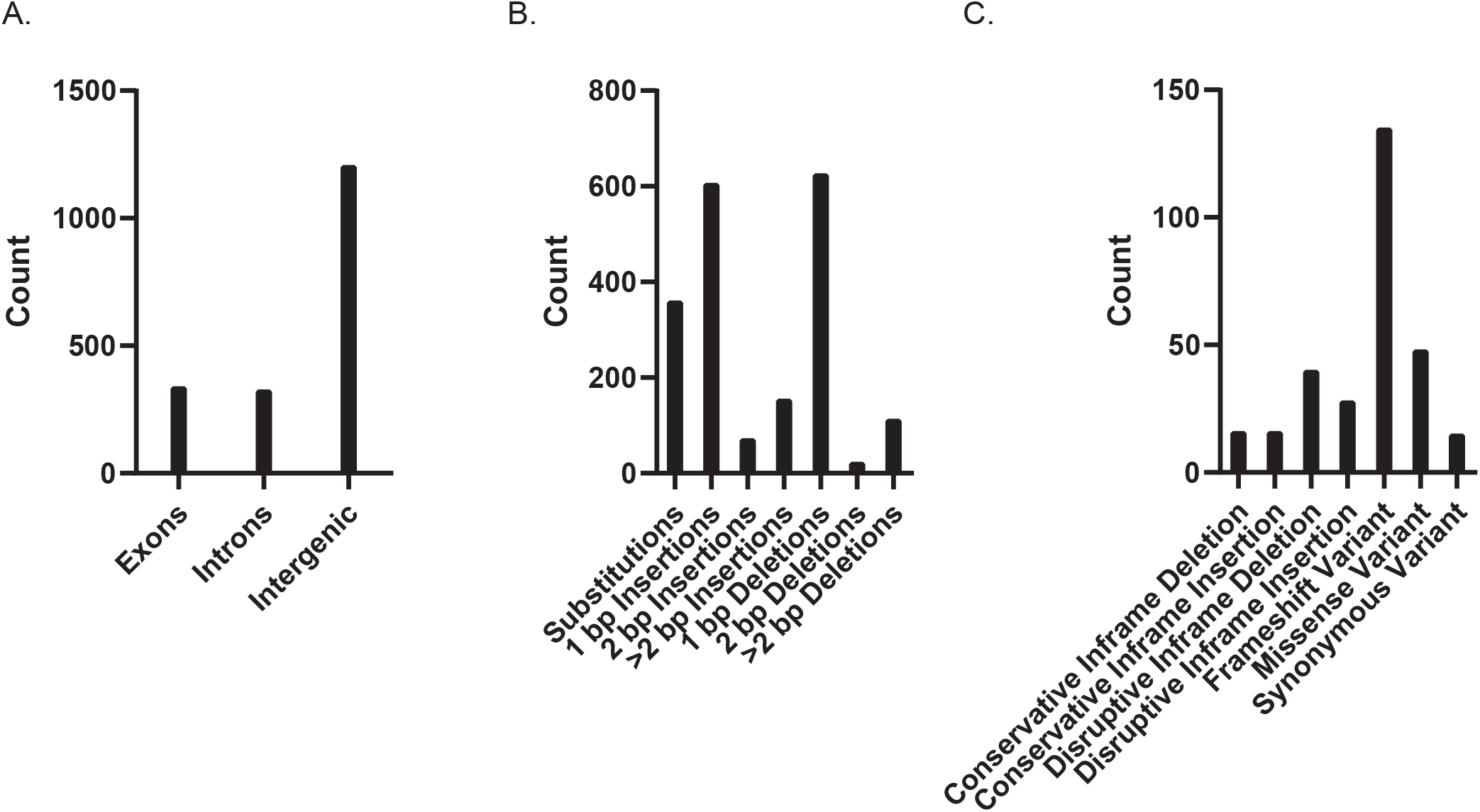
The location and potential impact on translation of variants between 17X and 17XNL_2 genome assemblies. (A) The distribution of variant locations throughout the entire Py17XNL_2 genome is shown. (B) The types of variants represented within the Py17XNL_2 genome with their respective counts are plotted. (C) The distribution of variant types within coding sequences is depicted as a bar graph.

To further interrogate those variants that occurred in well-characterized genes, we manually curated the results and verified the variant calls via various quality measures. We checked if the variant had sufficient PacBio ccs read support (80% of reads support the variant with a minimum of 5x coverage at the region), and when possible, also determined if additional Nanopore/Illumina DNA and RNA sequencing reads supported the variant (80% of reads support the variant with a minimum of 5x coverage for Illumina sequencing and 2x coverage for Nanopore sequencing). Through manual curation, a substantial number of variants had support from at least three sequencing methodologies. Due to the strict thresholds of this variant calling process, some sequencing methods did not capture the variant sufficiently enough to provide support, typically due to a lack of coverage at the position of the variant. An example of this occurring is with CSP, which had a large deletion that was adequately supported by PacBio and Nanopore DNA-seq data (Figure 6). Illumina DNA-seq reads, which should capture this variant due to their high accuracy, instead have a complete loss of coverage, with only one read correctly mapping to this repetitive region (Figure 6, Bottom Panel). As a result, we encourage the use of long-read sequencing platforms to identify variants that may be missed when using Illumina sequencing.

Based upon these criteria, we identified if these genes were expressed in asexual/sexual blood stages due to sequencing support from either Illumina or Nanopore direct RNA-seq and created separate variant lists accordingly (Supplemental Table 4). Finally, we filtered out genes with no annotated gene name and those that belong to a variable gene family (fam/pir gene families) (Figure 6). After this filtering, we focused our analyses on the remaining 13 blood stage-expressed genes (Table 2). Although the biological implications of the differences between Py17X and Py17XNL will need further experimental validation, many variants could have interesting impacts. One such example is *ap2-sp*, which has both synonymous and missense variants between the AT-hook and AP2 domain (43, 44). AP2-SP is an ApiAP2 transcription factor with many target genes that are expressed specifically in the sporozoite stage of the *Plasmodium* life cycle (43, 45, 46). It has also been shown that disruption of this gene results in the loss of sporozoite formation entirely in the related *P. berghei* parasite and has important activities in blood stages in *P. falciparum* (43, 45-47). Another affected gene is *pk4*, which encodes an essential eIF2α kinase related enzyme and contains changes in its non-cytoplasmic domain as determined by InterPro domain predictions across 17X and 17XNL strains (48-51). Further study of these genes and several other candidates is warranted to understand the biological role these variants may play across strains.

**Table 1.**
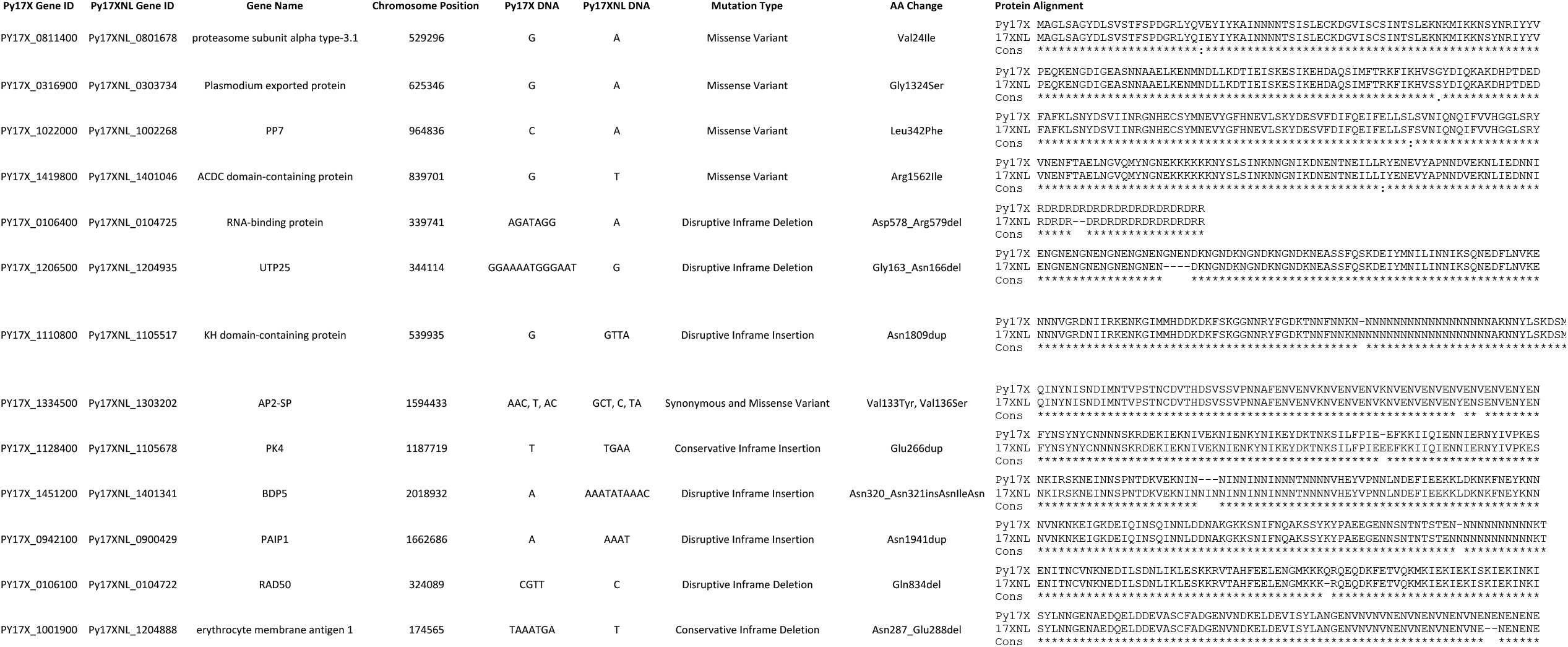
Prioritized list of coding sequence variants between the Py17X and Py17XNL genome.

Among the non-blood stage expressed genes, *trap, lisp2*, and *csp* all had variants in their coding sequences when comparing 17XNL to 17X. Of these, the most notable one is a large in-frame deletion within the repeat region of CSP, leading to the loss of six of the repeating units of D/PQGPGA in Py17XNL (Supplemental Figure 6). Similarly, the YM strain of *P. yoelii* is even shorter and lacks an additional repeating unit compared to 17XNL (Supplemental Figure 6) (15). In *P. berghei*, it was found that 25% of this repeat region could be eliminated before impacting parasite development, which is approximately the length reduction observed in 17XNL and YM as compared to 17X (52). Therefore, this may reflect a minimum repeat length for CSP functions.

## Discussion

Here we have created a high-quality genome assembly with experimentally validated gene models for the commonly used 17XNL strain of the *P. yoelii* malaria parasite species. We envision this will be an important resource to the malaria research community, as it provides a much-needed update to the Py17XNL_1 reference genome, which was among the first to be completed in the early days of the genomics era 20 years ago (10). By directly comparing the strengths and genome assemblies created from either PacBio HiFi sequencing or a combination of Nanopore DNA-seq and Illumina DNA-seq, we identified that while the hybrid Nanopore/Illumina approach yielded a robust genome assembly, the PacBio HiFi-based assembly consisted of fewer misassembles and covered a greater fraction of the genome. Therefore, we have chosen the PacBio-based genome assembly as our new working reference genome for *P. yoelii* 17XNL strain, which we have designated as Py17XNL_2 within this study. Our findings align with many recent studies conducted to improve the reference information on *Plasmodium* species, which also utilized an exclusively PacBio-based approach (12, 14, 17, 53). We also deemed it important to conduct both approaches to leverage the strengths of Nanopore sequencing, which permit greater detection of large-scale structural variants in the genome as compared to approaches with shorter read lengths (26). The strengths of Nanopore sequencing have also been leveraged by other sequencing efforts, most notably the recent telomere-to-telomere sequencing effort of the human genome that used ultra-long read approaches (54). During this study, advances in Nanopore basecalling software were made that enabled more accurate sequencing without the need for re-sequencing or new hardware. We therefore directly compared the previous “fast” vs. new “high accuracy” basecalling algorithms and observed a substantial increase in Qscores associated with the same DNA and RNA sequencing data (Supplemental Table 2). However, even with the use of high accuracy basecalling, PacBio data still enabled the most accurate Py17XNL genome assembly and covered the greatest fraction of the genome.

To provide an even more useful genome reference, here we also provide new gene annotations for the 17XNL strain of *P. yoelii* to facilitate more reliable forward and reverse genetic studies of this key model malaria species. Regardless of the rodent malaria RNA-seq studies that have been performed, gene models available on PlasmoDB for *P. berghei* and *P. yoelii* only consist of their putative coding sequences. Here we have now added experimentally validated information on alternatively spliced transcripts and untranslated regions (UTRs) of Py17XNL blood stage-expressed genes. To date, the only other comparable efforts in our field have been applied to *P. falciparum* with a focus on either identifying alternatively spliced transcripts or experimentally defining and annotating long noncoding RNAs (lncRNAs) (55-57). Additionally, because Nanopore direct RNA-seq reads initiate at the 3’ end of mRNAs and progress toward the 5’ end, it is also strong-suited in providing information about the 3’ UTRs of a population of mRNAs. From this, we created both a Nanopore-only and a Nanopore/Illumina hybrid gene model annotation that can both be useful to researchers depending on the questions they are pursuing. We are therefore providing both gene model files as resources to our community. These gene models include well-defined 3’UTRs for Py17XNL that are in agreement with the length distribution of those described for *P. falciparum* (13). Due to the strengths of this approach, we anticipate that Nanopore direct RNA sequencing will become a useful tool for future work on *Plasmodium* parasites, especially as sequencing chemistry and basecalling algorithms improve.

With this greatly improved Py17XNL reference genome, we also were able to critically analyze genomic variation across the 17X and 17XNL strains of *P. yoelii*. As it is currently common practice to use Py17X as a proxy genome for Py17XNL for genomic studies, we thought it was important to begin addressing whether biologically relevant differences were present that would impact such efforts. By aligning the two genomes, we saw that there was an excellent linear agreement between them, with most variation taking place in intergenic regions. In total, there were 1,955 variants across the entire sequence, with 334 of those being in the coding sequence of genes. Most of these variants were single bp indels that most likely accounted for the overrepresentation of frameshift variants in the respective amino acid sequence. Upon further analysis of these variants, some interesting questions arose regarding the biological implications that these changes could have. Specific examples of genes with impactful variants include the ApiAP2 transcription factor AP2-SP and PK4, which are essential for *Plasmodium* development and warrant follow-up studies (43, 45, 46, 48, 50, 51). Aside from these blood stage-expressed genes, it is also important to note the large-scale differences between the 17X, 17XNL, and YM strains of *P. yoelii* in the central repeat of CSP, which are 150, 114, and 108 amino acids long, respectively (Supplemental Figure 6). The in-frame deletions result in truncations of entire six amino acid repeating units of D/PQGPGA, with 17XNL having six fewer units and YM having seven fewer than 17X. In *P. berghei*, it was found that 25% of this repeat region could be eliminated before impacting parasite development, which reflects the approximate reduction in repeat length in 17XNL and YM strains as compared to 17X (52). We anticipate that this may reflect a minimum repeat length that is applicable to both highly related species. This is indirectly corroborated by the absence of any reports that have documented significant differences in sporozoite development, functions, or transmissibility between the 17X and 17XNL strains. Additionally, this particular variant was identified in both Nanopore and PacBio long-read DNA-sequencing datasets, with Illumina short-read sequencing lacking coverage at this site to accurately identify this deletion (Figure 6). This highlights the utility of long-read sequencing technologies to resolve highly repetitive genomes.

This *P. yoelii* 17XNL_2 reference genome and its more comprehensive gene annotations provide a resource that we believe will be helpful to the rodent malaria research community. We stress that while most genes are identical between the 17X and 17XNL strains, there is appreciable genomic variance in some important genes that should be considered when conducting genomic studies. Therefore, we conclude that for many efforts, 17X is a suitable genomic proxy for 17XNL, but caution against it for genes where variance exists, such as *csp, trap, lisp2, ap2-sp, pk4*, and others. Given the improvements for both the Py17XNL_2 genome assembly and gene models presented here, we would instead encourage their adoption as the working reference genome and gene annotation source for studies of *P. yoelii* 17XNL.

## Materials and Methods

### Animal Experiments Statement

All animal care strictly followed the Association for Assessment and Accreditation of Laboratory Animal Care (AAALAC) guidelines and was approved by the Pennsylvania State University Institutional Animal Care and Use Committee (IACUC# PRAMS201342678). All procedures involving vertebrate animals were conducted in strict accordance with the recommendations in the Guide for Care and Use of Laboratory Animals of the National Institutes of Health with approved Office for Laboratory Animal Welfare (OLAW) assurance.

### Experimental Animals

Six-to-eight-week-old female swiss webster mice from Envigo were used for all experiments in this work.

### Parasite Preparation and Isolation

Mice infected with wild-type Py17XNL Clone 1.1 parasites from BEI Resources until a parasitemia between 1-3% was reached. Approximately 1 mL of blood was collected from each euthanized mouse, which was then added to 5 mL of heparinized (200 U) 1X PBS to prevent coagulation. The infected blood was spun and the serum was aspirated to isolate the red blood cells (RBCs). Cells were resuspended in 10 mL 1X PBS and then passed through a cellulose column (Sigma #C6288) to remove mouse leukocytes. The RBCs were then lysed in 0.1% w/v saponin in 1X PBS for 5 minutes at room temperature, and parasite pellets were subsequently washed in 10 mL 1X PBS.

### gDNA Preparation

All gDNA samples used for Nanopore, Illumina, and PacBio sequencing were prepared using the NEB Monarch HMW DNA Extraction Kit for Cells and Blood (NEB #T3050) using the manufacturer’s protocol for fresh blood with slight modifications as we have previously described (28). Briefly, the saponin lysed parasite pellet was resuspended in 150 µL of Nuclei Prep Buffer containing RNase A. After resuspension of the pellet, 150 µL of Nuclei Lysis Buffer containing Proteinase K was added and mixed by inversion. The sample was then placed in a thermal mixer at 56°C with an agitation speed of 1500 rpm for 10 minutes. Next, 75 µL of precipitation enhancer was added and mixed by inversion. Two DNA capture beads were added to the tube, along with 275 µL of isopropanol. The sample was then mixed 30 times with manual, slow, end-over-end inversions to ensure the gDNA stuck to the capture beads. The supernatant was removed, and the beads were washed twice with 500 µL of gDNA wash buffer.

Subsequently, 100 µL of elution buffer II was added and the sample was incubated for five minutes at 56°C in a thermal mixer with agitation at 300 rpm. The beads were added to a bead retainer in an Eppendorf tube, and the sample was spun down for 30 seconds at 12,000 *xg*. All samples were stored at 4°C to minimize shearing from freeze-thaw cycles. Fresh gDNA samples were made for replicate 1 and replicate 2 for Nanopore sequencing and the sole sample for PacBio. The same gDNA samples used for Nanopore replicates 1 and 2 were used for Illumina DNA sequencing replicates 1 and 2. Sample concentration and purity were assessed via Qubit and Nanodrop, respectively (Thermo Fisher Scientific® Nanodrop® 2000 and Qubit® instruments with the Qubit dsDNA BR Assay Kit (Cat #Q32853)). Fragment length was assessed using an Agilent Technologies® TapeStation® 4200 system with Genomic DNA ScreenTapes (Cat #5067-5366 and 5067-5365).

### RNA Preparation

RNA samples were prepared from two biological replicates for Nanopore direct RNA sequencing. RNA samples were produced using the Qiagen RNeasy kit using the manufacturer’s protocol with slight modifications to improve yield (Cat # 74104). Briefly, 350 µL of Buffer RLT was added to resuspend the parasite pellet. The sample was passed through a 20-gauge needle five times and put back into the same microfuge tube. Next, 350 µL of 70% ethanol was then added and mixed by pipetting using wide-bore pipettes. The sample was then added to the spin column and was centrifuged for 15 seconds at 8,000 *xg*. The column was washed twice with 500 µL RPE buffer and was again centrifuged for 15-60 seconds at 8,000 *xg*. Residual ethanol was removed by a final spin at these parameters. RNA was eluted from the column into a fresh microfuge tube with 30 µL of DEPC-treated water. The sample was incubated for 15 minutes at room temperature to improve recovery yield. The sample was then collected by centrifugation for 1 minute at 8,000 *xg*. A second elution with 30 µL DEPC-treated water was performed as above to improve yield. To eliminate contaminating DNA, a Dnase I digestion was performed with slight modifications to the Sigma #AMPD1 technical bulletin. Briefly, 8 µL of the prepared RNA was mixed with 1 µL 10X Reaction buffer and 1 µL Dnase I, Amplification Grade, 1 unit/µL (Cat # AMPD1-KT). The sample was gently mixed and incubated at room temperature for 30 minutes. The digestion was terminated by the addition of 1 µL stop solution, followed by heat inactivation of the Dnase I. RNA was precipitated with ethanol by adding 0.1 volume of 3M sodium acetate pH5.5@RT, four volumes of reagent grade 200 proof ethanol, and 0.5 µL 20mg/ml glycogen. The solution was allowed to precipitate overnight at -80°C. The solution was then spun down at 4°C at 12,000 *xg* for 10 minutes. The supernatant was aspirated and 1 mL of 70% ethanol was added to wash the pellet. The pellet was spun down as above and the supernatant was aspirated. The pellet was then allowed to air dry with the tube inverted on a Kimwipe for 10 minutes. Sample concentration and purity were assessed via Qubit and Nanodrop, respectively (Thermo Fisher Scientific® Nanodrop® 2000 and Qubit® instruments with the Qubit dsDNA BR Assay Kit (Cat #Q32853)). RNA integrity was tested using an Agilent 2100 Bioanalyzer.

### Nanopore Ligation-based DNA Sequencing

DNA sequencing of Nanopore replicates 1 and 2 was performed using the SQK-LSK110 Ligation sequencing kit using the manufacturer’s protocol. Genomic DNA (∼1 µg as measured by Qubit) was sequenced on an R9.4.1 (Cat # FLO-MIN106D) flow cell for 24 hours, washing between samples as per manufacturer’s recommendations (EXP-WSH003).

### Nanopore Direct RNA Sequencing

RNA sequencing of Nanopore replicates 1 and 2 was performed using the SQK-RNA0002 Direct RNA-sequencing kit from Oxford Nanopore Technologies using 500ng RNA. All sequencing was performed on an R9.4.1 flow cell for 24 hours, washing between samples as per manufacturer’s recommendations (EXP-WSH004).

### Illumina DNA Sequencing

Illumina DNA sequencing libraries were created using the Illumina DNA PCR-Free Kit with 100 ng of total input (Cat # 20041794). Illumina libraries were sequenced on a NextSeq 550 Mid Output 150×150 paired-end sequencing run.

### PacBio Sequencing

PacBio libraries were created using the PacBio SMRTbell Express Template Prep kit 2.0 (Cat # TPK 2.0) using an input of 2 µg gDNA that was sheared with the Covaris g-TUBE to an average fragment length of 10kb (Cat # 520079). The library was sequenced on a PacBio Sequel using a SMRT Cell 1M v3 LR at a 10 pM library loading concentration with a 2-hour pre-extension time and a 20-hour movie time (Cat # 101-531-000).

### Data Analysis

High-quality HiFi reads were extracted from PacBio sequencing data requiring a minimum of 3 full passes in CCS command (v6.0.0) (31562484). The HiFi reads were *de novo* assembled using HiCanu, specifying a genome size of 23 Mb. The resulting 132 contigs were aligned to the Py17X reference genome using minimap2 (35), and all small contigs that had <2% alignment with a 17X chromosome were filtered out. This reduced the contig number to 30 with a 22.2 Mb genome size. Chromosome names were assigned based on alignment to Py17X. A consensus genome was generated based on the alignment of these 30 contigs to the 17X reference genome using a custom program (Supp File 1). This resulted in a final assembly of 16 contigs totaling 23.08 Mb. The apicoplast was circularized using Circlator (v.1.5.5) (58), followed by manual correction of coordinates based on Py17X alignment.

The second assembly approach utilized both Nanopore and Illumina DNA-seq reads. All Nanopore-based raw reads were first analyzed using Nanoplot (v1.33.0) (29) for quality control purposes. Nanopore reads were assembled using Flye (v2.9) (30) software which resulted in 26 contigs. The assembled contigs were scaffolded with Ragtag.py using the Py17X genome as a reference, which resulted in 17 contigs (31). The resulting assembly was polished in a multi-step approach. First, the assembly was polished using nextPolish (v1.3.1) (32). The homopolymer and long indel errors that were still present were corrected in the second step. For these, 150 base pair reads were simulated from the Py17X genome and mapped to the polished assembly in step 1 using bwa mem (59). Variants were called with freebayes (v 1.2.0) (60), and a consensus was created with bcftools (v 1.15) (61), resulting in a second round of polished assembly. To further correct errors, we mapped the Py17XNL Illumina genomic DNA to the resulting assembly, called variants, and generated a consensus. This resulted in a third-round polished assembly. Overlapping contigs were merged, and the apicoplast sequences were circularized, resulting in a genome assembly consisting of 14 nuclear chromosomes and two organellar chromosomes. The low complexity regions and tandem repeats in both assemblies were soft masked using the tantan program (62). Assembly reports were done to compare Nanopore-based and PacBio-based genomes using the Quast program. The variants between Py17X and Py17XNL genomes were obtained from the minimap2 (v2.18) (35) whole genome alignment using paftools.js. The variants were annotated, and variant effects were obtained using SnpEff (v.5.1d) (63). The assembly completeness was assessed using BUSCO (42). For this assessment, the *Plasmodium* lineage database (plasmodium_odb10), which contained 3642 sequences from 23 *Plasmodium* species, was searched to check for the presence and completeness of the single-copy marker genes.

To create gene models and assign gene names, Braker2 (v2.1.6) (39) was first used to predict genes, which was followed with the use of reciprocal blastp. Two sets of gene models were generated for this assembly. For the first set, Nanopore dRNA-Seq reads were mapped to the assembled genome with minimap2 (v2.18) (35). dRNA-Seq read alignments provided additional exon-intron evidence in Braker2-based gene model predictions. Gene names were assigned by a reciprocal blast of the predicted proteins against Py17X proteins. For the second set of gene-model predictions, both Nanopore dRNA-Seq and Illumina RNA-Seq datasets were used. Illumina RNA-seq reads were mapped to the assembled genome using Hisat2 (v.2.2.1) (64) and were merged with Nanopore dRNA-seq alignments, which were then used for Braker2 gene-model prediction. Additionally, Prokka (v 1.14.6) was used to make gene predictions in mitochondria and apicoplast. Finally, tRNAs were predicted using tRNASCAN-SE (v.2.0.9.) (65), and rRNAs were identified by a blast search of the assembled genome using Py17X rRNAs.

### Data Availability

Datasets associated with this study are available using the following identifiers: SRA BioProject: PRJNA769959, Nanopore assembly accessions: CP086268-CP086283, PacBio assembly accessions: CP115525-CP115540. All assembly files produced in this study are provided as Supplementary File 2, and will be provided to VEuPathDB/PlasmoDB for integration and community use.

## Acknowledgments

We acknowledge the Penn State Genomics Core Facility - University Park for sequencing library preparations and conducting the PacBio and Illumina sequencing described in this study. We thank Tanya Renner and her laboratory at Penn State for critical discussions on Nanopore sequencing. We thank New England Biolabs for early access to high molecular weight gDNA purification kits and their insights on the optimization of this process. We also thank members of the VEuPathDB and PlasmoDB.org teams for assistance with current data files and assemblies, Akhil Vaidya (Drexel University) for discussions about the *Plasmodium* mitochondrion, Photini Sinnis on discussions about CSP and its central repeat sequence, as well as the members of the Llinás and Lindner laboratories for critical discussions of this work.

## Figure Legends

**Supplementary Figure 1: Experimental workflow for all sequencing runs performed**. Four different sample/sequencing types were generated. For each, mice were infected with Py17XNL strain parasites until parasitemia reached 1-3%, at which point blood was collected, passed through a cellulose column, and saponin lysed prior to DNA or RNA recovery. For Illumina, PacBio, and Nanopore DNA samples, the NEB Monarch High Molecular Weight Blood Kit was used. For Nanopore RNA samples, a Qiagen RNeasy Kit with subsequent DNaseI treatment was used. For quality control purposes, a Nanodrop and Qubit were used to assess each biological sample. Additionally, TapeStation and Bionalyzer were used for DNA and RNA samples, respectively. The library preparation methods and sequencing devices used for each sample are also indicated. The Illumina RNA-seq data utilized in this study was previously published by our laboratories and was retrieved from the GEO depository (Accession #GSE136674) (37).

**Supplementary Figure 2: Determination of gDNA fragment length by TapeStation**. (A) The Qiagen Blood Amp Kit or the NEB Monarch High Molecular Weight Blood Kit were used to prepare gDNA and samples were run in parallel on an Agilent TapeStation 4150. High molecular weight gDNA from lane C1 was used for Nanopore replicate one. (B) High molecular weight gDNA used for Nanopore replicate two was run separately on the same Agilent TapeStation 4150 instrument. The hazard symbol in lane B1 indicates the sample was run outside of the manufacturer’s recommended concentration. (C) High molecular weight gDNA that was used for PacBio HiFi sequencing is shown in lane C2 on the right. All other lanes were samples from unrelated experiments.

**Supplementary Figure 3: A comparison of fast basecalling and high accuracy basecalling for Nanopore DNA sequencing**. (A and B) Nanopore ligation sequencing reads from replicate one were basecalled using the fast basecalling algorithm (A) or the high accuracy basecalling algorithm (B). The Qscore vs. read length distribution is depicted as a scatter plot (top), and the read length and their respective counts are plotted as a histogram (bottom). (C and D) The same comparisons as described in A and B were applied to replicate two.

**Supplementary Figure 4: Bioanalyzer results demonstrate that RNA samples are of high quality**. (A) Total RNA isolated for replicate one of Nanopore direct RNA sequencing was run on a Bioanalyzer for quality control purposes. The yellow hazard sign in lane B1 indicates that the markers ran outside of their standard position, leading to an edited RIN. The sample was also run at a 1:5 dilution. (B) The RNA sample used for replicate 2 of Nanopore direct RNA sequencing was run separately in the same way.

**Supplementary Figure 5: A comparison of fast basecalling and high accuracy basecalling for Nanopore direct RNA sequencing**. Nanopore direct RNA sequencing reads from replicate one (A and B) or two (C and D) were basecalled using the fast basecalling algorithm or the high accuracy basecalling algorithm. The Qscore vs. read length distribution is depicted as a scatter plot (top), and the read length and their respective counts are plotted as a histogram (bottom).

**Supplementary Figure 6: The CSP central repeat region length varies across sequenced *P. yoelii* strains**. The amino acid sequence for the central repeat region of circumsporozoite protein (CSP) is shown for *P. yoelii* 17X, *P. yoelii* 17XNL, and *P. yoelii* YM.

**Supplementary Table 1: Quality control measurements from Nanodrop, Qubit, TapeStation, and Bioanalyzer**.

**Supplementary Table 2: Nanopore sequencing statistics for replicates 1 and 2 with the fast basecaller (LA) and the high accuracy (HA) basecaller**.

**Supplementary Table 3: 5’ and 3’ untranslated region (UTR) information by gene name using Nanopore and Illumina (“hybrid”) or Nanopore-only approaches**.

**Supplementary Table 4: Complete list of identified CDS variants with variant sequence and sequencing support information**.

**Supplementary File 1: Makefile that details the bioinformatics workflow used in this study**.

**Supplementary File 2: All assembly files generated in this study, including genome fasta, transcript fasta, cds fasta, protein fasta, and GFF3 files**.

