## Supplemental Figures 1-6 for "Long-Read Genome Assembly and Gene Model Annotations for the Rodent Malaria Parasite *Plasmodium yoelii* 17XNL"

Godin and Sebastian *et al*. Supplementary Figure 1

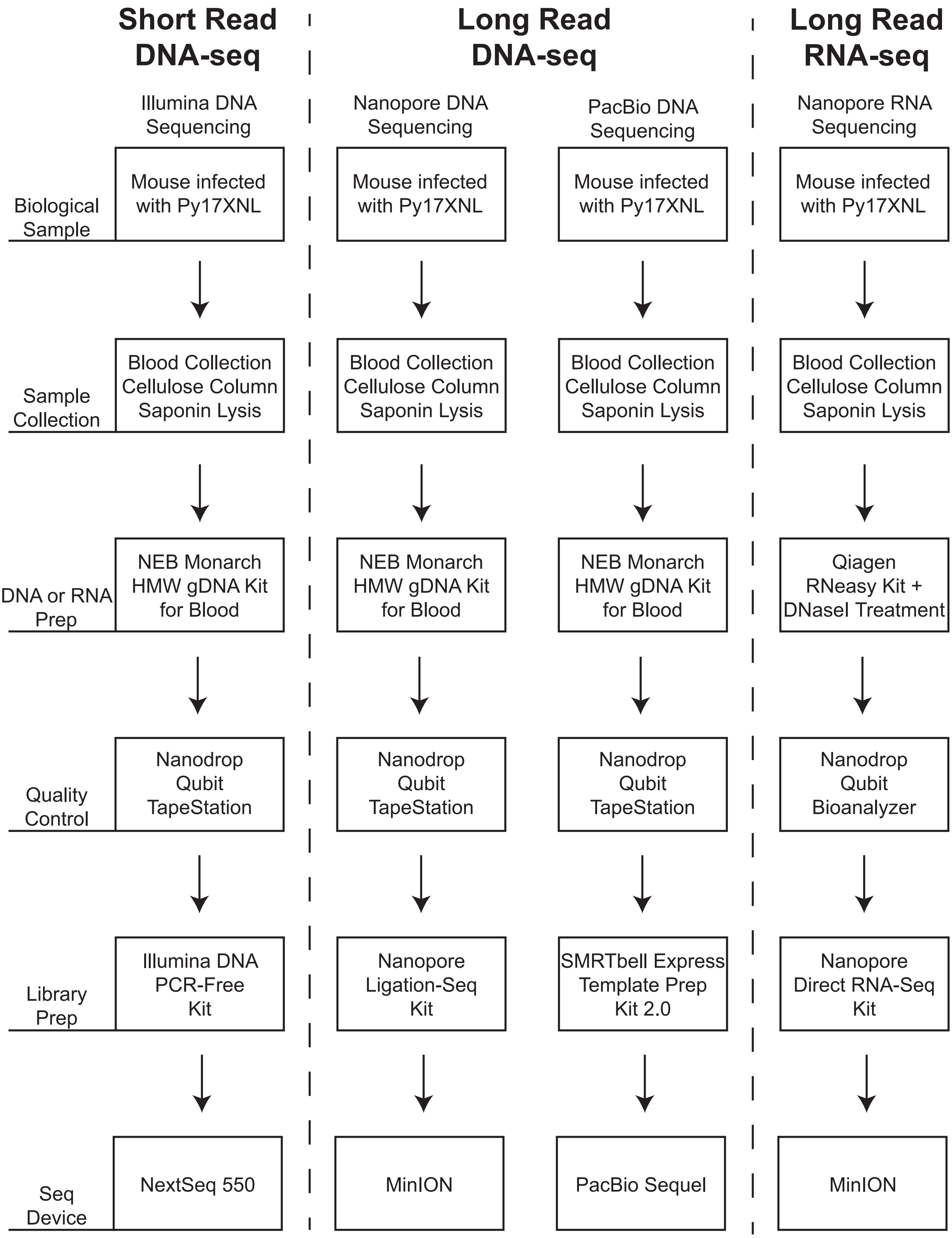


**Supplementary Figure 1: Experimental workflow for all sequencing runs performed.** Four different sample/sequencing types were generated. For each, mice were infected with Py17XNL strain parasites until parasitemia reached 1-3%, at which point blood was collected, passed through a cellulose column, and saponin lysed prior to DNA or RNA recovery. For Illumina, PacBio, and Nanopore DNA samples, the NEB Monarch High Molecular Weight Blood Kit was used. For Nanopore RNA samples, a Qiagen RNeasy Kit with subsequent DNaseI treatment was used. For quality control purposes, a Nanodrop and Qubit were used to assess each biological sample. Additionally, TapeStation and Bionalyzer were used for DNA and RNA samples, respectively. The library preparation methods and sequencing devices used for each sample are also indicated. The Illumina RNA-seq data utilized in this study was previously published by our laboratories and was retrieved from the GEO depository (Accession #GSE136674) (37).

**Godin and Sebastian *et al*. Supplementary Figure 2**

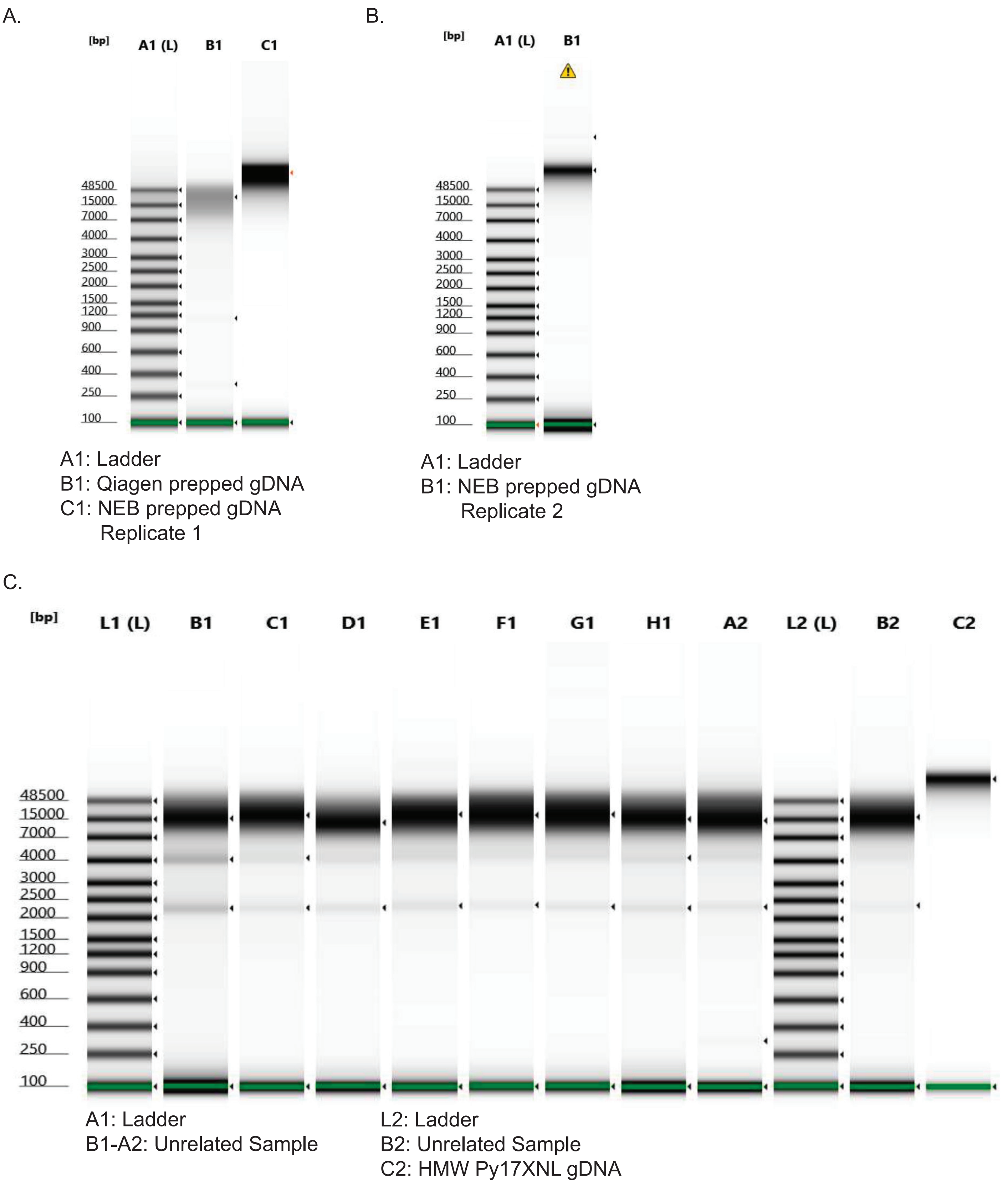


**Supplementary Figure 2: Determination of gDNA fragment length by TapeStation.** (A) The Qiagen Blood Amp Kit or the NEB Monarch High Molecular Weight Blood Kit were used to prepare gDNA and samples were run in parallel on an Agilent TapeStation 4150. High molecular weight gDNA from lane C1 was used for Nanopore replicate one. (B) High molecular weight gDNA used for Nanopore replicate two was run separately on the same Agilent TapeStation 4150 instrument. The hazard symbol in lane B1 indicates the sample was run outside of the manufacturer's recommended concentration. (C) High molecular weight gDNA that was used for PacBio HiFi sequencing is shown in lane C2 on the right. All other lanes were samples from unrelated experiments.

**Godin and Sebastian *et al*. Supplementary Figure 3**

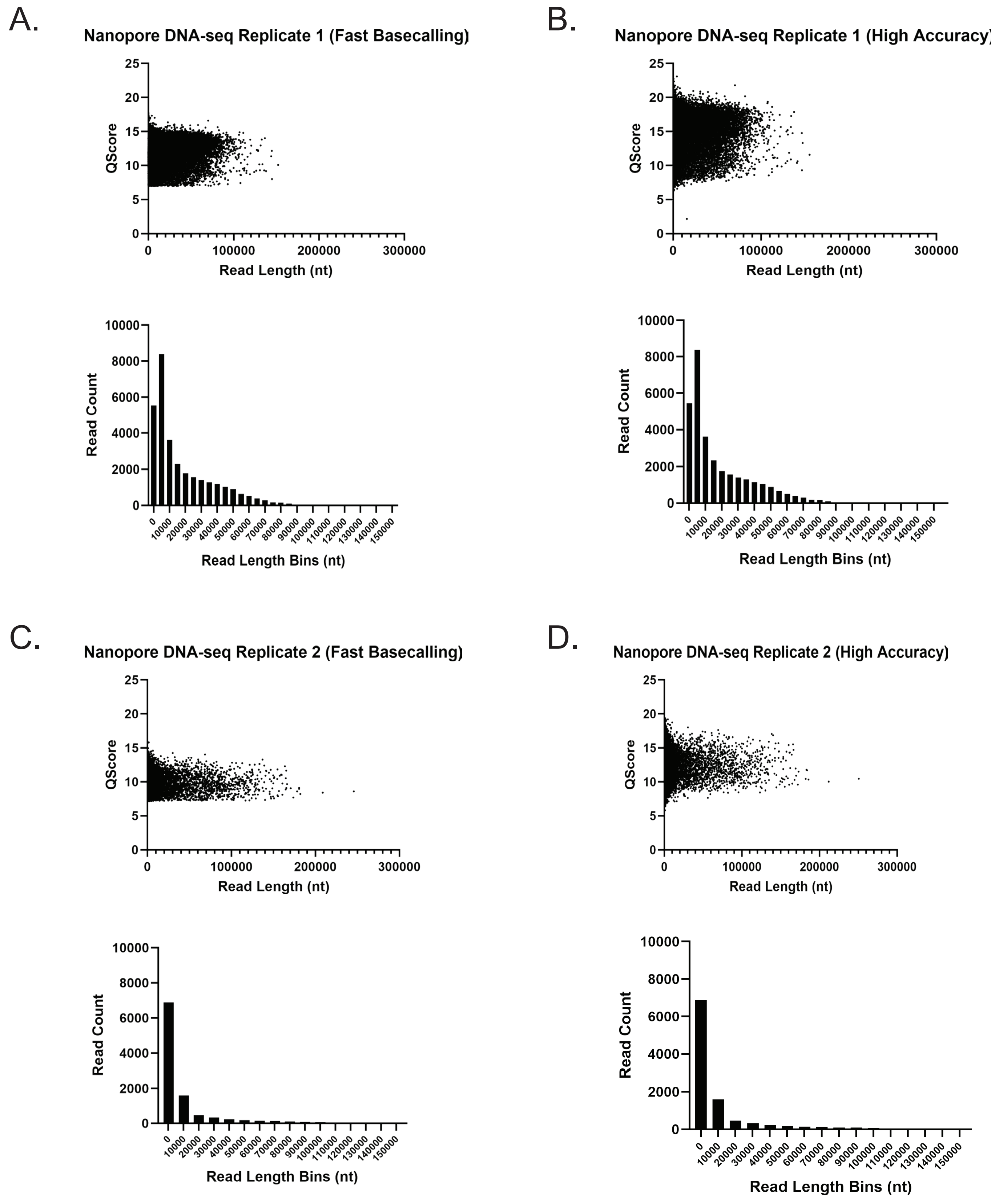


**Supplementary Figure 3: A comparison of fast basecalling and high accuracy basecalling for Nanopore DNA sequencing.** (A and B) Nanopore ligation sequencing reads from replicate one were basecalled using the fast basecalling algorithm (A) or the high accuracy basecalling algorithm (B). The Qscore vs. read length distribution is depicted as a scatter plot (top), and the read length and their respective counts are plotted as a histogram (bottom). (C and D) The same comparisons as described in A and B were applied to replicate two.

**Godin and Sebastian *et al*. Supplementary Figure 4**

A.

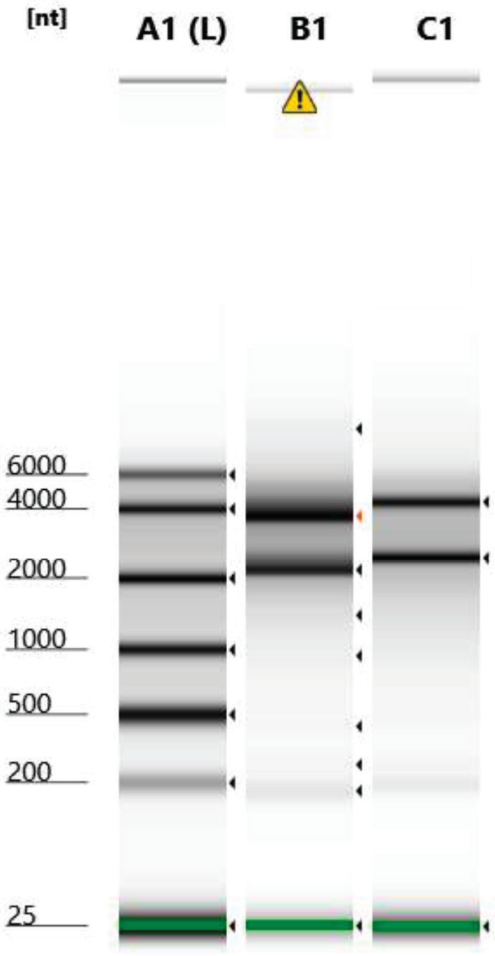


A1: Ladder
B1: RNA Nanopore Replicate 1
C1: RNA Nanopore Replicate 1
1:5 Dilution

B.

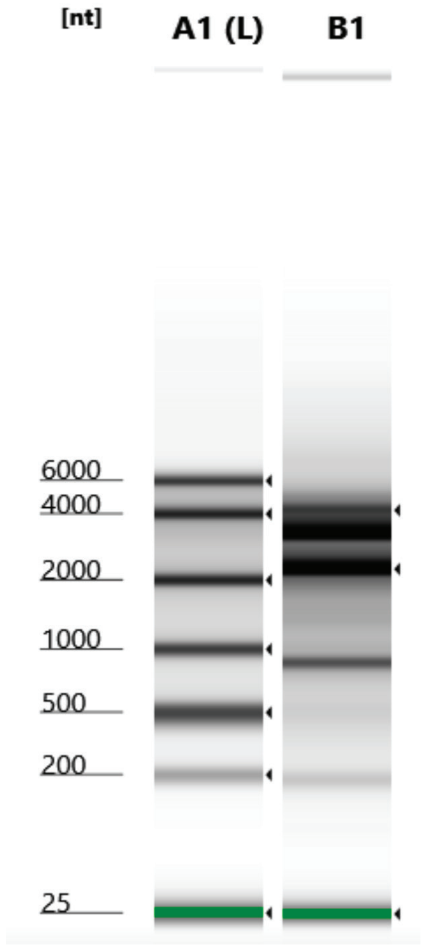


A1: Ladder
B1: RNA Nanopore
Replicate 2

**Supplementary Figure 4: Bioanalyzer results demonstrate that RNA samples are of high quality.** (A) Total RNA isolated for replicate one of Nanopore direct RNA sequencing was run on a Bioanalyzer for quality control purposes. The yellow hazard sign in lane B1 indicates that the markers ran outside of their standard position, leading to an edited RIN. The sample was also run at a 1:5 dilution. (B) The RNA sample used for replicate 2 of Nanopore direct RNA sequencing was run separately in the same way.

**Godin and Sebastian *et al*. Supp. Figure 5**

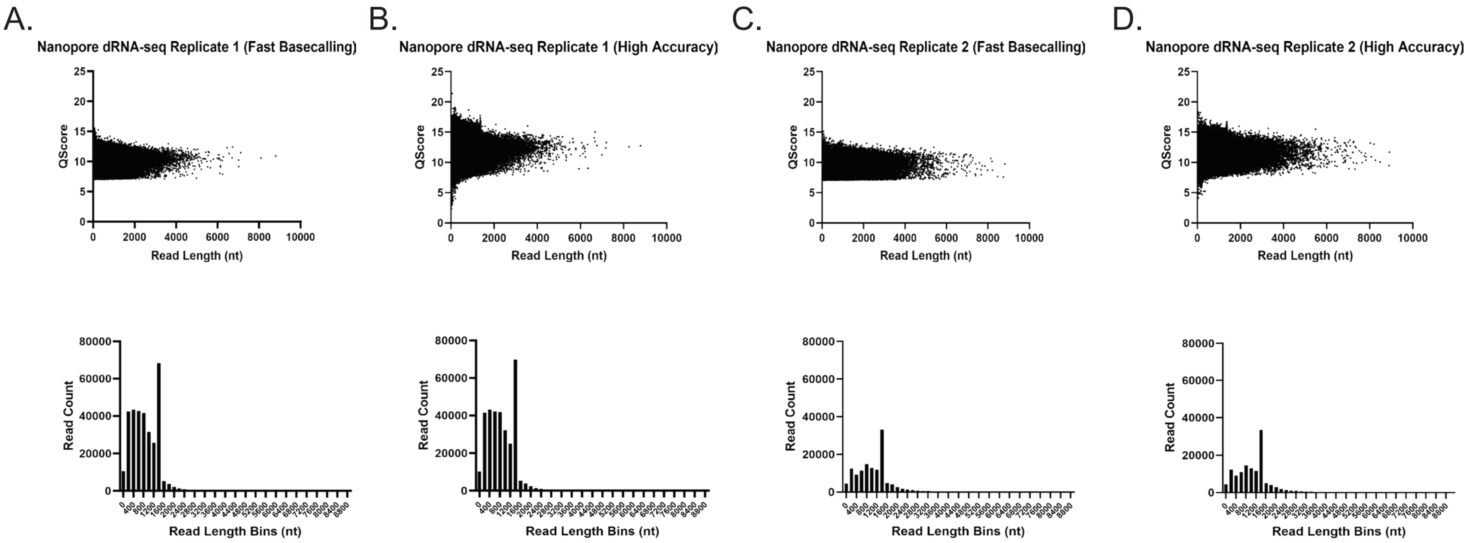


**Supplementary Figure 5: A comparison of fast basecalling and high accuracy basecalling for Nanopore direct RNA sequencing.** Nanopore direct RNA sequencing reads from replicate one (A and B) or two (C and D) were basecalled using the fast basecalling algorithm or the high accuracy basecalling algorithm. The Qscore vs. read length distribution is depicted as a scatter plot (top), and the read length and their respective counts are plotted as a histogram (bottom).

**Godin and Sebastian *et al*. Supplementary Figure 6**.

***P. yoelii* CSP Repeat Lengths**

**>Py17X_CSP-150AA**
DQGPGAPQGPGAPQGPGAPQGPGAPQGPGAPQGPGAPQGPGAPQGPGAPQGPGAP
QGPGAPQGPGAPQGPGAPQGPGAPQGPGAPQGPGAPQGPGAPQGPGAPQGPGAPQ
GPGAPQGPGAPQGPGAPQGPGAPQGPGAPQGPGAPQGPGA

**>Py17XNL_CSP-114AA_(6 D/PQGPGA units shorter than 17X)**
DQGPGAPQGPGAPQGPGAPQGPGAPQGPGAPQGPGAPQGPGAPQGPGAPQGPGAP
QGPGAPQGPGAPQGPGAPQGPGAPQGPGAPQGPGAPQGPGAPQGPGAPQGPGAPQ
GPGA

**>PyYM_CSP-108AA_(7 D/PQGPGA units shorter than 17X)**
DQGPGAPQGPGAPQGPGAPQGPGAPQGPGAPQGPGAPQGPGAPQGPGAPQGPGAP
QGPGAPQGPGAPQGPGAPQGPGAPQGPGAPQGPGAPQGPGAPQGPGAPQGPGA

**Supplementary Figure 6: The CSP central repeat region length varies across sequenced *P. yoelii* strains.** The amino acid sequence for the central repeat region of circumsporozoite protein (CSP) is shown for *P. yoelii* 17X, *P. yoelii* 17XNL, and *P. yoelii* YM.
