## Supplemental File 1 for "Long-Read Genome Assembly and Gene Model Annotations for the Rodent Malaria Parasite *Plasmodium yoelii* 17XNL": combine_prokka_braker_gff.py

```
import csv, sys, os
import gffutils
from collections import defaultdict

#
# This script produces a modified gff3 that combines both braker2 and prokka-rblast annotations
# for apicoplast and mitochondrial chromosomes.
#

sub_feats = ['start_codon', 'mRNA', 'exon', 'CDS', 'intron', 'stop_codon', 'five_prime_utr', 'three_prime_utr', 'transcription_start_site', 'transcription_end_site']


def read_ids(fname):
    """
    Reads a tab-delimited idmap file and returns a dictionary of list.
    """
    trans_store = defaultdict(dict)
    stream = csv.DictReader(open(fname), delimiter="\t")
    for row in stream:
        protein_id = "query_pid"
        pid = row.get(protein_id)
        trans_store[pid] = row
    return trans_store


def get_genes(gff, dbname):
    """
    Extract all gene names from a gff file
    """

    store = dict()
    dbname = dbname

    if not (os.path.exists(dbname)):
        # Create an sqlite3 database. This is a one time operation.
        db = gffutils.create_db(gff, dbname)

    db = gffutils.FeatureDB(dbname, keep_order=True)
    # Print all genes
    for g in db.all_features(featuretype='gene'):
        gid = g.id
        name = g.attributes['Name'][0]
        store[name] = gid
    return store


def print_all_feats(gene, dbname, include_gene=True):
    db = gffutils.FeatureDB(dbname, keep_order=True)

    if include_gene:
        # print gene
        print(db[gene])

    # Get all subfeatures for a gene
    for feat in sub_feats:
        for i in db.children(gene, featuretype=feat, order_by='start'):
            print(i)


def modify_gene_attr(gid, gname, source, qcovs, pident, matched_protein, org_gid=""):
    if gname is None or gname == "":
        # new_attr = f'ID={gid};gene_id={gid};gene_name={gid};Name={gid};best_match=augustus; original_gene_id={org_gid}'
        new_attr = f'ID={gid};gene_id={gid};gene_name={gid};Name={gid};best_match=augustus'
    else:
        new_attr = f'ID={gid};gene_id={gid};gene_name={gid};Name={gname};best_match={matched_protein};query_coverage={qcovs};percentage_identity={pident}'

    return new_attr


def modify_transcript_attr(tid, gname, source, qcovs, pident, evidence, matched_protein, org_tid=""):
    parent, transcript = tid.split(".")

    # if gname is None or gname == "":
    if evidence == "augustus":
        new_attr = f'ID={tid};Parent={parent};transcript_id={tid};gene_name={parent};Name={tid};best_match={evidence}'
    elif evidence == "locus":
        trans_id = ".".join([gname, transcript])
        new_attr = f'ID={tid};Parent={parent};transcript_id={tid};gene_name={gname};Name={trans_id};best_match={evidence}'

    else:
        trans_id = ".".join([gname, transcript])
        new_attr = f'ID={tid};Parent={parent};transcript_id={tid};gene_name={parent};Name={trans_id};best_match={matched_protein};query_coverage={qcovs};percentage_identity={pident}'

    return new_attr


def modify_attr(new_id, tid, gname, gid, org_tid=""):
    parent = tid

    if gname is None or gname == "":
        # new_attr = f'Parent={parent};transcript_id={parent}; original_transcript_id={org_tid}'
        new_attr = f'ID={new_id};Name={new_id};Parent={parent};transcript_id={parent}'
    else:
        new_attr = f'ID={new_id};Name={new_id};Parent={parent};transcript_id={tid};gene_name={gname}'

    return new_attr


def get_vals(feat):
    chrom = str(feat.chrom)
    start = str(feat.start)
    end = str(feat.end)
    strand = feat.strand
    frame = feat.frame
    return (chrom, start, end, strand, frame)


def make_gff_rows(feat, feat_id, dbh, gname, qcovs, pident, matched_protein):
    new_gid, new_tid, new_attr = "", "", ""

    source = "prokka"

    fid = feat_id + "_" + feat if feat == "gene" or feat == "mRNA" else feat_id

    pgene = dbh[fid]

    feat = str(pgene).split("\t")[2] if feat != "gene" or feat != "mRNA" else feat

    feat_vals = get_vals(pgene)
    chrom, start, end, strand, frame = feat_vals

    feats = "\t".join([str(chrom), source, feat, str(start), str(end), ".", strand, str(frame)])

    if feat == "gene":
        new_gid = chrom + str(feat_id.split('_')[1])
        new_attr = modify_gene_attr(new_gid, gname, source, qcovs, pident, matched_protein)

    elif feat == "mRNA":
        evidence = "rblast"
        new_tid = chrom + str(gid.split('_')[1]) + ".1"
        new_attr = modify_transcript_attr(new_tid, gname, source, qcovs, pident, evidence, matched_protein)

    else:
        new_id = chrom + str(gid.split('_')[1]) + ".1" + f"-{feat}1"
        new_tid = chrom + str(gid.split('_')[1]) + ".1"
        new_attr = modify_attr(new_id, new_tid, gname, new_gid, org_tid="")

    new_row = "\t".join([feats, new_attr])
    print(new_row)


def write_gff(gid, vals, braker_genes, braker_db, prokka_db, prokka_gff):
    gname = vals['gene_name']
    qcovs = vals['qcovs']
    pident = vals['qcovs']
    matched_protein = vals['matched_protein']

    # Get braker database handle
    db = gffutils.FeatureDB(braker_db, keep_order=True)

    # Create prokka gff database
    if not (os.path.exists(prokka_db)):
        # Create an sqlite3 database. This is a one time operation.
        dbp = gffutils.create_db(prokka_gff, prokka_db)

    # prokka db handle
    dbp = gffutils.FeatureDB(prokka_db, keep_order=True)

    # Check if the gene is already annotated by braker

    braker_id = braker_genes[gname] if gname in braker_genes else ""

    if braker_id != "":
        # print the rows from braker gff3
        # print("B", braker_id)
        print_all_feats(braker_id, braker_db, include_gene=True)
        return ()

    #
    # Modify rows from prokka gff file
    #

    # Modify gene row
    make_gff_rows("gene", gid, dbp, gname, qcovs, pident, matched_protein)

    # Modify mRNA row
    make_gff_rows("mRNA", gid, dbp, gname, qcovs, pident, matched_protein)

    # Modify CDS or other subfeatures
    make_gff_rows("sub", gid, dbp, gname, qcovs, pident, matched_protein)


if __name__ == "__main__":
    map_file = sys.argv[1]  # "annotated_gene_names.txt"
    braker_gff = sys.argv[2]  # "API_hybrid.gff3"
    prokka_gff = sys.argv[3]  # "PROKKA_11142022.gff"

    # Read rblast based annotation text file
    trans_map = read_ids(map_file)

    # Databases that stores gff files from braker and prokka annotation
    braker_db = "braker.db"
    prokka_db = "prokka.db"

    # Get the genes annotated by braker
    genes = get_genes(braker_gff, braker_db)

    # Produce a modified gff combining braker and prokka predictions.
    for gid, vals in sorted(trans_map.items()):
        write_gff(gid, vals, braker_genes=genes, braker_db=braker_db, prokka_db=prokka_db, prokka_gff=prokka_gff)
```
