## Supplemental File 1 for "Long-Read Genome Assembly and Gene Model Annotations for the Rodent Malaria Parasite *Plasmodium yoelii* 17XNL": lift_nano_genes.py

```
"""
This script adds genes from nanopore modelto the hybrid model.
The criteria for selecting genes that are to be added is
    1) The start-codon and stop-codon of a 17X gene is not present in the hybrid model but are annotated
     for a gene in nanopore model. This may be because the gene is not predicted in hybrid model (eg:CSP)
     or the gene is wrongly annotated in hybrid model (eg:NFU1). Such genes are lifted from nanopore model
     to the hybrid model.

Result:
This script added 80 genes from nanopore-only model to the hybrid model.
17 of these were already present in hybrid model but the start-codon and stop-codon were
 not matching with the corresponding 17X gene. The hybrid annotation copy of these genes
 were removed and nanopore-only annotations that matched with the 17X gene were kept for these 17 genes.

 IN addition the script writes out text files containing newids and corresponding nanopore-ids,
 duplicated hybrid gene ids and unique hybrid gids.

"""

import csv, sys, os
import gffutils
import re

sub_feats = ['start_codon', 'stop_codon', 'five_prime_utr', 'three_prime_utr',
             'transcription_start_site', 'transcription_end_site', 'mRNA', 'exon', 'CDS', 'intron']


def store_bed(bedfile):
    store = dict()

    stream = csv.reader(open(bedfile), delimiter="\t")

    for row in stream:
        chrom, start, end, name, strand = row[0], row[1], row[2], row[3], row[5]

        if strand == "-":
            start, end = end, int(start) + 1

        if strand == "+":
            start = int(start) + 1

        d = dict()
        d['chrom'] = chrom
        d['start'] = start
        d['end'] = end
        d['strand'] = strand
        d['name'] = name

        store[name] = d
    return store


def print_all_feats(gene, dbname, include_gene=True):
    vals = list()

    db = gffutils.FeatureDB(dbname, keep_order=True)

    if include_gene:
        # store gene
        # print(db[gene])
        vals.append(db[gene])

    # Get all subfeatures for a gene
    for feat in sub_feats:
        for i in db.children(gene, featuretype=feat, order_by='start'):
            # print(i)
            vals.append(i)

    return vals


def store_gff3(fname, dbname):
    """
    Stores the gff3 file in a database and returns the database handle.
    """
    dbname = dbname

    if not (os.path.exists(dbname)):
        # Create an sqlite3 database. This is a one time operation.
        db = gffutils.create_db(fname, dbname)

    db = gffutils.FeatureDB(dbname, keep_order=True)

    return db


def get_starts_stops(dbh):
    """
    Extract start and stop codons from a database.
    Input parameter is the database handle.
    Returns a list of all start codons and a list of stop codons.
    """

    starts, stops, matched = dict(), dict(), dict()

    for g in dbh.all_features(featuretype='gene'):
        gid = g.id
        gstrand = g.strand

        # Get the matched gene in 17X
        matched_17X = dbh[gid].attributes['best_match'][0]
        matched_17X = re.sub('\.\d-p\d', '', matched_17X)
        matched[matched_17X] = gid

        for i in dbh.children(gid, featuretype='start_codon', order_by='start'):
            s = i.start
            e = i.end
            chrom = i.chrom
            start_codon_start = chrom + ":" + str(s)
            start_codon_end = chrom + ":" + str(e)

            key = start_codon_start if gstrand == "+" else start_codon_end

            # starts[start_codon_start]=gid if gstrand == "+" else starts[start_codon_end] = gid

            starts[key] = gid

        for i in dbh.children(gid, featuretype='stop_codon', order_by='start'):
            s = i.start
            e = i.end
            chrom = i.chrom
            stop_codon_start = chrom + ":" + str(s)
            stop_codon_end = chrom + ":" + str(e)
            # stops[stop_codon_end] = gid if gstrand == "+" else stops[stop_codon_start] = gid
            key = stop_codon_end if gstrand == "+" else stop_codon_start
            stops[key] = gid

    return starts, stops, matched


def get_genes(gff, dbname):
    """
    Extract all gene names from a gff file
    """

    store = list()
    dbname = dbname

    if not (os.path.exists(dbname)):
        # Create an sqlite3 database. This is a one time operation.
        db = gffutils.create_db(gff, dbname)

    db = gffutils.FeatureDB(dbname, keep_order=True)
    # Print all genes
    for g in db.all_features(featuretype='gene'):
        # gid = g.id
        # name = g.attributes['Name'][0]
        # store[name] = gid
        store.append(g.id)
    return store


def generate_uid(last_gids, chrom):
    gid = last_gids[chrom]
    num = gid.replace(chrom, '')
    new_num = int(num) + 1
    new_num = str(new_num).zfill(5)
    new_id = f"{chrom}{new_num}"
    last_gids[chrom] = new_id
    return last_gids, new_id


def get_last_gid(ann, dbname):
    """
    Extract last geneid for each chromsome
    Returns a dictionary with chromosome as key and last gid as the value.
    """
    last = dict()
    genes = sorted(get_genes(ann, dbname))
    for g in genes:
        chrom, gnum = g.split("_")
        gnum = str(gnum)
        cnum = gnum[:2] if gnum[0].isdigit() else gnum[:3]
        chrom_id = f"{chrom}_{cnum}"
        last[chrom_id] = g
    return last


if __name__ == "__main__":
    hybrid_ann = sys.argv[1]  # "17XNL_PSU_hybrid.gff3"
    nano_ann = sys.argv[2]  # "17XNL_PSU_nano.gff3"
    py17x_model = sys.argv[3]  # "17X_transcripts_sensitive.bed"

    hybrid_genes = list()
    added_gnames = dict()
    duplicated_genes = list()
    added_gene_feats = list()

    # Store 17X transcript model bed file
    store_17X = store_bed(py17x_model)

    # Databases that stores gff files from hybrid and nano annotation
    hybrid_dbname = "hybrid.db"
    nano_dbname = "nano.db"

    nano_db = store_gff3(nano_ann, nano_dbname)
    hybrid_db = store_gff3(hybrid_ann, hybrid_dbname)

    # extract start codons and stop codons of all hybrid gene models
    hybrid_starts, hybrid_stops, hybrid_matched = get_starts_stops(hybrid_db)

    # extract start codons and stop codons of all nanopore-only gene models
    nano_starts, nano_stops, nano_matched = get_starts_stops(nano_db)

    # get the last hybrid gene ids for each chromosome.
    # Newly generated gids will be checked against these to make sure uniqueness.
    last_gids = get_last_gid(hybrid_ann, hybrid_dbname)

    # Go through 17X model and check if a gene exist in hybrid model at the same coordinate.

    for k, v in store_17X.items():
        vals = store_17X[k]
        chrom = vals['chrom']
        start = vals['start']
        end = vals['end']
        strand = vals['strand']
        name = vals['name']
        gname = re.sub('\.\d+', '', name)

        scoord = chrom + ":" + str(start)
        ecoord = chrom + ":" + str(end)

        # Cases where the Py17X start and end has no match in hybrid model
        if scoord not in hybrid_starts and ecoord not in hybrid_stops:
            # print(chrom, start, end, strand, k)

            # Check the hybrid missing gene is annotated in nanopore-only model.
            if scoord in nano_starts and ecoord in nano_stops:
                # print("***", chrom, start, end, strand, k, nano_starts[scoord], nano_stops[ecoord])

                # Generate a new unique id
                last_gids, new_id = generate_uid(last_gids, chrom)

                # Get all the annotated features of the gene from nanopore only model
                nano_gid = nano_starts[scoord]
                nano_gname = nano_db[nano_gid].attributes['Name'][0]
                added_gnames[new_id] = [nano_gid, nano_gname]

                # Check if gid is duplicated by the addition of new gene.
                duplicated_genes.append(hybrid_matched[gname]) if gname in hybrid_matched else duplicated_genes

                # Add the missing gene from nanoporeo-nly model replacing its id with the new id
                info = print_all_feats(nano_gid, nano_dbname, include_gene=True)
                for i in info:
                    a = str(i).replace(nano_gid, new_id)
                    a = a + f";note=annotation added from nanopore-only model"
                    # print(a)
                    added_gene_feats.append(a)

    # Newly added gene names
    fa = open("added_genes.txt", "w")
    h = "\t".join(['new_id', 'nano_id', 'name'])
    fa.write(h)
    fa.write("\n")
    # print(h)
    for k, v in added_gnames.items():
        out = "\t".join([k, v[0], v[1]])
        fa.write(out)
        fa.write("\n")
        # print(out)

    # print("--------------------")
    # Print hybrid gids of duplicated genes
    fd = open("duplicated_genes_hybrid_gids.txt", "w")
    for d in duplicated_genes:
        fd.write(d)
        fd.write("\n")
        # print(d)

    # Create a new gff3 file with the new gene addition
    #
    fn = open("hybrid_org_uniq_gids.txt", "w")
    header = "##gff-version 3"
    print(header)
    # Print newly added genes.
    for item in added_gene_feats:
        print(item)
    # Print the remaining genes
    for g in hybrid_db.all_features(featuretype='gene'):
        gid = g.id
        fn.write(gid)
        fn.write("\n")
        if gid not in duplicated_genes:
            ginfo = print_all_feats(gid, hybrid_dbname, include_gene=True)
            for i in ginfo:
                print(i)
```
