## Supplemental File 1 for "Long-Read Genome Assembly and Gene Model Annotations for the Rodent Malaria Parasite *Plasmodium yoelii* 17XNL": update_fasta.sh

```
PROT=17XNL_PSU_hybrid_proteins.fa 
CDS=17XNL_PSU_hybrid_cds.fa
TRANS=17XNL_PSU_hybrid_transcripts.fa 

PROT_FIX=MIT_fix_proteins.fa
CDS_FIX=MIT_fix_cds.fa
TRANS_FIX=MIT_fix_transcripts.fa

HEADER=Py17XNL_MIT

#Modify protein fasta file
echo
cat $PROT | grep ">" | grep -v ${HEADER} | sed 's/>//' >list.txt
seqtk subseq -l 60 ${PROT} list.txt >list.fa
cat list.fa ${PROT_FIX} >$PROT

# Modify cds fasta file
echo
cat $CDS | grep ">" | grep -v ${HEADER} | sed 's/>//' >list.txt
seqtk subseq -l 60 ${CDS} list.txt >list.fa
cat list.fa ${CDS_FIX} >$CDS

# Modify transcript fasta file
echo
cat ${TRANS} | grep ">" | grep -v ${HEADER} | sed 's/>//' >list.txt
seqtk subseq -l 60 ${TRANS} list.txt >list.fa
cat list.fa ${TRANS_FIX} >$TRANS
```
