## Supplemental File 1 for "Long-Read Genome Assembly and Gene Model Annotations for the Rodent Malaria Parasite *Plasmodium yoelii* 17XNL": assign.sh

```
set -ue

# Steps are

# 1) Map assembly to reference
# 2) Extract primary aligned contigs
# 3) Get the % coverage of each contig
# 4) Filter lowly covered contigs(<2% of the reference) and assign chromosome name to contigs

ASSEMBLY=$1
REF=$2
OUT_PREFIX=$3

CPU=15
#SRC=scripts/misc

# Directory to hold alignments
BAM=bam
mkdir -p $BAM

#OUTPUT
OUT=${BAM}/${OUT_PREFIX}"_asm"
OUT_PRIMARY=${BAM}/${OUT_PREFIX}"_asm_primary"

OUTBAM=${OUT}".bam"
OUT_PRIMARY_BAM=${OUT_PRIMARY}".bam"

# 1. Map assembly to refernce.
minimap2 -t $CPU -ax asm5 $REF $ASSEMBLY |samtools sort >$OUTBAM
samtools index $OUTBAM

# 2. Extract primary aligned contigs
samtools view -h -b -F 4 -F 2304 $OUTBAM >$OUT_PRIMARY_BAM
samtools index $OUT_PRIMARY_BAM

# 3. Get assembly coverages corresponding to reference.
cat $REF | grep ">" |cut -d ' ' -f 1 | sed 's/>//' >chroms.txt
python parse_aln.py $OUT_PRIMARY_BAM chroms.txt >${OUT_PREFIX}"_coverages.txt"
rm -f chroms.txt
```
