## Supplemental File 1 for "Long-Read Genome Assembly and Gene Model Annotations for the Rodent Malaria Parasite *Plasmodium yoelii* 17XNL": circularize.sh

```
set -uex

#
# Circularize API in pyoelli 17XNL assembly
#


# Assembly fasta file with contigs
ASM=asm/17XNL_PSU_v3.fa

# Reads used to generate the assembly
READS=data/ccs.fq 

# Sequence to be extracted from fasta file and
# needs to be circularized. This id should match fasta header.
SID=PY17XNL_API

# Output directory
OUT=api_circ

CPU=20

# Extract API
samtools faidx $ASM $SID >api.fa

# circlator
circlator all --threads $CPU api.fa  $READS $OUT
```
