## Supplemental File 1 for "Long-Read Genome Assembly and Gene Model Annotations for the Rodent Malaria Parasite *Plasmodium yoelii* 17XNL": modify_header.py

```
from Bio import SeqIO
import sys


def store_names(fname):
    store = dict()
    for line in open(fname):
        old, new = line.split("\t")
        store[old.strip()] = new.strip()
    return store


def change_header(fasta_file, names):
    name_store = store_names(names)
    for r in SeqIO.parse(fasta_file, "fasta"):
        new_id = name_store[r.id] if r.id in name_store else r.id
        print(">" + new_id)
        print(r.seq)


if __name__ == "__main__":
    #fasta = "selected_contigs.fa"
    #names = "contig_map.txt"
    fasta = sys.argv[1]
    names = sys.argv[2]
    change_header(fasta, names)
```
