## Supplemental File 1 for "Long-Read Genome Assembly and Gene Model Annotations for the Rodent Malaria Parasite *Plasmodium yoelii* 17XNL": parse_aln.py

```
import pysam, sys
from collections import defaultdict

NAME_PREFIX = "PY17XNL"

# coverage cut off in percentage to filter alignments
CUTOFF = 2.0


def parse(bam, chroms):
    bamfile = pysam.AlignmentFile(bam, "rb")

    aln_len, ref_len, covs = dict(), dict(), dict()

    for chr in open(chroms, "r"):
        chr = chr.strip()
        ref_len[chr] = bamfile.get_reference_length(chr)

        for read in bamfile.fetch(chr):
            if read.is_unmapped: continue
            key = ":".join([read.query_name, read.reference_name])
            aln_len[key] = aln_len[key] + read.query_alignment_length if key in aln_len else read.query_alignment_length

    # Calculate coverage
    for k, v in aln_len.items():
        query, ref = k.split(":")
        rlen = ref_len[ref]
        cov = round((v / rlen) * 100, 2)
        covs[k] = cov

    # Make a list of lists and sort it
    vals = list()
    res = defaultdict(list)
    for k in sorted(aln_len, key=aln_len.get, reverse=True):
        query, ref = k.split(":")
        mapped_len = aln_len[k]
        mapped_cov = covs[k]
        chrom_len = ref_len[ref]
        # Filter contigs with low coverage.
        if mapped_cov < CUTOFF: continue
        vals.append([query, ref, mapped_len, chrom_len, mapped_cov])

    sorted_vals = sorted(vals, key=lambda y: (y[1], -y[4]))

    # convert the list of lists into a dictionary keyed by chromosome
    for item in sorted_vals:
        chrom = item[1]
        res[chrom].append(item)

    # for k, v in res.items():
    #     print(k,v)
    # 1/0
    header = "\t".join(['query', 'chrom', 'mapped_len', 'chrom_len', 'cov_perc', 'assignment'])
    print(header)
    for chrom, vals in res.items():
        for idx, v in enumerate(vals):
            contig, chrom, mapped, ref_len, cov = v

            new_name = chrom if idx == 0 else chrom + f"_unloc{idx}"

            res = "\t".join([contig, chrom, str(mapped), str(ref_len), str(cov), new_name])
            print(res)


if __name__ == "__main__":
    #bam = "canu_asm.bam"
    #chroms = "chroms.txt"
    bam = sys.argv[1]
    chroms = sys.argv[2]
    parse(bam, chroms)
```
