## Supplemental File 1 for "Long-Read Genome Assembly and Gene Model Annotations for the Rodent Malaria Parasite *Plasmodium yoelii* 17XNL": rearrange.sh

```
set -uex
#
# Circlator merged the overlapping regions.
# But circlator did not cirularize correctly
# So,we mapped the circlator output to 17X
# and obtained the coordinates and rearranged it.

# Circlator output
FASTA=api_circ.fa
REF=refs/PlasmoDB-56_Pyoeliiyoelii17X_Genome.fasta

# Map circlator output to reference
minimap2 -ax asm5 $REF $FASTA |samtools sort >api_circ.bam
samtools index api_circ.bam

# Extract sequences to rearrange based on mapping coordinates.
samtools faidx ${FASTA} 'PY17XNL_API:2974-34324' >seq1.fa
samtools faidx ${FASTA} 'PY17XNL_API:1-2973' >seq2.fa

# Merge the sequences.
echo ">PY17XNL_API" >tmp1.fa
(cat seq1.fa | grep -v ">" & cat seq2.fa | grep -v ">")  >>tmp1.fa
seqtk seq -A -l 70  tmp1.fa >api_merge.fa
samtools faidx api_merge.fa
rm -f tmp1.fa
```
