## Supplemental File 1 for "Long-Read Genome Assembly and Gene Model Annotations for the Rodent Malaria Parasite *Plasmodium yoelii* 17XNL": consensus.py

```
from itertools import *
import csv, re
from Bio import SeqIO

def main(fname):

    stream = open("refs/17X_PLD.fa")
    stream = SeqIO.parse(stream, "fasta")
    data ={}
    for record in stream:
        data[record.id] = record.seq

    stream = open(fname, 'r')

    def transform(row):
        chrom, pos, ref, depth, bases, quals = row
        bases = bases.replace("^", "").replace("]", "").replace("$", "")
        bases = bases.replace(".", ref).replace(",", ref).upper()

        return chrom, pos, ref, depth, bases, quals

    def keep(row):
        chrom, pos, ref, depth, bases, quals = row
        return ref =='N' or bases == 'N'

    stream = csv.reader(stream, delimiter='\t')
    stream = map(transform, stream)

    #stream = filter(keep, stream)
    #stream = islice(stream, 100)

    # Pattern to split bases
    patt = re.compile(r'\+\d+')

    last, coll = None, []
    for row in stream:
        chrom, pos, ref, depth, bases, quals = row

        anchor = data[chrom]

        if chrom != last:
            if last:
                line = "".join(coll)
                print(f">{last}")
                print(f"{line}")

            last = chrom
            coll = []

        depth = int(depth)

        ref = anchor[int(pos)-1]

        base = ref

        if depth == 0 and ref == 'N':
            base = ref
        elif depth == 1 and ref == 'N':
            base = ref
        elif depth == 1 and bases == '*':
            continue
        elif depth == 1 and len(bases) == 1:
            base = bases
        elif depth == 1 and '+' in bases:
            nref, seq = patt.split(bases)
            num = int(len(seq))
            # Skip large insertions
            if num > 20000:
                print(chrom, pos, num)
                base = ref
            else:
                base = ref + seq

        coll.append(base)


    line = "".join(coll)
    print(f">{last}")
    print(f"{line}")

if __name__ == '__main__':
    fname = 'cons/pileup.txt'
    main(fname)
```
