## Supplemental File 1 for "Long-Read Genome Assembly and Gene Model Annotations for the Rodent Malaria Parasite *Plasmodium yoelii* 17XNL": add_gids.py

```
"""
This script adds geneids to an input file based on a common column.
"""

import sys, csv


def read_idmap(fname):
    store = dict()
    stream = csv.reader(open(fname), delimiter="\t")
    header = next(stream)
    for row in stream:
        comm_id = row[3].strip()
        id_match = row[5].strip()
        store[comm_id] = id_match
    return store


def join(idmap="", infile="", by_col=1):
    id_store = read_idmap(idmap)
    stream2 = csv.reader(open(infile), delimiter="\t")
    col = by_col - 1

    for row in stream2:
        gid = row[col].strip()
        gmatch = id_store.get(gid, "")
        out = "\t".join(row)
        out = "\t".join([out, gmatch])
        print(out)


if __name__ == "__main__":
    idmap = sys.argv[1]  # "gene_match.txt"
    infile = sys.argv[2]  # "UTR_length_hybrid_5P.txt"
    id_col = int(sys.argv[3])  # 5  # column by which files needs to be joined

    join(idmap=idmap, infile=infile, by_col=id_col)
```
