## Supplemental File 1 for "Long-Read Genome Assembly and Gene Model Annotations for the Rodent Malaria Parasite *Plasmodium yoelii* 17XNL": add_matched_prot.py

```
import sys,csv


def store_data(fname):

    store = dict()

    stream = csv.reader(open(fname), delimiter="\t")
    header = next(stream)
    for row in stream:
        chrom = row[0]
        start = int(row[1])
        end = int(row[2])
        try:
            xnl_gid, xnl_pid,xnl_pname,match = row[3:]

        except:
            print(row)
            sys.exit()
        store[match] = [chrom, start, end, xnl_gid, xnl_pid, xnl_pname]
    return store


def join(f1, f2):
    # store intersection file.
    match = store_data(f1)
    match_17XNL =""


    stream2 = csv.reader(open(f2), delimiter="\t")

    header = "\t".join(["17X_PLD_chrom", "17X_PLD_pos","17X_allele", "17XNL_allele","gene_id", "17X_PLD_prot", "17XNL_PSU_gid","17XNL_PSU_pid", "17XNL_PSU_pname" ])
    print(header)

    for row in stream2:
        chrom = row[0]
        pos = row[1]
        ref_allele = row[3]
        alt_allele = row[4]
        info = row[18]
        vals = info.split(";")
        gid = vals[2].replace("gene_id=", "")
        pid = vals[3].replace("protein_source_id=", "")
        if pid in match:
            #print(match[pid])
            #xnl_chrom = match[pid][0]
            #xnl_start = match[pid][1]
            #xnl_end = match[pid][2]
            xnl_gid, xnl_pid,xnl_pname =match[pid][3], match[pid][4],match[pid][5]
            match_17XNL = "\t".join([xnl_gid, xnl_pid,xnl_pname])
        else:

            match_17XNL = "\t".join(["", "",""])

        out = "\t".join([chrom, str(pos), ref_allele, alt_allele, gid, pid, match_17XNL])
        print(out)


if __name__ == "__main__":
    match = sys.argv[1] #"prot_match.txt"
    intersect = sys.argv[2] #"intersect.txt"
    join(match, intersect)
```
