## Supplemental File 1 for "Long-Read Genome Assembly and Gene Model Annotations for the Rodent Malaria Parasite *Plasmodium yoelii* 17XNL": attributes_gff.py

```
# This script extract attributes from 9th column corresponding to a feature.
# usage: python attributes.py <gff file> <feature in 3rd column> <: separated list of attributes to be printed>
# example : python attributes_gff.py criollo_v2.gff  gene ID:GeneID:Name

import sys,re
from collections import defaultdict


input_file = sys.argv[1]
feature = sys.argv[2]
input_list = sys.argv[3]

attributes = input_list.split(":")

infoDict = defaultdict(list)
gff_file = open(input_file, "r")
for stream in gff_file:
    if stream.startswith("#"):
        pass
    else:
        line = stream.split("\t")
      
        # similarity with 3rd column  
        if(re.search(feature,line[2])):
        #if line[2] == feature:   #exact match with 3rd column
            (chrom,start,end)=(line[0],line[3],line[4])
            arr = line[8].split(";")
            attr = set()

            for i in range(len(attributes)):
                key = attributes[i]
                attr.update([s.strip() for s in arr if key in s])
            coords="\t".join([chrom,start,end])
            attr_list=list(attr)
            for j in range(0,len(attributes)):
                 feat =attributes[j]
                 found = 0
                 for i in range(0,len(attr_list)):
                    if re.search(feat,attr_list[i]):
                        found =1
                        break
                 if found == 0:
                    attr_list.append(" ".join([feat,"\"None\""]))
            vals= "\t".join(sorted(attr_list))
            outline="\t".join([coords,vals])
            print(outline)
#        print(out)
```
