## Supplemental File 1 for "Long-Read Genome Assembly and Gene Model Annotations for the Rodent Malaria Parasite *Plasmodium yoelii* 17XNL": DotPrep.py

```
#! /usr/bin/env python

# Author: Maria Nattestad
#

# This script prepares a nucmer output delta file for visualization in Dot
# Parts of this code is adapted from Assemblytics unique anchor filtering


import argparse
import gzip
import time
import numpy as np
import operator
import re

def run(args):
	filename = args.delta
	unique_length = args.unique_length
	output_filename = args.out
	keep_small_uniques = True
	max_overview_alignments = args.overview

	# Read through the file and store information indexed by Query sequence names
	header_lines_by_query, lines_by_query = getQueryRefCombinations(filename)

	# Figure out which alignments contain sufficient unique anchor sequences
	unique_alignments = calculateUniqueness(header_lines_by_query, lines_by_query, unique_length, keep_small_uniques)

	# Write a filtered delta file, and coordinate files with uniqueness tags
	reference_lengths, fields_by_query = writeFilteredDeltaFile(filename, output_filename, unique_alignments, unique_length, header_lines_by_query)
	
	index_for_dot(reference_lengths, fields_by_query, output_filename, max_overview_alignments)


def scrub(string):
	return string.replace(",","_").replace("!","_").replace("~","_").replace("#", "_")


def getQueryRefCombinations(filename):
	print("header from delta file:")
	
	try:
		f = gzip.open(filename, 'rt')
		print(f.readline().strip())
	except:
		f = open(filename, 'r')
		print(f.readline().strip())

	# Ignore the first two lines for now
	print(f.readline().strip())

	linecounter = 0

	current_query_name = ""
	current_header = ""

	lines_by_query = {}
	header_lines_by_query = {}

	before = time.time()

	for line in f:
		if line[0]==">":
			linecounter += 1
			current_header = line.strip()
			current_query_name = scrub(current_header.split()[1])
			
			if header_lines_by_query.get(current_query_name, None) == None:
				lines_by_query[current_query_name] = []
				header_lines_by_query[current_query_name] = []
		else:
			fields = line.strip().split()
			if len(fields) > 4:
				# sometimes start and end are the other way around, but for this they need to be in order
				query_min = min([int(fields[2]),int(fields[3])])
				query_max = max([int(fields[2]),int(fields[3])])
				lines_by_query[current_query_name].append((query_min,query_max))
				header_lines_by_query[current_query_name].append(current_header)

	f.close()

	print("First read through the file: %d seconds for %d query-reference combinations" % (time.time()-before,linecounter))
	
	return (header_lines_by_query, lines_by_query)

def calculateUniqueness(header_lines_by_query, lines_by_query, unique_length, keep_small_uniques):
	before = time.time()
	unique_alignments = {}
	num_queries = len(lines_by_query)
	print("Filtering alignments of %d queries" % (num_queries))
	
	num_query_step_to_report = num_queries/100
	if num_queries < 100:
		num_query_step_to_report = num_queries/10
	if num_queries < 10:
		num_query_step_to_report = 1

	query_counter = 0

	for query in lines_by_query:
		unique_alignments[query] = summarize_planesweep(lines_by_query[query], unique_length_required = unique_length, keep_small_uniques = keep_small_uniques)
		query_counter += 1
		if (query_counter % num_query_step_to_report) == 0:
			print("Progress: %d%%" % (query_counter*100/num_queries))
	
	print("Progress: 100%")

	print("Deciding which alignments to keep: %d seconds for %d queries" % (time.time()-before,num_queries))

	return unique_alignments


def summarize_planesweep(lines,unique_length_required, keep_small_uniques=False):

	unique_alignments = []

	# If no alignments:
	if len(lines)==0:
		return []

	# If only one alignment:
	if len(lines) == 1:
		if keep_small_uniques == True or abs(lines[0][1] - lines[0][0]) >= unique_length_required:
			return [0]
		else:
			return []

	starts_and_stops = []
	for query_min,query_max in lines:
		starts_and_stops.append((query_min,"start"))
		starts_and_stops.append((query_max,"stop"))


	sorted_starts_and_stops = sorted(starts_and_stops,key=operator.itemgetter(0))

	current_coverage = 0
	last_position = -1
	sorted_unique_intervals_left = []
	sorted_unique_intervals_right = []
	for pos,change in sorted_starts_and_stops:
		if current_coverage == 1:
			sorted_unique_intervals_left.append(last_position)
			sorted_unique_intervals_right.append(pos)

		if change == "start":
			current_coverage += 1
		else:
			current_coverage -= 1
		last_position = pos


	linecounter = 0
	for query_min,query_max in lines:

		i = binary_search(query_min,sorted_unique_intervals_left,0,len(sorted_unique_intervals_left))

		exact_match = False
		if sorted_unique_intervals_left[i] == query_min and sorted_unique_intervals_right[i] == query_max:
			exact_match = True
		sum_uniq = 0
		while i < len(sorted_unique_intervals_left) and sorted_unique_intervals_left[i] >= query_min and sorted_unique_intervals_right[i] <= query_max:
			sum_uniq += sorted_unique_intervals_right[i] - sorted_unique_intervals_left[i]
			i += 1

		if sum_uniq >= unique_length_required:
			unique_alignments.append(linecounter)
		elif keep_small_uniques == True and exact_match == True:
			unique_alignments.append(linecounter)

		linecounter += 1

	return unique_alignments


def binary_search(query, numbers, left, right):
	#  Returns index of the matching element or the first element to the right
	
	if left >= right:
		return right
	mid = int((right+left)/2)
	

	if query == numbers[mid]:
		return mid
	elif query < numbers[mid]:
		return binary_search(query,numbers,left,mid)
	else: # if query > numbers[mid]:
		return binary_search(query,numbers,mid+1,right)


def natural_key(string_):
	"""See http://www.codinghorror.com/blog/archives/001018.html"""
	return [int(s) if s.isdigit() else s for s in re.split(r'(\d+)', string_)]

def writeFilteredDeltaFile(filename, output_filename, unique_alignments, unique_length, header_lines_by_query):
	before = time.time()
	f_out_delta = gzip.open(output_filename + ".uniqueAnchorFiltered_l%d.delta.gz" % (unique_length),'wt')
	
	try:
		f = gzip.open(filename, 'rt')
		header1 = f.readline()
	except:
		f = open(filename, 'r')
		header1 = f.readline()
		
	f_out_delta.write(header1) # write the first line that we read already
	f_out_delta.write(f.readline())
	
	linecounter = 0

	# For filtered delta file:
	list_of_unique_alignments = []
	alignment_counter = {}
	keep_printing = False

	# For coords:
	current_query_name = ""
	current_query_position = 0

	# For basic assembly stats:
	ref_sequences = set()
	query_sequences = set()
	reference_lengths = []
	query_lengths = {}
	fields_by_query = {}


	for line in f:
		linecounter += 1
		if line[0]==">":
			fields = line.strip().split()
			
			# For delta file output:
			query = scrub(fields[1])
			list_of_unique_alignments = unique_alignments[query]

			header_needed = False
			for index in list_of_unique_alignments:
				if line.strip() == header_lines_by_query[query][index]:
					header_needed = True
			if header_needed == True:
				f_out_delta.write(line) # if we have any alignments under this header, print(the header)
			alignment_counter[query] = alignment_counter.get(query,0)

			# For coords:
			current_reference_name = scrub(fields[0][1:])
			current_query_name = scrub(fields[1])

			current_reference_size = int(fields[2])
			current_query_size = int(fields[3])

			# For index:
			if not current_reference_name in ref_sequences:
				reference_lengths.append((current_reference_name, current_reference_size))
				ref_sequences.add(current_reference_name)
			if not current_query_name in query_sequences:
				query_lengths[current_query_name] = current_query_size
				query_sequences.add(current_query_name)

		else:
			fields = line.strip().split()
			if len(fields) > 4:
				# For coords:
				ref_start = int(fields[0])
				ref_end = int(fields[1])
				query_start = int(fields[2])
				query_end = int(fields[3])
				csv_tag = "repetitive"
				if alignment_counter[query] in list_of_unique_alignments:
					f_out_delta.write(line)
					csv_tag = "unique"
					keep_printing = True
				else:
					keep_printing = False
				fields = [ref_start, ref_end, query_start, query_end, current_reference_size, current_query_size, current_reference_name, current_query_name, csv_tag]
				if fields_by_query.get(current_query_name, None) == None:
					fields_by_query[current_query_name] = []
				fields_by_query[current_query_name].append(fields)
				alignment_counter[query] = alignment_counter[query] + 1

			elif keep_printing == True:
				f_out_delta.write(line)

	f.close()
	f_out_delta.close()
	# f_out_coords.close()

	print("Writing filtered delta file and capturing information for coords file: %d seconds for %d total lines in file" % (time.time()-before,linecounter))
	
	return reference_lengths, fields_by_query

def index_for_dot(reference_lengths, fields_by_query, output_prefix, max_overview_alignments):

	#  Find the order of the reference chromosomes
	reference_lengths.sort(key=lambda x: natural_key(x[0]))
	
	#  Find the cumulative sums
	cumulative_sum = 0
	ref_chrom_offsets = {}
	queries_by_reference = {}
	for ref,ref_length in reference_lengths:
		ref_chrom_offsets[ref] = cumulative_sum
		cumulative_sum += ref_length
		queries_by_reference[ref] = set()

	#  Calculate relative positions of each alignment in this cumulative length, and take the median of these for each query, then sort the queries by those scores
	flip_by_query = {}
	unique_references_by_query = {} # for index, only unique alignments
	all_references_by_query = {} # for index, including repetitive alignments
	relative_ref_position_by_query = [] # for ordering


	ordered_tags = ["unique", "repetitive"]


	f_out_coords = open(output_prefix + ".coords", 'w')
	f_out_coords.write("ref_start,ref_end,query_start,query_end,ref\n")

	query_byte_positions = {}
	query_lengths = {}

	all_alignments = []
	last_query = ""

	for query_name in fields_by_query:

		lines = fields_by_query[query_name]
		sum_forward = 0
		sum_reverse = 0
		ref_position_scores = []
		unique_references_by_query[query_name] = set()
		all_references_by_query[query_name] = set()

		for fields in lines:
			tag = fields[8]

			query_name = fields[7]
			query_lengths[query_name] = int(fields[5])

			all_references_by_query[query_name].add(ref)
			# Only use unique alignments to decide contig orientation
			if tag == "unique":
				query_stop = int(fields[3])
				query_start = int(fields[2])
				ref_start = int(fields[0])
				ref_stop = int(fields[1])
				alignment_length = abs(int(fields[3])-int(fields[2]))
				ref = fields[6]

				# for index:
				unique_references_by_query[query_name].add(ref)
				queries_by_reference[ref].add(query_name)

				# for ordering:
				ref_position_scores.append(ref_chrom_offsets[ref] + (ref_start+ref_stop)/2)

				# for orientation:
				if query_stop < query_start:
					sum_reverse += alignment_length
				else:
					sum_forward += alignment_length

		# orientation:
		flip = sum_reverse > sum_forward
		flip_by_query[query_name] = "-" if (flip == True) else "+"


		for tag in ordered_tags:
			query_byte_positions[(last_query, "end")] = f_out_coords.tell()
			query_byte_positions[(query_name, tag)] = f_out_coords.tell()
			f_out_coords.write("!" + query_name + "!" + tag +"\n")
			
			for fields in lines:
				if fields[8] == tag:
					if flip == True:
						fields[2] = int(fields[5]) - int(fields[2])
						fields[3] = int(fields[5]) - int(fields[3])

					output_fields = [fields[0], fields[1], fields[2], fields[3], fields[6]]
					f_out_coords.write(",".join([str(i) for i in output_fields]) + "\n")
					
					# For alignment overview:
					alignment_length = abs(int(fields[3])-int(fields[2]))
					all_alignments.append(([fields[0], fields[1], fields[2], fields[3], fields[6], fields[7], fields[8]], alignment_length))

		# ordering
		if len(ref_position_scores) > 0:
			relative_ref_position_by_query.append((query_name,np.median(ref_position_scores)))
		else:
			relative_ref_position_by_query.append((query_name,0))

		last_query = query_name


	query_byte_positions[(last_query, "end")] = f_out_coords.tell()

	relative_ref_position_by_query.sort(key=lambda x: x[1])

	f_out_index = open(output_prefix + ".coords.idx", 'w')

	f_out_index.write("#ref\n")
	f_out_index.write("ref,ref_length,matching_queries\n")
	# reference_lengths is sorted by the reference chromosome name
	for ref,ref_length in reference_lengths:
		f_out_index.write("%s,%d,%s\n" % (ref,ref_length,"~".join(queries_by_reference[ref])))

	f_out_index.write("#query\n")
	f_out_index.write("query,query_length,orientation,bytePosition_unique,bytePosition_repetitive,bytePosition_end,unique_matching_refs,matching_refs\n")
	# relative_ref_position_by_query is sorted by rel_pos
	for query,rel_pos in relative_ref_position_by_query:
		f_out_index.write("%s,%d,%s,%d,%d,%d,%s,%s\n" % (query, query_lengths[query], flip_by_query[query], query_byte_positions[(query,"unique")], query_byte_positions[(query,"repetitive")] - query_byte_positions[(query,"unique")], query_byte_positions[(query,"end")] - query_byte_positions[(query,"repetitive")], "~".join(unique_references_by_query[query]), "~".join(all_references_by_query[query])))

	f_out_index.write("#overview\n")
	f_out_index.write("ref_start,ref_end,query_start,query_end,ref,query,tag\n")

	num_overview_alignments = min(max_overview_alignments,len(all_alignments))
	if num_overview_alignments < len(all_alignments):
		print("Included the longest " + str(max_overview_alignments) + " alignments in the index under #overview (change this with the --overview parameter), out of a total of " + str(len(all_alignments)) + " alignments.")

	all_alignments.sort(key=lambda x: -x[1])
	overview_alignments = all_alignments[0:num_overview_alignments]
	for tup in overview_alignments:
		f_out_index.write(",".join([str(i) for i in tup[0]]) + "\n")

	f_out_index.close()

def main():
	parser=argparse.ArgumentParser(description="Take a delta file, apply Assemblytics unique anchor filtering, and prepare coordinates input files for Dot")
	parser.add_argument("--delta",help="delta file" ,dest="delta", type=str, required=True)
	parser.add_argument("--out",help="output file" ,dest="out", type=str, default="output")
	parser.add_argument("--unique-length",help="The total length of unique sequence an alignment must have on the query side to be retained. Default: 10000" ,dest="unique_length",type=int, default=10000)
	parser.add_argument("--overview",help="The number of alignments to include in the coords.idx output file, which will be shown in the overview for Dot. Default: 1000" ,dest="overview",type=int, default=1000)
	parser.set_defaults(func=run)
	args=parser.parse_args()
	args.func(args)

if __name__=="__main__":
	main()
```
