## Supplemental File 1 for "Long-Read Genome Assembly and Gene Model Annotations for the Rodent Malaria Parasite *Plasmodium yoelii* 17XNL": dotter.sh

```
set -uex

REF=$1
QUERY=$2
PREFIX=$3

# Nucmer: minimum length of a single match
L=100

# Nucmer: minimum length of a cluster of matches
C=500

# Nucmer parameters.
PARAMS="--maxmatch -l 100 -c 500"

# Run nucmer.
nucmer ${PARAMS} ${REF} ${QUERY} --prefix ${PREFIX}

# Convert the delta file.
python scripts/misc/DotPrep.py --delta  ${PREFIX}.delta --out  ${PREFIX}

# Cleanup.
rm -rf *.delta.gz
```
