## Supplemental File 1 for "Long-Read Genome Assembly and Gene Model Annotations for the Rodent Malaria Parasite *Plasmodium yoelii* 17XNL": format_gff.py

```
import csv, sys

"""
Adds locus tag and product tags
Modifies transcript_id, protein_id to the format gnl|dbname|id
Input gff3 file needs to be sorted by chromosome.
"""


def extract_attr(attr_str, attr_tag):
    attrs = attr_str.split(";")
    attrs = [a.strip() for a in attrs]  # remove all spaces
    try:
        feat_id = [val for val in attrs if val.startswith(attr_tag)][0]
    except:
        print(f'{attr_tag} not found for the feature in annotation file')
        print(f'{attr_str}')
        sys.exit()
    feat_id = feat_id.replace(attr_tag + "=", "")  # for gff
    feat_id = feat_id.replace(attr_tag + " ", "")  # for gtf
    feat_id = feat_id.replace('"', "")  # remove all quotes.
    # feat_id = feat_id.split(",")  # for cases like Parent=mRNA1,mRNA2,mRNA3 in gff3
    return feat_id


def modify_gene_attr(attr, locus_tag):
    new_attr = attr.strip() + f";locus_tag={locus_tag}"
    return new_attr


def modify_attr(attr, tid, locus_tag):
    # tid2 = f'gnl|psuPy17xnl|{tid}'
    tid2 = tid
    new_attr = attr.strip() + f";locus_tag={locus_tag}"
    curr_tid_tag = f'transcript_id={tid}'
    new_tid = f'transcript_id={tid2}'
    new_attr = new_attr.replace(curr_tid_tag, new_tid)

    return new_attr


def modify_mRNA_CDS_attr(feat, attr, tid, locus_tag, product):
    t, no = tid.split(".")

    pid = tid + f"-p{no}"
    # pid2 = f'gnl|PSU_Py17XNL|{pid}'
    # tid2 = f'gnl|PSU_Py17XNL|{tid}'
    pid2 = pid
    tid2 = tid

    new_attr = attr.strip() + f";locus_tag={locus_tag};protein_id={pid2};product={product}"
    curr_tid_tag = f'transcript_id={tid}'
    new_tid = f'transcript_id={tid2}'

    new_attr = new_attr.replace(curr_tid_tag, new_tid)

    if feat == "CDS":
        cds_name = extract_attr(attr, "Name")
        name_tag = f'Name={cds_name}'
        new_name_tag = f'Name={pid}'
        new_attr = new_attr.replace(name_tag, new_name_tag)

    return new_attr


def read_desc(fname):
    store = dict()

    stream = csv.DictReader(open(fname), delimiter="\t")

    for row in stream:
        id, d = row['protein_id'], row['description']
        store[id] = d
    return store


def read_chroms(fname):
    store = dict()
    stream = csv.reader(open(fname), delimiter="\t")

    for row in stream:
        chrom, size = row[0], row[1]
        store[chrom] = size
    return store


def add_region(chrom, size):
    start = 1
    end = size
    cno = chrom.replace("Py17XNL_", "")

    if cno not in ('API', 'MIT'):
        genome = "chromosome"
    elif cno == "MIT":
        genome = "MIT"
    else:
        genome = "apicoplast"

    rid = chrom if "unloc" in cno else f"{chrom}_chr{cno}"

    chrom_attr = f'ID={rid};Name={chrom};genome={genome};mol_type=genomic DNA'
    out = [chrom, "AUGUSTUS", "region", str(start), str(end), ".", "+", ".", chrom_attr]
    rline = "\t".join(out)
    rheader = f"##sequence-region {chrom} 1 {end}"
    return rheader, rline


def reformat(fname, desc, sizes):
    stream = csv.reader(open(fname), delimiter="\t")
    # stream = islice(stream, LIMIT)
    match = "Unknown"
    locus_tag, tid, new_attr = "", "", ""
    prev_chr = ""

    for row in stream:
        if row[0].startswith("#"):
            print("\t".join(row))
            continue

        chrom, feat, attr = row[0], row[2], row[8]

        if feat == "five_prime_utr":
            row[2] = "five_prime_UTR"

        if feat == "three_prime_utr":
            row[2] = "three_prime_UTR"

        if chrom != prev_chr:
            clen = sizes[chrom]
            rheader, rline = add_region(chrom, clen)
            print(rheader)
            print(rline)

        prev_chr = chrom

        if feat == "gene":
            gid = extract_attr(attr, "gene_id")
            locus_tag = gid
            new_attr = modify_gene_attr(attr, locus_tag)

        elif feat == "mRNA" or feat == "exon" or feat == "CDS":
            tid = extract_attr(attr, "transcript_id")
            if feat == "mRNA":
                match = extract_attr(attr, "best_match")
            product = desc.get(match, "Unknown")
            new_attr = modify_mRNA_CDS_attr(feat, attr, tid, locus_tag, product)

        else:
            new_attr = modify_attr(attr, tid, locus_tag)

        row[8] = new_attr
        print("\t".join(row))


if __name__ == "__main__":
    #fname = "17XNL_PSU_mod1.gff3"
    fname = sys.argv[1]
    desc_f = sys.argv[2] #"accession_map_PY17X_desc.txt"
    chroms_f = sys.argv[3] #"chroms.txt"
    desc = read_desc(desc_f)
    chroms = read_chroms(chroms_f)
    reformat(fname, desc, chroms)
```
