## Supplemental File 1 for "Long-Read Genome Assembly and Gene Model Annotations for the Rodent Malaria Parasite *Plasmodium yoelii* 17XNL": make_var_table.py

```
import csv,sys


def store_data(gnames):
    store = dict()
    stream = csv.reader(open(gnames, 'r'), delimiter='\t')

    for row in stream:
        gid = row[4]
        store[gid] = row

    return store


def combine(cds, gnames):
    store = store_data(gnames)

    stream2 = csv.reader(open(cds, 'r'), delimiter='\t')

    for row in stream2:
        gid = row[7]
        if gid in store:
            data = store[gid]
            row.extend(data)
            print('\t'.join(row))


if __name__ == "__main__":
    cds = sys.argv[1] #"17XNL_PSU_cds_ann_short.txt"
    gnames = sys.argv[2] #"out.txt"

    combine(cds, gnames)
```
