## Supplemental File 1 for "Long-Read Genome Assembly and Gene Model Annotations for the Rodent Malaria Parasite *Plasmodium yoelii* 17XNL": replace_header.py

```
import sys, csv,re
from itertools import groupby
from collections import defaultdict


def read_map(mapfile):
    store = defaultdict(list)
    stream = csv.DictReader(open(mapfile), delimiter="\t")

    for row in stream:
        org_id = row['org_id']
        new_id = row['new_id']
        store[org_id] = new_id
    return store


def is_header(line):
    return line.startswith(">")


def rename_header(fname, store):
    fa = open(fname, "r")

    for header, seq in groupby(fa, key=is_header):
        if header:
            hline = next(seq).strip()
            hline = hline.lstrip(">")
            prefix, suffix = hline.split(".")
            new_prefix = store[prefix] if prefix in store else ""
            #newh = f"{new_prefix}.{suffix}"
            newh = re.sub(prefix, new_prefix, hline)
            newh = ">" + newh #+ f' ID={hline}'
        else:
            seq = "".join([line.strip() for line in seq])
            seq = seq.upper()
            # split seq into 70 characters each line
            seq1 = "\n".join([seq[i:i + 60] for i in range(0, len(seq), 60)])
            yield (newh, seq1)


if __name__ == "__main__":

    fname = sys.argv[1] #"transcripts.fa"
    mapfile = sys.argv[2] # "transcript_ann.txt"

    id_store = read_map(mapfile)
    fa_store = dict(rename_header(fname, id_store))
    for header, seq in fa_store.items():
        print(header)
        print(seq)
```
