## Supplemental File 1 for "Long-Read Genome Assembly and Gene Model Annotations for the Rodent Malaria Parasite *Plasmodium yoelii* 17XNL": vcf2table.py

```
"""
Create a table out of snpEff annotated VCF file based on first annotations.
"""
import csv, sys


def make_table(vcf):
    stream = csv.reader(open(vcf), delimiter="\t")

    header = "\t".join(
        ["chrom", "pos", "ref", "alt", "effect", "impact", "gene_name", "gene_id", "feat", "feat_id", "biotype",
         "aa_change"])
    print(header)
    for data in stream:
        if data[0].startswith("#"):
            continue
        chrom, pos, ref, alt, ann = data[0], data[1], data[3], data[4], data[7]

        try:
            first_ann = ann.split("ANN=")[1].split(",")[0]
        except:
            prot, impact, gene_name, gene_id, feat, feat_id, biotype = "","","","","","",""
            effect = "either upstream or downstream variant"
            out = "\t".join([chrom, pos, ref, alt, effect, impact, gene_name, gene_id, feat, feat_id, biotype, prot])
            print(out)
            continue

        # print(first_ann)
        ann_vals = first_ann.split("|")
        effect, impact, gene_name, gene_id, feat, feat_id, biotype = ann_vals[1:8]
        prot = ann_vals[10]
        out = "\t".join([chrom, pos, ref, alt, effect, impact, gene_name, gene_id, feat, feat_id, biotype, prot])
        print(out)


if __name__ == "__main__":
    vcf_file = sys.argv[1]  # "17X_PLD_cds_ann_short.vcf"
    make_table(vcf_file)
```
