## Supplemental File 1 for "Long-Read Genome Assembly and Gene Model Annotations for the Rodent Malaria Parasite *Plasmodium yoelii* 17XNL": bed.py

```
"""
Convert tRNASCAN-SE output to a bed file.
"""
import sys, csv

fname = sys.argv[1]

stream = csv.reader(open(fname))
next(stream)
next(stream)
next(stream)
next(stream)
next(stream)
for row in stream:
    line = row[0].split()
    chrom, start, end, desc = line[0], line[2], line[3], line[4]
    chrom = chrom.strip()
    start = int(start)  # - 1
    end = int(end)
    strand = "+"
    if start > end:
        start, end = end-1, start
        strand = "-"
    start = start -1 if strand == "+"  else start
    start = str(start).strip()
    end = str(end).strip()
    name = f"tRNA {desc}"

    print("\t".join([chrom, start, end, name, ".", strand]))
```
