## Supplemental File 1 for "Long-Read Genome Assembly and Gene Model Annotations for the Rodent Malaria Parasite *Plasmodium yoelii* 17XNL": best.py

```
import csv, sys

# Extract best hit from blast output
# Best hits satisfy the following conditions
# 1. Both query and hit are in the same chromosome
# 2. pident= 100 and query_length=hit_length


def parse(fname):
    stream = csv.reader(open(fname), delimiter="\t")
    header = next(stream)

    found = set()

    for row in stream:
        query, hit, pident, length, mismatch, gapopen, qstart, qend, sstart, send, sstrand = row

        if query in found:
            continue

        chrm = hit.replace('Py17XNL', 'Py17X')

        pident = int(float(pident))
        length = int(length)
        qstart = int(qstart)
        qend = int(qend)
        sstart = int(sstart)
        send = int(send)
        if query.startswith(chrm):
            if int(pident) == 100 and abs(send - sstart) + 1 == length:
                print("\t".join(row))
                found.add(query)

if __name__ == "__main__":
    fname = sys.argv[1] # "17XNL_PSU_all_rRNA_blastn.txt"
    parse(fname)
```
