## Supplemental File 1 for "Long-Read Genome Assembly and Gene Model Annotations for the Rodent Malaria Parasite *Plasmodium yoelii* 17XNL": get_last_gid.py

```
import csv, os, sys
import gffutils

"""
This script generate a text file with the last unique geneid per chromosome.
"""


def store_gff3(fname, dbname):
    """
    Stores the gff3 file in a database and returns the database handle.
    """

    if not (os.path.exists(dbname)):
        # Create an sqlite3 database. This is a one time operation.
        db = gffutils.create_db(fname, dbname)

    db = gffutils.FeatureDB(dbname, keep_order=True)

    return db


def get_genes(gff, dbname):
    """
    Extract all gene names from a gff file
    """

    store = list()
    dbname = dbname

    if not (os.path.exists(dbname)):
        # Create an sqlite3 database. This is a one time operation.
        db = gffutils.create_db(gff, dbname)

    db = gffutils.FeatureDB(dbname, keep_order=True)
    # Print all genes
    for g in db.all_features(featuretype='gene'):
        # gid = g.id
        # name = g.attributes['Name'][0]
        # store[name] = gid
        store.append(g.id)
    return store


def get_last_geneid(ann, dbname):
    """
    Extract last geneid for each chromsome
    Returns a dictionary with chromosome as key and last gid as the value.
    """
    last = dict()
    genes = sorted(get_genes(ann, dbname))
    for g in genes:
        chrom, gnum = g.split("_")
        gnum = str(gnum)
        cnum = gnum[:2] if gnum[0].isdigit() else gnum[:3]
        chrom_id = f"{chrom}_{cnum}"
        last[chrom_id] = g
    return last


if __name__ == "__main__":
    ann =  sys.argv[1] #"hybrid_new.gff3"
    dbname = "ann.db"

    dbh = store_gff3(ann, dbname)
    last_gids = get_last_geneid(ann, dbname)

    header = "\t".join(['chrom', 'gid'])
    print(header)

    for k, v in last_gids.items():
        out = "\t".join([k, v])
        print(out)
```
