## Supplemental File 1 for "Long-Read Genome Assembly and Gene Model Annotations for the Rodent Malaria Parasite *Plasmodium yoelii* 17XNL": get_rRNAs.sh

```
# 
# This scripts identifies 17XNL rRNAs based on blast searches using 17X rRNA sequences.
#

set -uex

# Reference genome file
REF_GENOME=$1  # refs/17X_PLD.fa

#Reference annotation file
REF_GFF=$2 # refs/17X_PLD.gff

# Genome assembly fasta
ASM=$3 #asm/17XNL_PSU.fa

# Output folder
OUT=$4 #ann/ncRNA/rRNA

SRC=scripts/ncRNA

ASM_PREFIX=$(basename ${ASM%.fa})
echo $ASM_PREFIX

REF_PREFIX=$(basename ${REF_GENOME%.fa})
echo $REF_PREFIX

# Create output directory
mkdir -p ${OUT}

# Get 17X rRNA coordinates.
cat ${REF_GFF} | awk 'BEGIN{OFS="\t"} $3=="rRNA" {print $1,$4,$5, ".",$7}' >${OUT}/${REF_PREFIX}_rRNA.txt

# Get 17X rRNA sequences in the plus strand.
cat ${OUT}/${REF_PREFIX}_rRNA.txt | awk '$5=="+" {print}' |parallel --colsep "\t" samtools faidx ${REF_GENOME} {1}:{2}-{3} >${OUT}/rRNA_plus.fa

sed -i 's/:/_/' ${OUT}/rRNA_plus.fa

# Get sequences in rev strand and reverse complement it.
cat ${OUT}/${REF_PREFIX}_rRNA.txt | awk '$5=="-" {print}' |parallel --colsep "\t" samtools faidx ${REF_GENOME} {1}:{2}-{3} >${OUT}/rRNA_minus.fa

sed -i 's/:/_/' ${OUT}/rRNA_minus.fa
revseq -sequence ${OUT}/rRNA_minus.fa -outseq ${OUT}/rRNA_minus_rev.fa

# Concatenate all
cat ${OUT}/rRNA_plus.fa ${OUT}/rRNA_minus_rev.fa >${OUT}/all_rRNA.fa

# Make blast database out 17XNL_PSU assembly.
mkdir -p ${OUT}/db
makeblastdb -in ${ASM} -dbtype nucl -out ${OUT}/db/${ASM_PREFIX}.fa

# Run blast
echo -e "qaccver\tsaccver\tpident\tlength\tmismatch\tgapopen\tqstart\tqend\tsstart\tsend\tsstrand" >${OUT}/${ASM_PREFIX}_all_rRNA_blastn.txt
blastn -task blastn -query ${OUT}/all_rRNA.fa -db ${OUT}/db/${ASM_PREFIX}.fa -outfmt '6 qaccver saccver pident length mismatch gapopen qstart qend sstart send sstrand' >>${OUT}/${ASM_PREFIX}_all_rRNA_blastn.txt

# Parse blast to extract best hits

python ${SRC}/best.py ${OUT}/${ASM_PREFIX}_all_rRNA_blastn.txt | awk 'BEGIN{OFS="\t"}{print $2,$9,$10,$1,".",$11}' | sed -e 's/plus/+/' -e 's/minus/-/' >${OUT}/${ASM_PREFIX}_all_rRNA_blastn_best.txt

# Create a bed file
cat ${OUT}/${ASM_PREFIX}_all_rRNA_blastn_best.txt| awk 'BEGIN{OFS="\t"} { if($6=="+" ){print $1,$2-1,$3,$4,$5,$6} else {print $1,$3-1,$2,$4,$5,$6}}' > ${OUT}/${ASM_PREFIX}_rRNA.bed


# Additional notes.
# Manually modify the bed file to include rRNA names in the bed file.
# Manually added the best hit for Py17X_MIT_v3_3429-3544 as it is only 98% identical. 
# Corrected 5S best hits
# The resulting is final file is 17XNL_PSU_rRNA.bed 
```
