## Supplemental File 1 for "Long-Read Genome Assembly and Gene Model Annotations for the Rodent Malaria Parasite *Plasmodium yoelii* 17XNL": get_tRNAs.sh

```
#
# This script make tRNA gene predictions and creates a bed file for it
# 
#  Assembly fasta and output folder are needed as input
#
# tRNAs are predicted with tRNASCAN_SE

ASM=$1  #results/17XNL_PSU.fa
OUT=$2 # outdir ann/tRNA

SCRIPTS=scripts/ncRNA

# Create outdir
mkdir -p ${OUT}

PREFIX=$(basename ${ASM%.fa})
echo $PREFIX

tRNA_OUT=${OUT}/${PREFIX}"_tRNA.txt"
tRNA_BED=${OUT}/${PREFIX}"_tRNA.bed"

# Predict tRNAs
echo "Making tRNA predictions."
tRNAscan-SE -E $ASM >$tRNA_OUT

# Make a bed file
echo "Making tRNA bed file."
python ${SCRIPTS}/bed.py $tRNA_OUT  >${tRNA_BED}
```
