## Supplemental File 1 for "Long-Read Genome Assembly and Gene Model Annotations for the Rodent Malaria Parasite *Plasmodium yoelii* 17XNL": ncRNA_gff.py

```
"""

Convert bed to gff3 format.
This script was used to create ncRNA gff file.

The geneid of the last gene in each chromosome is to be inputted along with a bed file of ncRNAs.
Geneid file is needed to assign the (next) geneid to the newly added genes

"""
import sys, csv, re


def parse(fname):
    """
    Parses the tRNASCAN output
    """
    stream = csv.reader(open(fname), delimiter="\t")
    for line in stream:
        chrom, start, end, desc, score, strand = line
        chrom = chrom.strip()
        start = int(start)  # - 1
        end = int(end)
        #strand = "+"
        if start > end:
            start, end = end, start
            strand = "-"
        start = start + 1 
        start = str(start).strip()
        end = str(end).strip()
        name = f"{desc}"

        yield chrom, start, end, name, strand


def read_gids(gid_file):
    store = dict()

    data = csv.reader(open(gid_file), delimiter="\t")

    for row in data:
        chrom, gid = row
        store[chrom] = gid
    return store


def make_gene_row(chrom, start, end, name, strand, gid, source, feat):
    # source = "tRNASCAN-SE"
    # feat = "ncRNA_gene"
    # feat = "ncRNA"

    gattr = f"ID={gid};gene_id={gid};gene_name={gid};Name={name};best_match={source};description={name};locus_tag={gid}"
    out = "\t".join([chrom, source, feat, start, end, ".", strand, ".", gattr])
    return out


def make_ncRNA_row(chrom, start, end, name, strand, gid, source, feat):
    # source = "tRNASCAN-SE"
    # feat = "tRNA"
    trna_id = gid + ".1"
    parent = gid
    #tattr = f"ID={trna_id};Parent={parent};description={name};best_match={source};locus_tag={gid};product={name}"
    tattr = f"ID={trna_id};Parent={parent};description={name};best_match={source};locus_tag={gid}"
    out = "\t".join([chrom, source, feat, start, end, ".", strand, ".", tattr])
    return out


def make_exon_row(chrom, start, end, name, strand, gid, source, feat):
    # source = "tRNASCAN-SE"
    # feat = "exon"
    exon_id = gid + ".1-exon1"
    parent = gid + ".1"
    #eattr = f"ID={exon_id};Name={exon_id};Parent={parent};gene_name={gid};best_match={source};locus_tag={gid};product={name}"
    eattr = f"ID={exon_id};Name={exon_id};Parent={parent};gene_name={gid};best_match={source};locus_tag={gid}"
    out = "\t".join([chrom, source, feat, start, end, ".", strand, ".", eattr])
    return out


def make_gff(bed_file, gid_file):
    # Read gids file
    gids = read_gids(gid_file)
    items = ""

    # Read bed file and parse.
    for res in parse(bed_file):
        chrom, start, end, name, strand = res

        name = name.replace(' ', '_')

        # Get gid and increment it
        gid = gids[chrom]

        # Split between non-digit and digits
        match = re.match(r"([a-zA-z]+.*[a-zA-z])([0-9]+)", gid, re.I)
        if match:
            items = match.groups()

        cid, gid_idx = items
        gid_idx = int(gid_idx) + 1
        gid_idx = str(gid_idx)
        if cid not in ("Py17XNL_MIT", "Py17XNL_API"):
            gid_idx = gid_idx.zfill(7)
        else:
            gid_idx = gid_idx.zfill(5)
        
        new_gid = "".join([cid, str(gid_idx)])

        gids[chrom] = new_gid

        if "rRNA" in name  or "ribosomal" in name:
            feat = "rRNA"
            # source = "RNAmmer"
            source = "blastn"
        else:
            feat = "tRNA"
            source = "tRNASCAN-SE"

        gene_row = make_gene_row(chrom, start, end, name, strand, new_gid, source, feat="ncRNA_gene")
        ncRNA_row = make_ncRNA_row(chrom, start, end, name, strand, new_gid, source, feat)
        exon_row = make_exon_row(chrom, start, end, name, strand, new_gid, source, feat="exon")

        print(gene_row)
        print(ncRNA_row)
        print(exon_row)


if __name__ == "__main__":
    bed_fname = sys.argv[1]  # "17XNL_PSU_tRNA.bed"
    gid_fname = sys.argv[2]  # "gids.txt"
    make_gff(bed_fname, gid_fname)
```
