## Supplemental File 1 for "Long-Read Genome Assembly and Gene Model Annotations for the Rodent Malaria Parasite *Plasmodium yoelii* 17XNL": flye.sh

```
#!/usr/bin/env bash
set -uex

# Requirements and installation
# conda install flye
# git clone https://github.com/rrwick/Porechop.git
# cd Porechop
# python3 setup.py install


READS=../reads/nano_dna.fq.gz

REF=../refs/PlasmoDB-52_Pyoeliiyoelii17X_Genome.fasta
OUT=flye_py_v2.9

mkdir -p $OUT

TRIMMED=${OUT}/trimmed.fq

# Remove adapters with Porechop
porechop -i $READS -o $TRIMMED

# Get stats on before and after trimming.
#seqkit stats $READS $TRIMMED >$OUT/stats.txt

#Flye assembler
flye --nano-corr $TRIMMED --out-dir $OUT --genome-size 23m -t 20

#Optional: Polishing with Flye
# flye --polish-target  ${OUT}/assembly.fasta --nano-raw $TRIMMED -t 20 -o ${OUT}/polished

# Scaffold
ragtag.py scaffold ${REF} ${OUT}/assembly.fasta -o ${OUT}/scaffold
samtools faidx ${OUT}/scaffold/ragtag.scaffold.fasta


# Copy final assembly.
cp ${OUT}/assembly.fasta  .

# Copy final scaffold
cp ${OUT}/scaffold/ragtag.scaffold.fasta* .

#Index fasta files
ls *.fasta| parallel samtools faidx {}

#Generate quast report
quast.py -o quast_asm  -r $REF *fa *fasta

# Clean
# rm -rf $OUT
#rm -f list
```
