## Supplemental File 1 for "Long-Read Genome Assembly and Gene Model Annotations for the Rodent Malaria Parasite *Plasmodium yoelii* 17XNL": map_ont.sh

```
#!/usr/bin/env bash
set -uex

#
# Map different datasets to assembly
#

REF=$1
OUTDIR="bam" #"asm_map"

PY17X_GENOME=../refs/PlasmoDB-52_Pyoeliiyoelii17X_Genome.fasta
rm -rf $OUTDIR

mkdir -p $OUTDIR
cd $OUTDIR
ASM="Py17XNL_assembly.fa"
cp ../$REF $ASM

CPU=10


# Reads
CONTIG1=../reads/17XNL_1P.fq.gz
CONTIG2=../reads/17XNL_2P.fq.gz
CONTIG_WT=../reads/WT.fq.gz

CONTIG_NANO_DNA=../reads/nano_dna.fq.gz
CONTIG_NANO_RNA=../reads/nano_rna.fq.gz

TRANSCRIPT_X_DNA=../refs/PlasmoDB-52_Pyoeliiyoelii17X_AnnotatedTranscripts.fasta
TRANSCRIPT_XNL_DNA=../refs/PlasmoDB-52_Pyoeliiyoelii17XNL_AnnotatedTranscripts.fasta

mkdir -p index
INDEX=index/py17xnl_assm

illumina_dna_bam=$ASM".illumina.dna.bam"
illumina_rna_bam=$ASM".illumina.rna.bam"
nano_dna_bam=$ASM".nano.dna.bam"
nano_rna_bam=$ASM".nano.rna.bam"
transcript_X_bam=$ASM".trans.X.bam"
transcript_XNL_bam=$ASM".trans.XNL.bam"

VCF=$ASM".fb.vcf"
FILT=$ASM".fb.filt.vcf.gz"


# Make read mappings

bwa index -p $INDEX $ASM  >> log.txt  2>&1
hisat2-build $ASM $INDEX >> log.txt 2>&1

# Illumina
bwa mem -t $CPU $INDEX $CONTIG1 $CONTIG2 2>> log.txt | samtools sort > ${illumina_dna_bam} 2>> log.txt
hisat2 -p $CPU --max-intronlen 1500 -x $INDEX -U $CONTIG_WT 2>> log.txt | samtools sort > ${illumina_rna_bam}  2>> log.txt

# Nanopore
minimap2 -t $CPU -a -x map-ont $ASM $CONTIG_NANO_DNA 2>> log.txt | samtools sort > ${nano_dna_bam} 2>> log.txt
minimap2 -t $CPU -a -x splice $ASM -G 3000 $CONTIG_NANO_RNA 2>> log.txt | samtools sort > ${nano_rna_bam} 2>> log.txt

# Transcripts
minimap2 -t $CPU -a -x splice $ASM $TRANSCRIPT_X_DNA   2>> log.txt | samtools sort > ${transcript_X_bam} 2>> log.txt
minimap2 -t $CPU -a -x splice $ASM $TRANSCRIPT_XNL_DNA 2>> log.txt | samtools sort > ${transcript_XNL_bam}  2>> log.txt

# Call variants with freebayes.
freebayes -f $ASM ${illumina_dna_bam} -C 5 -p 1 |  bcftools norm --threads $CPU -f $ASM -o $VCF
bgzip $VCF
# Filter variants
cat $VCF".gz" |bcftools filter -Oz --IndelGap 5 -i  'AF!=0 && GT=="1"' >$FILT
bcftools index $FILT

# Map Py17X to assembly
minimap2 -ax asm5 $ASM $PY17X_GENOME |samtools sort >Py17X.bam
samtools index Py17X.bam

# Index and cleanup
ls -1 *.bam | parallel samtools index {}
rm -f *.ht2 *.ann *.bwt *.amb
```
