## Supplemental File 1 for "Long-Read Genome Assembly and Gene Model Annotations for the Rodent Malaria Parasite *Plasmodium yoelii* 17XNL": multi_polish.py

```
import sys

HAS_DIRNAME = len(sys.argv) > 1

if HAS_DIRNAME:
    dirname = sys.argv[1]
else:
    dirname = "polish"

# Evaluation mode.
EVAL_MODE = not HAS_DIRNAME

USE_FREEBAYES = True

print(f"# DIRNAME={dirname}")
print(f"# EVAL_MODE={EVAL_MODE}")

CPU = 20
#REF = '../asm/Pyoelii_consensus_00.fa'
REF = '../asm/scaffold.fasta'
CONTIG1 = '../reads/17XNL_1P.fq.gz'
CONTIG2 = '../reads/17XNL_2P.fq.gz'
CONTIG_WT = '../reads/WT.fq.gz'

CONTIG_NANO_DNA = '../reads/nano_dna.fq.gz'
CONTIG_NANO_RNA = '../reads/nano_rna.fq.gz'

TRANSCRIPT_X_DNA = '../refs/PlasmoDB-52_Pyoeliiyoelii17X_AnnotatedTranscripts.fasta'
TRANSCRIPT_XNL_DNA = '../refs/PlasmoDB-52_Pyoeliiyoelii17XNL_AnnotatedTranscripts.fasta'

GENOME = '../refs/PlasmoDB-52_Pyoeliiyoelii17X_Genome.fasta'

#
# Override for evaluation mode
#
if EVAL_MODE:
    CPU = 4
    REF = '../asm/contig.fa'
    CONTIG1 = '../reads/contig_1.fq.gz'
    CONTIG2 = '../reads/contig_2.fq.gz'

    CONTIG_WT = '../reads/contig_wt.fq.gz'

    CONTIG_NANO_DNA = '../reads/contig_nano_dna.fq.gz'
    CONTIG_NANO_RNA = '../reads/contig_nano_rna.fq.gz'

# Read simulation parameters.
ERR = 0
N = 1000000
LEN = 150
R1 = 'read1.fq'
R2 = 'read2.fq'

# Nanopolish output
POLISH1 = 'step_2.fa'
# Homopolymer corrected output
POLISH2 = 'step_3.fa'
# Final correction.
POLISH3 = 'step_4.fa'

ALL1 = f"{POLISH1}.all.vcf.gz"
FILT1 = f"{POLISH1}.fb.sim.filt.vcf.gz"
# FILT1 = f"{POLISH1}.filt.vcf.gz"


setup = f"""
# Initialization

set -uex

rm -rf {dirname}
mkdir -p {dirname}
mkdir -p {dirname}/index
mkdir -p {dirname}/nextpolish
mkdir -p {dirname}/homopolymer_correction
cd {dirname}
cat {REF} > step_1.fa
echo > log.txt
"""

print(setup)

# Polish1 : Polish with nextPolish.
cmd_nextPolish = f"""
# *** Polish assmebly with nextPolish
cd nextpolish
ls ../{CONTIG1} ../{CONTIG2} >sgs.fofn
genome=../step_1.fa
echo -e "task = best\nworkdir=./rundir\ngenome = $genome\nsgs_fofn = sgs.fofn" > run.cfg
nextPolish run.cfg
cat ./rundir/genome.nextpolish.fasta >../{POLISH1}
samtools faidx ../{POLISH1}
cd ../
        """

print(cmd_nextPolish)

index1 = f"index/{POLISH1}"
bam_sim = f"{POLISH1}.sim.dna.bam"

# Polish2 : Homopolymer correction
cmd_correct = f"""
# *** Correct homopolymer erros.

cd homopolymer_correction
# simulate error-free reads.
pywgsim -S 0 -r 0 -R 0 -N {N} -e {ERR} -1 {LEN} -2 {LEN} ../{GENOME} {R1} {R2} > log.txt 2>&1

# Build index
bwa index -p ../{index1} ../{POLISH1} >> log.txt  2>&1

# Align the simulated reads to assembly
bwa mem -t {CPU} ../{index1} {R1} {R2} 2>> log.txt | samtools sort > {bam_sim} 2>> log.txt
samtools index {bam_sim}

# Call variants with bcftools
#bcftools mpileup --threads {CPU} -Ou -f ../{POLISH1} {bam_sim} 2>> log.txt | bcftools call  --threads {CPU} --ploidy 1 \
# -mv | bcftools norm --threads {CPU} -f ../{POLISH1} -o {ALL1}

# Call variants with freebayes.
freebayes -f ../{POLISH1} {bam_sim} -C 5 -p 1 | bcftools norm --threads {CPU} -f ../{POLISH1} -o {POLISH1}.fb.sim.vcf
bgzip {POLISH1}.fb.sim.vcf
bcftools index {POLISH1}.fb.sim.vcf.gz

# Filter long indels
cat {POLISH1}.fb.sim.vcf.gz |bcftools filter -Oz --IndelGap 5 -i 'TYPE!~"snp" && abs(strlen(ALT)-strlen(REF))>=5  \
&& FORMAT/AO >5 && GT=="1" ' >{FILT1}
bcftools index  {FILT1}


# Filter long indels
#cat {ALL1} |bcftools filter -Oz --IndelGap 5 -i 'TYPE!~"snp" && abs(strlen(ALT)-strlen(REF))>=5  && IDV >5 && \
#GT=="1" ' >{FILT1}
#bcftools index  {FILT1}

# Consensus call, creating new build
cat ../{POLISH1} | bcftools consensus -H 1 {FILT1} > {POLISH2}
cp {POLISH2} ../
#cp {POLISH1}.fb.vcf.gz* ../
#cp {FILT1}* ../
cd ../
"""
print(cmd_correct)

# Polish3 : Correction with illumina reads.

STEPS = 4

for x in range(3, STEPS + 1):

    ref = f"step_{x}.fa"
    bam_dna = f"{ref}.illumina.dna.bam"
    bam_rna = f"{ref}.illumina.rna.bam"
    all = f"{ref}.vars.all.vcf.gz"
    vcf = f"{ref}.vars.trusted.vcf.gz"
    index = f"index/{ref}"
    cmd1 = f'''

#
#
# *** alignment step {x}
#

bwa index -p {index} {ref}  >> log.txt  2>&1
hisat2-build {ref} {index} >> log.txt 2>&1

# Illumina
bwa mem -t {CPU} {index} {CONTIG1} {CONTIG2} 2>> log.txt | samtools sort > {bam_dna} 2>> log.txt
hisat2 -p {CPU} --max-intronlen 1500 -x {index} -U {CONTIG_WT} 2>> log.txt | samtools sort > {bam_rna}  2>> log.txt

# Nanopore
minimap2 -t {CPU} -a -x map-ont {ref} {CONTIG_NANO_DNA} 2>> log.txt | samtools sort > {ref}.nano.dna.bam 2>> log.txt
minimap2 -t {CPU} -a -x map-ont {ref} {CONTIG_NANO_RNA} 2>> log.txt | samtools sort > {ref}.nano.rna.bam 2>> log.txt

# Transcripts
minimap2 -t {CPU} -a -x map-ont {ref} {TRANSCRIPT_X_DNA}   2>> log.txt | samtools sort > {ref}.trans.X.bam 2>> log.txt
minimap2 -t {CPU} -a -x map-ont {ref} {TRANSCRIPT_XNL_DNA} 2>> log.txt | samtools sort > {ref}.trans.XNL.bam  2>> log.txt

# Call variants
#bcftools mpileup --threads {CPU} -Ou -f {ref} {bam_dna} 2>> log.txt | bcftools call  --threads {CPU} --ploidy 1  -mv | bcftools norm --threads {CPU} -f {ref} -o {all}
#cat {all} | bcftools filter -Oz --IndelGap 5 -i 'QUAL>40 && DP>5 && GT=="1"' > {vcf}
#bcftools index {vcf}

# Call variants with freebayes.
freebayes -f {ref} {bam_dna} -C 5 -p 1 |  bcftools norm --threads {CPU} -f {ref} -o {ref}.fb.vcf 
bgzip {ref}.fb.vcf
# Filter variants
cat {ref}.fb.vcf.gz |bcftools filter -Oz --IndelGap 5 -i  'AF!=0 && GT=="1"' >{ref}.fb.filt.vcf.gz
bcftools index {ref}.fb.filt.vcf.gz


# Index and cleanup
ls -1 *.bam | parallel samtools index {{}}
rm -f *.ht2 *.ann *.bwt *.amb
    '''
    print(cmd1)

    if x != STEPS:
        seqid = f"{x + 1}"
        new = f"step_{seqid}.fa"

        cmd_freebayes = f'''
        # *** consensus call, creating build {seqid}
        cat {ref} | bcftools consensus -H 1 {ref}.fb.filt.vcf.gz > {new}
        '''

        cmd_bcftools = f'''
    # *** consensus call, creating build {seqid}
    cat {ref} | bcftools consensus -H 1 {vcf} > {new}
            '''

        if USE_FREEBAYES:
            print(cmd_freebayes)
        else:
            print(cmd_bcftools)

# A few additional mappings  and variant calls to produce.
pairs = [
    ("../refs/PlasmoDB-52_Pyoeliiyoelii17X_Genome.fasta", "py17X.fa"),
    ("../refs/PlasmoDB-52_Pyoeliiyoelii17XNL_Genome.fasta", "py17XNL.fa"),
    ("step_1.fa", "step_1.fa"),
    ("step_2.fa", "step_2.fa"),
]

for ref, name in pairs:
    index = f'index/{name}'
    cmd = f'''\
#
# Additional mappings  and variant calls.
#
mkdir -p index
# Additional evaluation mappping
bwa index -p {index} {ref} 2>> log.txt
hisat2-build {ref} {index} >> log.txt 2>&1
bwa mem -t {CPU} {index} {CONTIG1} {CONTIG2} 2>> log.txt | samtools sort > {name}.illumina.dna.bam 2>> log.txt
minimap2 -t {CPU} -a -x map-ont {ref} {CONTIG_NANO_DNA} 2>> log.txt | samtools sort > {name}.nano.dna.bam 2>> log.txt
hisat2 -p {CPU} --max-intronlen 1500 -x {index} -U {CONTIG_WT} 2>> log.txt | samtools sort > {name}.illumina.rna.bam  2>> log.txt
#
# Variant calling
#
freebayes -f {ref} {name}.illumina.dna.bam -C 5 -p 1 |  bcftools norm --threads 20 -f {ref} -o {name}.fb.vcf
bgzip {name}.fb.vcf
# 
# Filter variants
cat {name}.fb.vcf.gz |bcftools filter -Oz --IndelGap 5 -i  'AF!=0 && GT=="1"' >{name}.fb.filt.vcf.gz
bcftools index {name}.fb.filt.vcf.gz

'''
    print(cmd)

# Genome mapping between reference and final polished.
polished=f'step_{seqid}.fa'
name=[name for ref, name in pairs if ref==GENOME ][0]

mcmd = f'''
# Genome mapping between reference and final polished.
minimap2 -ax asm5 {GENOME} {polished} |samtools sort >{polished}.bam
minimap2 -ax asm5 {polished} {GENOME} |samtools sort >{name}.bam
'''

print(mcmd)

footer = '''
ls -1 *.bam | parallel samtools index {}
'''

print(footer)
```
