## Supplemental File 1 for "Long-Read Genome Assembly and Gene Model Annotations for the Rodent Malaria Parasite *Plasmodium yoelii* 17XNL": sim_map.sh

```
#!/usr/bin/env bash
set -uex

OUTDIR=sim
#rm -rf $OUTDIR
mkdir -p $OUTDIR

cd $OUTDIR

#Link genome to a simple name
ln -sf ../refs/PlasmoDB-52_Pyoeliiyoelii17X_Genome.fasta  py17X.fa

#ASM=../multi_polish_all_scaff_fb/step_4.fa
ASM=../asm/Py17XNL_assembly.fa
GENOME=py17X.fa

REF=$ASM


if  [[ $REF == $GENOME ]]; then
SIM=$ASM
else
SIM=$GENOME
fi


FNAME1=$(basename $SIM)
OUT=$FNAME1"_sim.dna.bam"

FNAME2=$(basename $REF)
INDEX=index/$FNAME2

# Create index folder.
mkdir -p index

# Read simulation parameters.
ERR=0
N=1000000
LEN=150
R1='read1.fq'
R2='read2.fq'


# simulate reads from step_4
pywgsim -S 0 -r 0 -R 0 -N $N -e $ERR -1 $LEN -2 $LEN $SIM $R1 $R2 > log.txt 2>&1

# Create reference genome index.
bwa index -p $INDEX $REF > log.txt 2>&1

# Map simulated reads to reference.
bwa mem $INDEX $R1 $R2 | samtools sort >$OUT
samtools index $OUT

# Call variants with freebayes.
freebayes -f $REF $OUT -C 5 -p 1  >$FNAME1".sim.fb.vcf"
cat $FNAME1".sim.fb.vcf" |bcftools norm --threads 20 -f $REF -o $FNAME1".sim.fb.norm.vcf"
bgzip $FNAME1".sim.fb.norm.vcf"

# Filter VCF
cat $FNAME1".sim.fb.norm.vcf.gz" |bcftools filter -Oz --IndelGap 5 -i  'AF!=0 && GT=="1"' >$FNAME1".sim.fb.norm.filt.vcf.gz"
bcftools index $FNAME1".sim.fb.norm.filt.vcf.gz"
```
