## Supplemental File 1 for "Long-Read Genome Assembly and Gene Model Annotations for the Rodent Malaria Parasite *Plasmodium yoelii* 17XNL": annotate_augustus_gff.py

```
import sys, csv
from collections import defaultdict


def read_ids(fname):
    """
    Reads a tab-delimited idmap file and returns a dictionary of list.
    """
    trans_store = defaultdict(dict)
    stream = csv.DictReader(open(fname), delimiter="\t")
    for row in stream:
        protein_id = "query_pid"
        pid = row.get(protein_id)
        trans_store[pid] = row
    return trans_store


def format_id(chrom, id_, feat):
    new_id = None

    if feat == "gene":
        id_ = id_.replace("g", "")
        # Pad the number with leading zeros
        id_ = id_.zfill(5)
        new_id = "".join([chrom, id_])

    if feat == "transcript" or feat == "mRNA":
        parent, transcript = id_.split(".")
        transcript = transcript.replace("t", "")
        id_ = parent.replace("g", "")
        id_ = id_.zfill(5)
        new_id = "".join([chrom, id_])
        new_id = ".".join([new_id, transcript])

    return new_id


def get_source_gname(chrom, attr, feat, idmap):
    """
    Returns the gname and source of the ID given in attribute.
    If there are multiple names for the same gid, get the one with highest pident.
    """

    id_ = attr.split(";")[0].split("=")[1]
    # curr_id = "_".join([chrom, id_])
    curr_id = id_

    if feat == "gene":
        gname, source, qcovs, pident, matched_protein = get_vals_from_gid(curr_id, idmap)
    else:
        gname, source, qcovs, pident, matched_protein = get_vals_from_tid(curr_id, idmap)
    # var = species + "_"
    var = "query_pid"
    id_ = curr_id.replace(var, "")

    return id_, gname, source, qcovs, pident, matched_protein


def get_vals_from_gid(key, data):
    """
    If there are multiple names for the same gid, get the one with highest pident.
    """
    gname, source, qcovs, pident, matched_protein = None, None, None, None, None

    collect = list()

    for tid, vals in data.items():
        gid = "query_gid"
        if vals.get(gid) == key:
            try:
                gname = vals.get('gene_name')
                source = vals.get('source')
                qcovs = float(vals.get('qcovs'))
                pident = float(vals.get('pident'))
                matched_protein = vals.get('matched_protein')
            except:
                print(vals.get(gid))
                print(key)
                print(vals.get('gene_name'))
                print(vals.get('source'))
                print(vals.get('qcovs'))
                print(vals.get('pident'))
                print("Error")
                sys.exit()
            collect.append((source, qcovs, pident, gname, matched_protein))

    # sort and get the hit largest qcovs and pident
    collect = sorted(collect, key=lambda tup: (tup[2]), reverse=True)

    if collect:
        source, qcovs, pident, gname, matched_protein = collect[0]

    return gname, source, qcovs, pident, matched_protein


def get_vals_from_tid(key, data):
    gname, source, qcovs, pident, matched_protein = None, None, None, None, None

    if key in data:
        gname = data[key].get('gene_name')
        source = data[key].get('source')
        qcovs = data[key].get('qcovs')
        pident = data[key].get('pident')
        matched_protein = data[key].get('matched_protein')
    return gname, source, qcovs, pident, matched_protein


def modify_gene_attr(gid, gname, source, qcovs, pident, matched_protein, org_gid=""):
    if gname is None or gname == "":
        # new_attr = f'ID={gid};gene_id={gid};gene_name={gid};Name={gid};best_match=augustus; original_gene_id={org_gid}'
        new_attr = f'ID={gid};gene_id={gid};gene_name={gid};Name={gid};best_match=augustus'
    else:
        new_attr = f'ID={gid};gene_id={gid};gene_name={gid};Name={gname};best_match={matched_protein};query_coverage={qcovs};percentage_identity={pident}'

    return new_attr


def modify_transcript_attr(tid, gname, source, qcovs, pident, evidence, matched_protein, org_tid=""):
    parent, transcript = tid.split(".")

    # if gname is None or gname == "":
    if evidence == "augustus":
        new_attr = f'ID={tid};Parent={parent};transcript_id={tid};gene_name={parent};Name={tid};best_match={evidence}'
    elif evidence == "locus":
        trans_id = ".".join([gname, transcript])
        new_attr = f'ID={tid};Parent={parent};transcript_id={tid};gene_name={gname};Name={trans_id};best_match={evidence}'

    else:
        trans_id = ".".join([gname, transcript])
        new_attr = f'ID={tid};Parent={parent};transcript_id={tid};gene_name={parent};Name={trans_id};best_match={matched_protein};query_coverage={qcovs};percentage_identity={pident}'

    return new_attr


def modify_attr(attr, tid, gname, gid, org_tid=""):
    parent = tid

    transcript = tid.split(".")[1]

    if gname is None or gname == "":
        # new_attr = f'Parent={parent};transcript_id={parent}; original_transcript_id={org_tid}'
        new_attr = f'Parent={parent};transcript_id={parent}'
    else:
        # trans_id = ".".join([gname, transcript])
        trans_id = ".".join([gid, transcript])
        new_attr = f'Parent={parent};transcript_id={trans_id};gene_name={gname}'

    if attr.startswith("ID"):
        feat = attr.split(";")[0].split(".")[-1]
        id_ = "-".join([parent, feat])
        id1 = "ID=" + id_
        name = "Name=" + id_
        new_attr = ";".join([id1, name, new_attr])

    return new_attr


def modify_gff(gff, idmap):
    tid, gid, gname, source, curr_gname = "", "", "", "", ""

    # Dictionaries to store original ids and corresponding new ids.
    tid_store, gid_store = dict(), dict()

    # print header
    print("##gff-version 3")

    stream = csv.reader(open(gff), delimiter="\t")
    # stream = islice(stream, LIMIT)

    for row in stream:
        if row[0].startswith("#"):
            continue

        chrom, feat, attr = row[0], row[2], row[8]

        if feat == "gene":
            curr_gname = ""
            gid, gname, source, qcovs, pident, matched_protein = get_source_gname(chrom, attr, feat, idmap)
            # if gid=="g118" :
            #     print(gid, gname, source, qcovs, pident)
            #     gene_attr = modify_gene_attr(gid, gname, source, qcovs, pident)
            #     row[8] = gene_attr
            #     print("\t".join(row))
            #     curr_gname = gname
            #     1/0

            org_gid = gid
            gid = format_id(chrom, gid, feat)
            gid_store[gid] = org_gid
            # gene_attr = modify_gene_attr(gid, gname, source, qcovs, pident, matched_protein, org_gid)
            gene_attr = modify_gene_attr(gid, gname, source, qcovs, pident, matched_protein)
            row[8] = gene_attr
            print("\t".join(row))
            curr_gname = gname
            continue

        if feat == "transcript" or feat == "mRNA":
            # We assign a transcript to a gene based on the following evidence
            # 1. species (eg:homo sapiens) - have rblast hits; If so, we add best match organism along with gname.
            # 2. locus  - transcript don't have a hit, but other transcripts of the same family have a hit.
            # 3. augustus - we obtained the transcript from augustus prediction but no reciprocal match to this or related transcripts.

            tid, gname, source, qcovs, pident, matched_protein = get_source_gname(chrom, attr, feat, idmap)

            evidence = 'species'
            if gname is None or gname == "" or gname != curr_gname:
                # This means no evidence for this transcript.
                #  However there may be other transcripts that are annotated from the same gene
                #
                gid = tid.split(".")[0]
                # gid = "_".join(["giraffe", gid])
                gname, source, qcovs, pident, matched_protein = get_vals_from_gid(gid, idmap)
                evidence = 'augustus' if (gname is None or gname == "") else 'locus'

            org_tid = tid
            tid = format_id(chrom, tid, feat)
            tid_store[tid] = org_tid

            tran_attr = modify_transcript_attr(tid, gname, source, qcovs, pident, evidence, matched_protein)
            # tran_attr = modify_transcript_attr(tid, gname, source, qcovs, pident, evidence, matched_protein, org_tid)
            row[8] = tran_attr
            print("\t".join(row))
            continue

        if feat != "gene" or feat != "transcript":
            # new_attr = modify_attr(attr, tid, gname, gid, org_tid)
            new_attr = modify_attr(attr, tid, gname, gid)
            row[8] = new_attr
            print("\t".join(row))
            continue

    return gid_store, tid_store


if __name__ == "__main__":
    map_file = sys.argv[1]  # "recp_protein_map.txt"
    gff_file = sys.argv[2]  # "genes.gff3"
    trans_map = read_ids(map_file)
    org_gid_store, org_tid_store = modify_gff(gff_file, trans_map)

    # Write original-id map file.
    outfile = open("transcript_map.txt", "w")
    header = "\t".join(["org_id", "new_id"])
    outfile.write(header)
    outfile.write("\n")
    for k, v in org_tid_store.items():
        outfile.write("\t".join([v, k]))
        outfile.write("\n")
```
