## Supplemental File 1 for "Long-Read Genome Assembly and Gene Model Annotations for the Rodent Malaria Parasite *Plasmodium yoelii* 17XNL": extract_best_gname.py

```
import csv, sys
from collections import defaultdict


def parse_data(fname):
    data = defaultdict(list)
    stream = csv.DictReader(open(fname), delimiter=",")
    col_names = stream.fieldnames
    query_id, targets = get_targets(col_names)

    def get_vals(row, species):
        qcovs = species + "_qcovs"
        pident = species + "_pident"
        gname = species + "_gene_name"
        pid = species + "_pid"
        qval = 0 if row.get(qcovs) == "" else row.get(qcovs)
        pident = 0 if row.get(pident) == "" else row.get(pident)
        gene = "" if row.get(gname) == "" else row.get(gname)
        sp_pid = "" if row.get(pid) == "" else row.get(pid)
        vals = (species, float(qval), float(pident), gene, sp_pid)
        return vals

    for row in stream:
        pid = row.get(query_id)

        for species in targets:
            data[pid].append(get_vals(row, species))

    return data


def get_targets(col_names):
    """
    Get the targets from column names.
    """
    qid, c = None, set()

    for idx, item in enumerate(col_names):
        if idx == 0:
            qid = item
            continue
        name = item.split("_")[0]
        c.add(name)
    return qid, c


def select_best_gname(store):
    genes = defaultdict(list)

    for tid, vals in store.items():
        gid = ".".join(tid.split(".")[:-1])

        # sort and get the hit largest qcovs and pident
        vals = sorted(vals, key=lambda tup: (tup[1], tup[2]), reverse=True)

        # Filter values where gname is empty.
        filtered = [x for x in vals if x[3] != ""]

        # continue if no gnames.
        if not filtered:
            continue

        # select  the best hit based on qcos and pident
        selected = filtered[0]
        source, qcovs, pident, gene_name, matched_pid  = selected

        genes[gid].append([tid, gid,gene_name, source, qcovs, pident, matched_pid])

    return genes


def extract_duplicates(selected):
    """
    Separates the slected into uniq and duplicated
    """
    uniq, dups = list(), list()

    for gene, vals in selected.items():

        gids = set([x[1] for x in vals])

        if len(gids) == 1:
            uniq.append(gene)
        else:
            dups.append(gene)

    return uniq, dups


def correct_duplicates(selected, data):
    """
    Check if any of the genes are assigned more than ones.
    Correct if so and returns a corrected idmap.
    """
    uniq, duplicates = extract_duplicates(selected)

    named = dict()
    corrected = defaultdict(list)

    # correct duplicates
    for dup in duplicates:

        idx = 1

        hits = selected[dup]

        # assign gname to the hit with best vals.
        hits = sorted(hits, key=lambda x: (x[4], x[5]), reverse=True)
        best = hits[0]
        gid, gname = best[1:3]

        if gid not in named and gname in named.values():
            # Best gene is already assigned. Treat this as duplicate and not as best.
            dup_hits = hits
        else:
            # New best gene.
            named[gid] = gname
            corrected[dup].append(best)
            dup_hits = hits[1:]

        def check_uniq_and_assign(elms, uniq):
            species, qcovs, pident, gene = "", "", "", ""
            for h in elms:
                species, qcovs, pident, gene = h
                if gene not in uniq:
                    break
            return species, qcovs, pident, gene

        # assign unique gnames to the rest of the duplicates.

        for hit in dup_hits:

            trans_id = hit[0]
            gene_id = ".".join(trans_id.split(".")[:-1])

            res = data[trans_id]
            res = sorted(res, key=lambda tup: (tup[1], tup[2]), reverse=True)

            # Filter values where gname is empty.
            filt = [x for x in res if x[3] != ""]

            new_hit = [x for x in filt if x[3] not in named.values()]

            if new_hit and gene_id not in named:

                # if it is a new hit, make sure the gene that is getting assigned is not in uniq.
                # species, qcovs, pident, gene = new_hit[0]
                species, qcovs, pident, gene, = check_uniq_and_assign(new_hit, uniq)

                if gene in uniq and gene_id != selected[gene][0][1]:
                    # if gene id is the same, then it is an isoform,
                    #  so no need to increment idx.
                    gene = gene + "." + str(idx)
                    idx += 1

            else:
                species, qcovs, pident, gene = filt[0]

                if gene_id not in named:
                    gene = gene + "." + str(idx)
                    idx += 1
                else:
                    gene = named[gene_id]

            named[gene_id] = gene
            corrected[gene].append([trans_id, gene_id, gene, species, qcovs, pident])

    # Update corrected to include unique gnames.
    for g in uniq:
        corrected[g].extend(selected[g])

    return corrected


if __name__ == "__main__":
    fname = sys.argv[1]  # "all_union_reciprocal_table.csv"
    header = "\t".join(['query_pid', 'query_gid', 'gene_name', 'source', 'qcovs', 'pident' , 'matched_protein'])
    print(header)
    data = parse_data(fname)

    gene_hits = select_best_gname(data)

    corrected = correct_duplicates(gene_hits, data)

    for gene, results in corrected.items():
        for res in results:
            res = list(map(str, res))
            print("\t".join(res))
```
