## Supplemental File 1 for "Long-Read Genome Assembly and Gene Model Annotations for the Rodent Malaria Parasite *Plasmodium yoelii* 17XNL": get_best_hit.py

```
import argparse, sys, os, csv, re
from itertools import groupby
from operator import itemgetter


def get_basename(file):
    return os.path.splitext(os.path.basename(file))[0]


def read_blast(fname):
    for row in csv.DictReader(open(fname), delimiter="\t"):
        row['qcovs'] = float(row['qcovs'])
        row['pident'] = float(row['pident'])
        row['bitscore'] = float(row['bitscore'])
        row['qlen'] = float(row['qlen'])
        row['slen'] = float(row['slen'])
        yield row


def get_reciprocal_hits(parsed1, parsed2, by_best=False):
    reciprocals = []
    if by_best:
        reciprocals = get_first_best_reciprocal_hit(parsed1, parsed2)
    else:
        reciprocals = get_best_reciprocal_hit(parsed1, parsed2)
    return reciprocals


def get_first_best_reciprocal_hit(parsed1, parsed2):
    """
    Both inputs are list of dictionaries.
    Returns a list of dictionaries of reciprocal hits.
    """

    # Get (qseqid, sseqid) pair as set from parsed1.
    # Get (sseqid,qseqid) pair as set from parsed2.
    # Reciprocals are the intersection of two sets.

    s1, s2 = set(), set()

    for key, val in parsed1.items():
        subject = val['sseqid']
        s1.add((key, subject))

    for key, val in parsed2.items():
        subject = val['sseqid']
        s2.add((subject, key))

    intersected = s1.intersection(s2)
    intersected_id1 = [x[0] for x in intersected]
    reciprocals = [y for x, y in parsed1.items() if x in intersected_id1]

    return reciprocals


def get_best_reciprocal_hit(parsed1, parsed2):
    """
    Both inputs are list of dictionaries.
    Returns a list of dictionaries of reciprocal hits.
    """

    # Get (qseqid, sseqid) pair as set from parsed1.
    # Get (sseqid,qseqid) pair as set from parsed2.
    # Reciprocals are the intersection of two sets.

    s1, s2 = set(), set()

    for key, vals in parsed1.items():
        for val in vals:
            subject = val['sseqid']
            s1.add((key, subject))

    for key, vals in parsed2.items():
        for val in vals:
            subject = val['sseqid']
            s2.add((subject, key))

    intersected = s1.intersection(s2)
    reciprocals = list()

    for key, vals in parsed1.items():
        for val in vals:
            query = val['qseqid']
            sub = val['sseqid']
            if (query, sub) in intersected:
                reciprocals.append(val)
                break

    return reciprocals


def parse_blast(fname, qcovs=0.0, pident=0.0, by_best=False):
    """
    This script filters blast results by qcovs and pident and then extracts the best scoring hit.
    It returns the best scoring hit that satisfied the filtering criteria as a list of dictionaries.
    """

    # Read data.
    stream = read_blast(fname)

    def cond(row):
        return row['qcovs'] >= qcovs and row['pident'] >= pident

    # Filter by condition.
    data = list(filter(cond, stream))

    # Get the hit with highest bit score from the filtered list.
    data.sort(key=itemgetter('qseqid', 'bitscore'), reverse=True)

    for key, items in groupby(data, key=itemgetter('qseqid')):
        if by_best:
            best = next(items)
            key = best['qseqid']
            yield key, best
        else:
            yield key, list(items)


def write_csv(data, outfile):
    """
    Data is a list of dictionaries.
    """
    header = data[0].keys()
    fh = open(outfile, 'w', newline="")
    writer = csv.DictWriter(fh, fieldnames=header, delimiter=",")
    writer.writeheader()
    for row in data:
        writer.writerow(row)
    fh.close()
    return


def parse_files(f1, **kwargs):
    parsed_f1 = dict(parse_blast(fname=f1, qcovs=kwargs['qcov'], pident=kwargs['pident'], by_best=by_best))

    return parsed_f1


def parse_rblast(file1, qcov, pident, by_best):
    parsed1 = parse_files(file1, qcov=qcov, pident=pident, by_best=by_best)

    header = ['qseqid', 'sseqid', 'pident', 'qcovs', 'bitscore','qlen', 'slen']
    print("\t".join(header))

    if by_best:
        for k, v in parsed1.items():
            qid = v['qseqid']
            sid = v['sseqid']
            pident = v['pident']
            qcovs = v['qcovhsp']
            bitscore = v['bitscore']
            qlen=v['qlen']
            slen=v['slen']
            out = [qid, sid, str(pident), str(qcovs), str(bitscore), str(qlen),str(slen)]
            print("\t".join(out))

    else:
        for k, items in parsed1.items():
            for v in items:
                qid = v['qseqid']
                sid = v['sseqid']
                pident = v['pident']
                qcovs = v['qcovhsp']
                qstart = v['qstart']
                qend = v['qend']
                sstart = v['sstart']
                send = v['send']
                qlen = v['qlen']
                slen = v['slen']
                aln_len = v['length']
                bitscore = v['bitscore']


                out = [qid, sid, str(pident), str(qcovs), str(bitscore), str(qlen), str(slen)]
                print("\t".join(out))


if __name__ == "__main__":
    parser = argparse.ArgumentParser(description=''' This script filters and annotates the results of reciprocal blast to produce a table with best matching pairs. \
    It takes the blast results in tabular format and a tabular idmap for annotation. \
    See argument list to know the details of idmap file.
    Output is a  table with source id, target_id, target_gene_name, target_gene_id, target_transcript_id,source_pident, source_qcovhs, source_bitscore. \
    If idmap file is not provided target_gene_name, target_gene_id and target_transcript_id columns will be empty.  
    ''')

    parser.add_argument('--file1', dest='file1', type=str, required=True,
                        help='Tabular blast output file1')
    parser.add_argument('--qcov', dest='qcov', action='store', default=0,
                        help='Minimum query coverage criteria for filtering blast results.')
    parser.add_argument('--pident', dest='pident', action='store', default=0,
                        help='Minimum percentage identity criteria for filtering blast results')
    parser.add_argument('--by_best', dest='by_best', action='store_true', default=False,
                        help="""If specified, extracts the best scoring hit that has a reciprocal match.
                        When not specified, program extracts any of the high scoring hit that has a reciprocal match
                        ( and satisfying other criteria when specified). By default it is False.""")

    args = parser.parse_args()
    file1 = args.file1
    qcov = float(args.qcov)
    pident = float(args.pident)
    by_best = args.by_best
    parse_rblast(file1, qcov, pident, by_best)
```
