## Supplemental File 1 for "Long-Read Genome Assembly and Gene Model Annotations for the Rodent Malaria Parasite *Plasmodium yoelii* 17XNL": make_recpiprocal_table.sh

```
set -ue
#usage: bash make_recpiprocal_table.sh <species_list> 
# first species in the species list should be the source.
#usage: bash make_recpiprocal_table.sh giraffe cow goat sheep human okapi

SCRIPT_DIR=$(dirname "$0")

#
# Extract qseqid, pid, pident, qcovs, bitscore, gene_id, gene_name, transcript_id
# from reciprocal hits and modify header. These are the only columns we need in the final output.
#
source=$1; shift
species_list=$@

rm -f list.txt

for species in $species_list
do
echo "modifying header for $species"
python $SCRIPT_DIR/modify_species_header.py --recp_table ${source}_${species}_reciprocal_hits.csv --outfile g${species}.csv  --qspecies $source --hspecies $species

ls g${species}.csv >>list.txt
done

merge_col=${source}"_id"

# create union table
python $SCRIPT_DIR/merge_reciprocal_hits.py --file_list list.txt --outfile all_union.csv --merge_how union --merge_col $merge_col

# create intersection table
python $SCRIPT_DIR/merge_reciprocal_hits.py --file_list list.txt --outfile all_intersection.csv --merge_how intersection --merge_col $merge_col

#rm -f list.txt
```
