## Supplemental File 1 for "Long-Read Genome Assembly and Gene Model Annotations for the Rodent Malaria Parasite *Plasmodium yoelii* 17XNL": merge_reciprocal_hits.py

```
import pandas as pd
from functools import reduce
import glob, sys, argparse


def main():
    parser = argparse.ArgumentParser(description='''Merges tables''')

    parser.add_argument('--file_list', dest='file_list', type=str, required=True,
                        help='File listing all the tables to merge.')
    parser.add_argument('--outfile', dest='outfile', type=str, required=True,
                        help='Output file name')
    parser.add_argument('--merge_how', dest='merge_how', type=str, required=True,
                        help='''Specify how the tables should be merged. It shpuld be either union, intersection or merge.\
                                if merge is specified a left join on the first table is performed.''' )
    parser.add_argument('--merge_col', dest='merge_col', type=str, required=True,
                        help=''' Specify the ID column to be used for merged.''')   
 
    args = parser.parse_args()
    flist = args.file_list
    outfile = args.outfile
    merge_how = args.merge_how # merge, union or intersection
    merge_column = args.merge_col # 'giraffe_id'

    merge_type = ""

    if merge_how == "merge":
        merge_type = 'left'  # merge on the 1st table.
    elif merge_how == "intersection":
        merge_type = "inner"
    else:
        merge_type = "outer"
    
    frames = list()
     
    fh = open(flist)
    for f in fh:
       df = pd.read_csv(f.strip())
       frames.append(df)

    #frames = [df1, df2, df3, df4, df5]

    result = reduce(lambda left, right: pd.merge(left, right, on=merge_column,
                                                 how=merge_type), frames)

    result.to_csv(outfile, index=False)

    return


if __name__ == "__main__":
    main()
 
```
