## Supplemental File 1 for "Long-Read Genome Assembly and Gene Model Annotations for the Rodent Malaria Parasite *Plasmodium yoelii* 17XNL": modify_species_header.py

```
# This script extracts specific columns from the reciprocal hits table
# and modified the header to include source-species and hit-species names.
# source-species : species you are interested in annotating
# hit-species : target species  

import csv, sys,argparse


def make_header(colnames, hspecies, qspecies):
    new_cols = ["_".join([hspecies, c]) for c in colnames]

    new_cols[0] = qspecies + "_id"  #'giraffe_id'
    new_cols[1] = new_cols[1].replace('sseqid', 'pid')
    return new_cols


if __name__ == "__main__":

    parser = argparse.ArgumentParser(description='''Extracts columns qseqid,sseqid,pident,qcovs,bitscore,gene_name,gene_id,transcript_id from reciprocal hits table and adds hit-species name to the column header''')

    parser.add_argument('--recp_table', dest='recp_fname', type=str, required=True,
                        help='Tabular blast reciprocal output file')
    parser.add_argument('--outfile', dest='outfile', type=str, required=True,
                        help='Output file name')
    parser.add_argument('--qspecies', dest='qspecies', type=str, required=True,
                        help='Query species name')
    parser.add_argument('--hspecies', dest='hspecies', type=str, required=True,
                        help='hit species name')

    args = parser.parse_args()
    fname = args.recp_fname
    outfile = args.outfile
    qspecies = args.qspecies
    hspecies = args.hspecies

    outstream = csv.writer(open(outfile, "w"))
    instream = csv.DictReader(open(fname))

    cols = ['qseqid', 'sseqid', 'pident', 'qcovs',
            'bitscore', 'gene_name', 'gene_id', 'transcript_id']

    new_header = make_header(cols, hspecies, qspecies)

    outstream.writerow(new_header)
   
    for row in instream:
        out = [row['qseqid'], row['sseqid'], row['pident'], row['qcovs'],
               row['bitscore'], row['gene_name'], row['gene_id'], row['transcript_id']]
        outstream.writerow(out)
```
