## Supplemental File 1 for "Long-Read Genome Assembly and Gene Model Annotations for the Rodent Malaria Parasite *Plasmodium yoelii* 17XNL": rblast.sh

```
#!/usr/bin/env bash
set -ue

# set variables.
FASTA1=""
FASTA2=""
OUTDIR="."
FWD_BLAST=""
REV_BLAST=""
EVAL=10
NPROC=15

usage="$(basename "$0") [-1 -2 -f -r -e -o -h] \n
This program runs reciprocal blast between file1 and file two and produce results
in tabular format. \n

    -h    show this help text \n
    -1    Protein or Nucleotide fasta file1 \n
    -2    Protein or Nucleotide fasta file2 \n
    -f    Forward blast from file1 to file2.\n
    -r    Reverse blast from file2 to file1. \n
    -e    Blast evalue threshold for saving hits \n 
    -p    No.of processors \n
    -o    Output directory \n "

# Option strings
SHORT=1:2:f:r:e:o:p:h

# read the options
OPTS=`getopt $SHORT $*`

if [ $? != 0 ] ; then echo "Failed to parse options...exiting." >&2 ; exit 1 ;fi

eval set -- "$OPTS"

for i; do
   case "$i" in
       -1 ) FASTA1="$2"; shift 2 ;;
       -2 ) FASTA2="$2" ; shift 2 ;;
       -o ) OUTDIR="$2" ; shift 2 ;;
       -e ) EVAL="$2" ; shift 2 ;;
       -f ) FWD_BLAST="$2" ; shift 2 ;;
       -r ) REV_BLAST="$2" ; shift 2 ;;
       -p ) NPROC="$2" ; shift 2 ;;
       -h | --help ) echo -e $usage ; exit ;;
       -- ) shift; break ;;
       esac
done

# check if mandatory options are present
if  [ -z "$FASTA1" ] || [ -z "$FASTA2" ] || [ -z "$FWD_BLAST" ] || [ -z "$REV_BLAST" ]; then
   echo "File1, file2, fwd-task and rev-task must be specified.\n"
   echo "Use bash rblast.sh -h  to see the options" 
   exit
fi

# check if blast options are valid.
case $FWD_BLAST in
   blastn|blastp|tblastn|tblastx|blastx) ;;
   *) echo "Invalid blast option in input." ;exit
esac

case $REV_BLAST in
   blastn|blastp|tblastn|tblastx|blastx) ;;
   *) echo "Invalid blast option in input." ;exit
esac

# check if input files exists
if [ ! -f "$FASTA1" ]  || [ ! -f "$FASTA2" ]; then
    echo "One or both input files cannot be found. Exiting"
    exit
fi

# check if outdir exists
if [ ! -d "$OUTDIR" ]; then
    mkdir -p $OUTDIR
fi

# create db directory
db_dir=$OUTDIR/blast_dbs
mkdir -p $db_dir

file1=$(basename $FASTA1)
file2=$(basename $FASTA2)
prefix1=${file1%%.*}
prefix2=${file2%%.*}

out1=${prefix1}"_vs_"${prefix2}".txt"
out2=${prefix2}"_vs_"${prefix1}".txt"

# create blast databases.
case $FWD_BLAST in
    blastp)
      db_type2="prot"
      db_file2=$FASTA2
      ;;
    blastn)
      db_type2="nucl"
      db_file2=$FASTA2
      ;;
    tblastn)
      db_type2="nucl"
      db_file2=$FASTA2
      ;;   
    blastx)
      db_type2="prot"
      db_file2=$FASTA2
      ;;
    tblastx)
      db_type2="nucl"
      db_file2=$FASTA2
      ;;
esac

case $REV_BLAST in
   blastp)
      db_type1="prot"
      db_file1=$FASTA1
      ;;
    blastn)
      db_type1="nucl"
      db_file1=$FASTA1
      ;;
    tblastn)
      db_type1="nucl"
      db_file1=$FASTA1
      ;;
    blastx)
      db_type1="prot"
      db_file1=$FASTA1
      ;;
    tblastx)
      db_type1="nucl"
      db_file1=$FASTA1
      ;;
esac

db1=${db_dir}/$(basename $db_file1)
db2=${db_dir}/$(basename $db_file2)

makeblastdb -in $db_file1 -dbtype $db_type1  -out $db1
makeblastdb -in $db_file2 -dbtype $db_type2  -out $db2

fwd_flag=""
rev_flag=""

# run blast
HEADER="qseqid\tsseqid\tstitle\tstaxids\tpident\tmismatch\tgaps\tppos\tqcovhsp\tqcovs\tqlen\tslen\tlength\tqstart\tqend\tsstart\tsend\tsstrand\tevalue\tbitscore\tscore"
echo -e $HEADER >header.txt


# tblastx do not have -task option; so running it  separately.
# Forward tblastx
if [ $FWD_BLAST = "tblastx" ]; then
   # run fwd-blast with query1 on query2-db
   echo "Running forward  tblastx between $FASTA1 and $FASTA2 (DB=$db2)"
   cat $FASTA1 | parallel -j $NPROC --block 1000k --recstart '>' --pipe $FWD_BLAST -evalue $EVAL -outfmt '"6 qseqid sseqid stitle staxids pident mismatch gaps ppos qcovhsp qcovs qlen slen length qstart qend sstart send sstrand evalue bitscore score"' -db $db2 -query - > ${OUTDIR}/$out1

    cat header.txt ${OUTDIR}/$out1 >tmp.txt
    mv tmp.txt ${OUTDIR}/$out1
    fwd_flag="done"
fi


# Reverse tblastx
if [ $REV_BLAST = "tblastx" ]; then
   echo "Running reverse tblastx between $FASTA2 and $FASTA1 (DB=$db1)"
   cat $FASTA2 | parallel -j $NPROC --block 1000k --recstart '>' --pipe $REV_BLAST -evalue $EVAL -outfmt '"6 qseqid sseqid stitle staxids pident mismatch gaps ppos qcovhsp qcovs qlen slen length qstart qend sstart send sstrand evalue bitscore score"' -db $db1 -query - > ${OUTDIR}/$out2

   cat header.txt ${OUTDIR}/$out2 >tmp.txt
   mv tmp.txt ${OUTDIR}/$out2
   rev_flag="done"
fi


# Run forward blast with file1 on file2
if [ -z "$fwd_flag" ]; then
echo "Running forward $FWD_BLAST between $FASTA1 and $FASTA2 (DB=$db2)"
cat $FASTA1 | parallel -j $NPROC --block 1000k --recstart '>' --pipe $FWD_BLAST -task $FWD_BLAST -evalue $EVAL -outfmt '"6 qseqid sseqid stitle staxids pident mismatch gaps ppos qcovhsp qcovs qlen slen length qstart qend sstart send sstrand evalue bitscore score"' -db $db2 -query - > ${OUTDIR}/$out1

cat header.txt ${OUTDIR}/$out1 >tmp.txt
mv tmp.txt ${OUTDIR}/$out1
fi


# Run reverse blast with file2 on file1
if [ -z "$rev_flag" ]; then
echo "Running reverse $REV_BLAST between $FASTA2 and $FASTA1 (DB=$db1)"
cat $FASTA2 | parallel -j $NPROC --block 1000k --recstart '>' --pipe $REV_BLAST -task $REV_BLAST -evalue $EVAL -outfmt '"6 qseqid sseqid stitle staxids pident mismatch gaps ppos qcovhsp qcovs qlen slen length qstart qend sstart send sstrand evalue bitscore score"' -db $db1 -query - > ${OUTDIR}/$out2

cat header.txt ${OUTDIR}/$out2 >tmp.txt
mv tmp.txt ${OUTDIR}/$out2
fi

rm -f header.txt
```
