## Supplemental File 1 for "Long-Read Genome Assembly and Gene Model Annotations for the Rodent Malaria Parasite *Plasmodium yoelii* 17XNL": rblast_combine.py

```
import argparse, sys, os, csv, re
from itertools import groupby
from operator import itemgetter


def get_basename(file):
    return os.path.splitext(os.path.basename(file))[0]


def read_blast(fname):
    for row in csv.DictReader(open(fname), delimiter="\t"):
        row['qcovs'] = float(row['qcovs'])
        row['pident'] = float(row['pident'])
        row['bitscore'] = float(row['bitscore'])
        yield row


def get_reciprocal_hits(parsed1, parsed2, by_best=False):
    reciprocals = []
    if by_best:
        reciprocals = get_first_best_reciprocal_hit(parsed1, parsed2)
    else:
        reciprocals = get_best_reciprocal_hit(parsed1, parsed2)
    return reciprocals


def get_first_best_reciprocal_hit(parsed1, parsed2):
    """
    Both inputs are list of dictionaries.
    Returns a list of dictionaries of reciprocal hits.
    """

    # Get (qseqid, sseqid) pair as set from parsed1.
    # Get (sseqid,qseqid) pair as set from parsed2.
    # Reciprocals are the intersection of two sets.

    s1, s2 = set(), set()

    for key, val in parsed1.items():
        subject = val['sseqid']
        s1.add((key, subject))

    for key, val in parsed2.items():
        subject = val['sseqid']
        s2.add((subject, key))

    intersected = s1.intersection(s2)
    intersected_id1 = [x[0] for x in intersected]
    reciprocals = [y for x, y in parsed1.items() if x in intersected_id1]

    return reciprocals


def get_best_reciprocal_hit(parsed1, parsed2):
    """
    Both inputs are list of dictionaries.
    Returns a list of dictionaries of reciprocal hits.
    """

    # Get (qseqid, sseqid) pair as set from parsed1.
    # Get (sseqid,qseqid) pair as set from parsed2.
    # Reciprocals are the intersection of two sets.

    s1, s2 = set(), set()

    for key, vals in parsed1.items():
        for val in vals:
            subject = val['sseqid']
            s1.add((key, subject))

    for key, vals in parsed2.items():
        for val in vals:
            subject = val['sseqid']
            s2.add((subject, key))

    intersected = s1.intersection(s2)
    reciprocals = list()

    for key, vals in parsed1.items():
        for val in vals:
            query = val['qseqid']
            sub = val['sseqid']
            if (query, sub) in intersected:
                reciprocals.append(val)
                break

    return reciprocals


def parse_blast(fname, qcovs=0.0, pident=0.0, by_best=False):
    """
    This script filters blast results by qcovs and pident and then extracts the best scoring hit.
    It returns the best scoring hit that satisfied the filtering criteria as a list of dictionaries.
    """

    # Read data.
    stream = read_blast(fname)

    def cond(row):
        return row['qcovs'] >= qcovs and row['pident'] >= pident

    # Filter by condition.
    data = list(filter(cond, stream))

    # Get the hit with highest bit score from the filtered list.
    data.sort(key=itemgetter('qseqid', 'bitscore'), reverse=True)

    for key, items in groupby(data, key=itemgetter('qseqid')):
        if by_best:
            best = next(items)
            key = best['qseqid']
            yield key, best
        else:
            yield key, list(items)


def write_csv(data, outfile):
    """
    Data is a list of dictionaries.
    """
    header = data[0].keys()
    fh = open(outfile, 'w', newline="")
    writer = csv.DictWriter(fh, fieldnames=header, delimiter=",")
    writer.writeheader()
    for row in data:
        writer.writerow(row)
    fh.close()
    return


def read_idmap(fname):
    """
    This function reads a tab delimiterd idmap file with header
    gene_name, gene_id, transcript_id, protein_id and returns
    two dictionaries, one keyed with protein_id and the other with trasncript_id
    """
    tid_store, pid_store = dict(), dict()

    stream = csv.DictReader(open(fname), delimiter="\t")
    for row in stream:
        pid = row.get('protein_id')
        tid = row.get('transcript_id')

        # store without the version
        # pid = re.sub(r'\.\d+$', '', pid)
        # tid = re.sub(r'\.\d+$', '', tid)

        if pid and pid != "-":
            pid_store.setdefault(pid, {}).update(row)

        if tid and tid != "-":
            tid_store.setdefault(tid, {}).update(row)

    return tid_store, pid_store


def get_annotations(data, idmap):
    """
    This function gets gene_id, gene_name and transcript_id
    corresponding to protein_id in input data.
    """
    if not os.path.isfile(idmap):
        print(f"{idmap} file not found. Gene_name and other id columns will be empty.")
        return data

    tid_store, pid_store = read_idmap(idmap)

    for item in data:
        item['gene_name'], item['gene_id'] = "", ""
        item['transcript_id'], item['protein_id'] = "", ""

        id_ = item['sseqid']

        # remove accession version
        id_ = re.sub(r'\.\d+$', '', id_)

        if id_ in pid_store:
            values = pid_store.get(id_)

        else:
            values = tid_store.get(id_)

        if values:
            item['gene_name'] = values.get('gene_name')
            item['gene_id'] = values.get('gene_id')
            item['protein_id'] = values.get('protein_id')
            item['transcript_id'] = values.get('transcript_id')

    return data


def parse_files(f1, f2, **kwargs):
    parsed_f1 = dict(parse_blast(fname=f1, qcovs=kwargs['qcov'], pident=kwargs['pident'], by_best=by_best))
    parsed_f2 = dict(parse_blast(fname=f2, qcovs=kwargs['qcov'], pident=kwargs['pident'], by_best=by_best))

    return parsed_f1, parsed_f2


def filter_annotate_rblast(file1, file2, out, qcov, pident, by_best=False, idmap=''):
    # Filter blast results based on thresholds and get highest scored hit.
    parsed1, parsed2 = parse_files(file1, file2, qcov=qcov, pident=pident, by_best=by_best)
    # Get reciprocal hits in list of dictionaries.
    reciprocals = get_reciprocal_hits(parsed1, parsed2, by_best=by_best)
    outdir = os.path.dirname(os.path.realpath(out))

    # check if output directory exists
    if not os.path.isdir(outdir):
        try:
            os.mkdir(outdir)
        except OSError:
            print(f'Creation of the directory {outdir} failed')
            sys.exit()

    def add_columns(reciprocals):
        for item in reciprocals:
            item['protein_id'], item['gene_id'], item['gene_name'], item['transcript_id'] = "", "", "", ""
        return reciprocals

    if reciprocals:
        reciprocals = get_annotations(reciprocals, idmap) if idmap else reciprocals
        reciprocals = add_columns(reciprocals) if not idmap else reciprocals
        write_csv(reciprocals, out)
    else:
        print(f"No reciprocal hits are found.")
    return


if __name__ == "__main__":
    parser = argparse.ArgumentParser(description=''' This script filters and annotates the results of reciprocal blast to produce a table with best matching pairs. \
    It takes the blast results in tabular format and a tabular idmap for annotation. \
    See argument list to know the details of idmap file.
    Output is a  table with source id, target_id, target_gene_name, target_gene_id, target_transcript_id,source_pident, source_qcovhs, source_bitscore. \
    If idmap file is not provided target_gene_name, target_gene_id and target_transcript_id columns will be empty.  
    ''')

    parser.add_argument('--file1', dest='file1', type=str, required=True,
                        help='Tabular blast output file1')
    parser.add_argument('--file2', dest='file2', type=str, required=True,
                        help='Tabular blast output file2')
    parser.add_argument('--qcov', dest='qcov', action='store', default=0,
                        help='Minimum query coverage criteria for filtering blast results.')
    parser.add_argument('--pident', dest='pident', action='store', default=0,
                        help='Minimum percentage identity criteria for filtering blast results')
    parser.add_argument('--by_best', dest='by_best', action='store_true', default=False,
                        help="""If specified, extracts the best scoring hit that has a reciprocal match.
                        When not specified, program extracts any of the high scoring hit that has a reciprocal match
                        ( and satisfying other criteria when specified). By default it is False.""")
    parser.add_argument('--out', dest='out', action='store', default='./reciprocals.csv',
                        help='Specify the output file. default is ./reciprocals.csv')
    parser.add_argument('--idmap', dest='idmap', type=str,
                        help='''Tab delimited file with four columns that contains the mapping of different ids. 
                            The ids in this file corresponds to the hit species in file1. 
                            First line should contain the header with column names as gene_id,transcript_id, protein_id, 
                            gene_name. If this file is not given, the corresponding columns in the output file will be 
                            empty.
                            .''')
    args = parser.parse_args()
    file1 = args.file1
    file2 = args.file2
    idmap = args.idmap
    qcov = float(args.qcov)
    pident = float(args.pident)
    by_best = args.by_best
    out = args.out
    if not idmap:
        print(f"{idmap} file not given. Gene_name and other id columns will be empty.")

    filter_annotate_rblast(file1, file2, out, qcov, pident, by_best, idmap)
```
