## Supplemental File 1 for "Long-Read Genome Assembly and Gene Model Annotations for the Rodent Malaria Parasite *Plasmodium yoelii* 17XNL": run_rblast.sh

```
set -ue
# usage:bash run_rblast.sh blastp blastp QUERY TARGET(s)
# usage: bash run_rblast.sh blastp blastp giraffe_proteins.fa cow_proteins.fa sheep_proteins.fa

SCRIPT_DIR=$(dirname "$0")

# set variables.

acc_map=""
fwd_blast="blastp"
rev_blast="blastp"
qcov=0 #70
pident=0 #70
evalue=10
nproc=15
source=""
#target=${arr[@]}
target=()
outdir="rblast_res"

usage="$(basename "$0") [-a -f -r -q -i -e -p -s -t -o -h] \n
This program runs reciprocal blast between file1 and file two and produce results
in tabular format. \n

    -h    Show this help text \n
    -a    Accession map file in tab delimited format with "gene_id,transcript_id,protein_id,gene_name" columns   \n
    -f    Forward blast from file1 to file2.\n
    -r    Reverse blast from file2 to file1. \n
    -q    Query coverage threshold used for parsing blast output [70]  \n
    -i    Percentage identity threshold used for parsing blast output [70] \n 
    -e    Evalue threshold [10] \n
    -p    No.of processors \n
    -o    Output directory \n 
    -s    Source/Query fasta file \n
    -t    Target or target(s) of fasta files \n "    


while getopts ":a:f:r:q:i:e:p:o:s:t:h" opt; do
    case $opt in
        a) acc_map="${OPTARG}";;
        f) fwd_blast="${OPTARG}";;
        r) rev_blast="${OPTARG}";;
	q) qcov="${OPTARG}";;
	i) pident="${OPTARG}";;
	e) evalue="${OPTARG}";;
	p) nproc="${OPTARG}";;
	o) outdir="${OPTARG}";;
	s) source="${OPTARG}";;
	t) target="${OPTARG}";;
	h) echo -e $usage ; exit ;;
        \?) exit "Invalid option -$OPTARG";;
    esac
done


source_prefix=${source%.*}
source_prefix=${source_prefix##*/}

target_species=()

mkdir -p $outdir

for species in $target
do


species_prefix=${species%.*}
species_prefix=${species_prefix##*/}
target_species+=(${species_prefix})

echo "running rblast between $source and $species"
time bash $SCRIPT_DIR/rblast.sh -1 ${source} -2 ${species} -f $fwd_blast -r $rev_blast -o $outdir -e $evalue -p $nproc

echo "creating reciprocal tables between $source_prefix and $species_prefix"

# highest scoring best hit
#time python $SCRIPT_DIR/rblast_combine.py --file1 ${outdir}/${source_prefix}_vs_${species_prefix}.txt --file2 ${outdir}/${species_prefix}_vs_${source_prefix}.txt --qcov 0 --pident 0 --idmap $acc_map  --out ${outdir}/${source_prefix}_${species_prefix}_reciprocal_hits.csv --by_best

if [ -z "$acc_map" ]; then
# any of the best with qcov>=$qcov pident>=$pident
time python $SCRIPT_DIR/rblast_combine.py --file1 ${outdir}/${source_prefix}_vs_${species_prefix}.txt --file2 ${outdir}/${species_prefix}_vs_${source_prefix}.txt --qcov $qcov --pident $pident --out ${outdir}/${source_prefix}_${species_prefix}_reciprocal_hits.csv

else
# any of the best with qcov>=$qcov pident>=$pident
time python $SCRIPT_DIR/rblast_combine.py --file1 ${outdir}/${source_prefix}_vs_${species_prefix}.txt --file2 ${outdir}/${species_prefix}_vs_${source_prefix}.txt --qcov $qcov --pident $pident --out ${outdir}/${source_prefix}_${species_prefix}_reciprocal_hits.csv --idmap $acc_map

echo -e "qcovs threshold is $qcov and pident threshold is $pident"

fi

echo "--------"
done

echo "rblast is completed. Results are in files ending 'reciprocal_hits.csv' "

#
# Combine the reciprocal hits from multiple target species and get the gene name from the best target species hit
# with highest query coverage and percentage identity.
#

arr_len=${#target_species[@]}

# Remove exiting list file if any
rm -f ${outdir}/list.txt

for ((i = 0; i < $arr_len; ++i))
do
sp=${target_species[$i]}

echo "modifying header for $sp"
python $SCRIPT_DIR/modify_species_header.py --recp_table ${outdir}/${source_prefix}_${sp}_reciprocal_hits.csv --outfile ${outdir}/g_${sp}.csv  --qspecies $source_prefix --hspecies $sp
ls ${outdir}/g_${sp}.csv >>${outdir}/list.txt

done


# create union table
merge_col=${source_prefix}"_id"

python $SCRIPT_DIR/merge_reciprocal_hits.py --file_list ${outdir}/list.txt --outfile ${outdir}/all_union.csv --merge_how union --merge_col $merge_col

if [ ! -z "$acc_map" ]; then
# Extract the best gene name from target species with highest query coverage and percentage identity.
python $SCRIPT_DIR/extract_best_gname.py ${outdir}/all_union.csv >${outdir}/annotated_gene_names.txt

fi
# clean up

rm -f ${outdir}/g_*.csv
```
