## Supplemental File 1 for "Long-Read Genome Assembly and Gene Model Annotations for the Rodent Malaria Parasite *Plasmodium yoelii* 17XNL": README.txt

scripts: folder containing all commands used in the generation of P.yoelii 17XNL agenome assembly and annotation.

Makefile.pacbio : commands  used in Pacbio genome assembly and annotation.

Makefile.ont : commands used in the nanopore genome assembly and annotation.

1) run/ : contains shell scripts used in the analysis. These scripts uses accessory code that are in other subfolders under scripts/

2) ann_fix/ : contains commands used to make modifications to the braker annotation output files.

3) asm_mod/ : scripts used to make modifications to canu output assembly.

4) ncRNA/ : scripts used to find and add non-coding RNAs to annotation

5) rblast/ : scripts used in the reciprocal blast based annotation.

6) ont_asm : scripts used in the Nanopore based genome assembly

7) cons : scripts used on consensus genome generation.

8) misc : miscelanious scripts used in various steps of the analysis.


Description of scripts in each folder is given below.

run/:

This folder contains the wrapper scripts that needs to be run. It uses the accessory scripts in other subfolders under scripts/

1) braker.sh : commands to run braker2
2) rblast_ann.sh : commands to annotate braker2 output using reciprocal blast
3) map.sh : commands used in mapping reads to assembly
4) prep_cons.sh : commands used in modifying the header of consensus fasta
5) get_ncRNA.sh : commands used in finding ncRNAs and adding them to annotation.
6) hybrid_v2.sh : commands used in generating version2 of hybrid annotation which contains the genes that are present in nanopore-only model but were absent in hybrid-model version1
7) get_hybrid_v2_fasta.sh : creates transcript/protein/cds fasta files for hybrid_v2 annotation
8) fix_ann.sh : commands used in fixing annotations in apicoplast/mitochondria using prokka and rblast 
9) diffs.sh :  commands used in generating genomic differences and annotating them
10) cds_diffs.sh : commands used in generating cds variants
11) utr_len.sh : commands for generating UTR length table.
12) submission.sh : commands useful for preparing GFF3 file for genbank submission

ncRNA/ :
This folder contains scripts used in generating ncRNA annotations

1) bed.py : convert tRNASCAN-SE output to a bed file
2) get_tRNAs.sh : makes tRNA predictions
3) get_rRNAs.sh : identifies rRNAs in the genome.
4) ncRNA_gff.py : creates ncRNA gff file from bed file.
5) best.py : extracts the best blast hit.
5) get_last_gid.py : extracts the geneid of the last gene in each chromosome

ann_fix/ :
This folder contains the commands and scripts used in modifying annotation

1) combine_prokka_braker_gff.py : script produces a modified gff3 that combines both braker2 and prokka-rblast annotations for apicoplast and mitochondrial chromosomes
2) lift_nano_genes.py : script adds missing genes from nanopore-only model to the hybrid model.
3) API_fix_commands.txt  : commands used in fixing apicoplast annotations by combining braker2 and prokka predictions.
4) MIT_fix_commands.txt  :commands used in fixing mitochondria annotations by combining braker2 and prokka predictions.

rblast/ :
This folder contains the accessory scripts to create reciprocal blast based annotation

1) run_rblast.sh : the wrapper script for running reciprocal blast
2) rblast.sh : runs reciprocal blast
3) rblast_combine.py : merges forward and reverse blast to create reciprocal hits table.
4) merge_reciprocal_hits.py : creates a union table of reciprocal blast hits from multiple species.
5) extract_best_gname.py : creates a table with the best reciprocal hit that was used in annotating gene names
6) annotate_augustus_gff.py : used to create an annotated braker2 file using the results formreciprocal blast

ont_asm/ :
This folder contains scripts used in generation Nanopore assembly.

1) flye.sh : Nanopore assembly creation using flye
2) multi_polish.py : Polish the assembly in multiple iterations
3) map_ont.sh  : commands used to create nanopore-assembly based bam files
4) extra_commands.txt : miscellanious commands used in creating the final assembly


asm_mod/ :
This folder contains the scripts used in modifying the canu assembly output.

1) asm_mod_commands.txt : all commands used in making changes to canu assembly output
2) parse_aln.py : parses bam file to create coverage percentage per chromosome
3) assign.sh : assign chromosome names based on coverage percentage.
4) circularize.sh : used in the circularization of apicoplast with circlator
5) rearrange.sh : fix circularization output
6) mt_correct.sh : creates a single copy of MIT
7) modify_header.py : modifies fasta header

cons/ :
This folder contains the scripts used in generating consensus genome assembly

1)consensus.py : script that produces consensus genome

misc/ :
This folder contains the miscellanious scripts

1) vcf2table.py : converts vcf file to table format
2) make_var_table.py : used in generating cds variant table
3) add_matched_prot.py : adds 17XNL protein names to variants
4) format_gff.py : adds locus tag and other Genbank specifications to GFF3 file.
5) attributes_gff.py : parses GFF3 file to extract different attributes
6) add_gids.py : adds gene-ids to the input file.
7) replace_header.py : modifies fasta header.
8) atMask.mat: matrix used in masking repeats.
9) DotPrep.py : prepares a nucmer output delta file for visualization in Dot
10) dotter.sh : wrapper script used to produce dotplot.
