## Supplemental File 1 for "Long-Read Genome Assembly and Gene Model Annotations for the Rodent Malaria Parasite *Plasmodium yoelii* 17XNL": braker.sh

```
# 
# These commands are used to generate the files required for running braker2 annotation and use them to run braker2 annotation
#

# conda activate braker2

ASM_DIR=asm
ASM_IN=${ASM_DIR}/cons2.fa
ASM=${ASM_DIR}/17XNL_PSU.fa
IDX_DIR=${ASM_DIR}/index
IDX=${IDX_DIR}/PY17XNL_PSU

ALIGN_DIR=bam
ILLUM_RNA=data/WT_all.fq.gz
ONT_RNA=data/nano_rna.fq.gz
ILLUM_BAM=${ALIGN_DIR}/illum_rna_17XNL_PSU.bam
ILLUM_FWD=${ALIGN_DIR}/illum_fwd.bam
ILLUM_REV=${ALIGN_DIR}/illum_rev.bam

ONT_BAM=${ALIGN_DIR}/ont_rna_17XNL_PSU.bam
ONT_FWD=${ALIGN_DIR}/ont_fwd.bam
ONT_REV=${ALIGN_DIR}/ont_rev.bam

ANN_DIR=ann
BRAKER_HYBD=${ANN_DIR}/braker2_hybrid
BRAKER_NANO=${ANN_DIR}/braker2_nano

SPECIES_HYBD=PY_hybrid
SPECIES_NANO=PY_nano

#BRAKER_OUT=${BRAKER_HYBD}
#SPECIES=${SPECIES_HYBD}

# Mask the genome
echo
echo make mask_genome ASM_IN=${ASM_IN} ASSEMBLY=${ASM}

# Make hisat2 indices
echo
echo make hisat2_index ASSEMBLY=$ASM INDEX=$IDX INDEX_DIR=$IDX_DIR

# Align illumina reads
echo
echo make hisat2 INDEX=$IDX ILLUM_RNA=$ILLUM_RNA OUTPUT=${ILLUM_BAM}

# Split illumina mapping by strand
echo
echo make split_by_strand INBAM=${ILLUM_BAM} OUT_FWD=${ILLUM_FWD} OUT_REV=${ILLUM_REV}

# Align ont reads
echo
echo make minimap2_rna ASSEMBLY=$ASM NANO_RNA=${ONT_RNA} OUTPUT=${ONT_BAM} SAMPLE=17XNL_PSU_ont

# Split ont mapping by strand
echo
echo make split_by_strand INBAM=${ONT_BAM} OUT_FWD=${ONT_FWD} OUT_REV=${ONT_REV}

# Run braker annotation in hybrid mode 
echo 
echo make braker_hybrid_ann ASSEMBLY=$ASM SPECIES=${SPECIES_HYBD} ILLUM_FWD=${ILLUM_FWD} ILLUM_REV=${ILLUM_REV} ONT_FWD=${ONT_FWD} ONT_REV=${ONT_REV} BRAKER_OUT=${BRAKER_HYBD}

# Run braker annotation in nanopore only mode.
echo
echo make braker_nano_ann ASSEMBLY=$ASM SPECIES=${SPECIES_NANO} ONT_FWD=${ONT_FWD} ONT_REV=${ONT_REV} BRAKER_OUT=${BRAKER_NANO}
```
