## Supplemental File 1 for "Long-Read Genome Assembly and Gene Model Annotations for the Rodent Malaria Parasite *Plasmodium yoelii* 17XNL": cds_diffs.sh

```
#
# The commands produce an intersection of CDS feature and genomic differences.
# Ouput is a text file of CDS variants 
# Also cds variants vcf file is generated.
#

DIFFS=diffs
CDS=${DIFFS}/cds_diffs
SNPEFF=${DIFFS}/snpEff
REFS=refs
ANN=ann
SCRIPTS=scripts/misc

GFF=${REFS}/PlasmoDB-56_Pyoeliiyoelii17X.gff
REF_CDS_GFF=${CDS}/17X_PLD_cds.gff3
REF_CDS_BED=${CDS}/17X_PLD_cds.bed
PROT_MATCH=${ANN}/prot_match.txt

VCF=${DIFFS}/17XNL_PSU_to_17X_PLD.vcf.gz
SNPEFF_DB=PY17X_PLD

#Output
CDS_VCF=${CDS}/17XNL_PSU_cds_ann_short.vcf
CDS_TXT=${CDS}/17XNL_PSU_cds_ann_short.txt
OUT=${CDS}/17XNL_PSU_cds_variants.txt

#Create directories
echo mkdir -p $CDS

# Extract CDS feature from GFF3 file
echo
echo "cat ${GFF} |awk '\$3==\"CDS\" {print}' >${REF_CDS_GFF}"

# Create CDS bed file
echo
echo "cat ${REF_CDS_GFF} | awk 'BEGIN{OFS=\"\t\"} {print \$1,\$4,\$5,\".\",\".\",\$7}' >${REF_CDS_BED}"

# Get annotated CDS variants
echo

echo "java -jar ~/src/snpEff/snpEff.jar ${SNPEFF_DB} ${VCF}  -fi ${REF_CDS_BED} -no-upstream -no-downstream -no-intergenic -no-utr -s ${CDS}/cds_diffs_summary.html  >${CDS_VCF}"

# Create a table from vcf file
echo
echo "python ${SCRIPTS}/vcf2table.py ${CDS_VCF} | grep "protein_coding" >${CDS_TXT}"

# Create index
echo
echo bgzip ${CDS_VCF}
echo bcftools index ${CDS_VCF}.gz

#Intersect cds variants with gff3 so as to get the pids.
echo
echo "bedtools intersect -wb -a ${CDS_VCF}.gz -b ${REF_CDS_GFF}  >${CDS}/intersect.txt"

# Adding 17XNL protein names to the intersection file.
echo
echo "python ${SCRIPTS}/add_matched_prot.py ${PROT_MATCH} ${CDS}/intersect.txt >${CDS}/out.txt"

echo
echo "python ${SCRIPTS}/make_var_table.py ${CDS_TXT} ${CDS}/out.txt >${OUT}"
```
