## Supplemental File 1 for "Long-Read Genome Assembly and Gene Model Annotations for the Rodent Malaria Parasite *Plasmodium yoelii* 17XNL": diffs.sh

```
REFS=refs
ASM=asm
PY17X=${REFS}/PlasmoDB-56_Pyoeliiyoelii17X_Genome.fasta
PY17XNL=${ASM}/17XNL_PSU.fa


DIFFS=diffs
SNPEFF=${DIFFS}/snpEff
VIEW_17X=17XNL_PSU_to_17X_PLD.vcf
VIEW_17X_STATS=17XNL_PSU_to_17X_PLD.vcfstats.txt
VIEW_17X_ANN=17XNL_PSU_to_17X_PLD_ann.vcf
VIEW_17X_ANN_HTML=17XNL_PSU_to_17X_PLD_ann_summary.html
VIEW_17X_ANN_CSV=17XNL_PSU_to_17X_PLD_ann_summary.csv

VIEW_17XNL=17X_PLD_to_17XNL_PSU.vcf
VIEW_17XNL_STATS=17X_PLD_to_17XNL_PSU.vcfstats.txt
VIEW_17XNL_ANN=17X_PLD_to_17XNL_PSU_ann.vcf
VIEW_17XNL_ANN_HTML=17X_PLD_to_17XNL_PSU_ann_summary.html
VIEW_17XNL_ANN_CSV=17X_PLD_to_17XNL_PSU_ann_summary.csv


# Create directory
echo mkdir -p ${SNPEFF}

# Differences in assembly from 17X view
echo
echo make asm_diffs GENOME=${PY17X} ASSEMBLY=${PY17XNL} SAMPLE=17XNL_PSU_asm DIFFS=${DIFFS} DIFF_VCF=${DIFFS}/${VIEW_17X} STATS=${DIFFS}/${VIEW_17X_STATS}

# Annotate vcf file.
# Build snpEff database for 17X view
echo
echo java -jar ~/src/snpEff/snpEff.jar build -gff3 -v PY17X_PLD
# Annotate vcf
echo "java -jar ~/src/snpEff/snpEff.jar PY17X_PLD ${DIFFS}/${VIEW_17X}.gz -csvStats ${SNPEFF}/${VIEW_17X_ANN_CSV} -s ${SNPEFF}/${VIEW_17X_ANN_HTML} >${SNPEFF}/${VIEW_17X_ANN}"
echo
echo bgzip ${SNPEFF}/${VIEW_17X_ANN}
echo
echo bcftools index ${SNPEFF}/${VIEW_17X_ANN}.gz

# Differences in assembly from 17X view
echo
echo make asm_diffs GENOME=${PY17XNL} ASSEMBLY=${PY17X} SAMPLE=17X_PLD DIFFS=${DIFFS} DIFF_VCF=${DIFFS}/${VIEW_17XNL} STATS=${DIFFS}/${VIEW_17XNL_STATS}

# Build snpEff database for 17XNL view
echo
echo java -jar ~/src/snpEff/snpEff.jar build -gff3 -v PY17XNL_PSU
#Annotate vcf
echo "java -jar ~/src/snpEff/snpEff.jar PY17XNL_PSU ${DIFFS}/${VIEW_17XNL}.gz -csvStats ${SNPEFF}/${VIEW_17XNL_ANN_CSV} -s ${SNPEFF}/${VIEW_17XNL_ANN_HTML} >${SNPEFF}/${VIEW_17XNL_ANN}"

echo
echo bgzip ${SNPEFF}/${VIEW_17XNL_ANN}
echo
echo bcftools index ${SNPEFF}/${VIEW_17XNL_ANN}.gz
```
