## Supplemental File 1 for "Long-Read Genome Assembly and Gene Model Annotations for the Rodent Malaria Parasite *Plasmodium yoelii* 17XNL": fix_ann.sh

```
#
# Fix annotations in apicoplast/mitochondria using prokka and rblast
#

ASM_DIR=asm
ASM=17XNL_PSU.fa

REFS=refs
REF_PROT=${REFS}/PlasmoDB-56_Pyoeliiyoelii17X_AnnotatedProteins.fasta

SCRIPTS=scripts/ann_fix
# Chromosome to fix ,API or MIT
CHROM="API"
#CHROM="MIT"

ANN_DIR=ann
OUT_DIR=${ANN_DIR}/${CHROM}_prokka

BRAKER_HYBD_DIR=${ANN_DIR}/braker2_hybrid
BRAKER_NANO_DIR=${ANN_DIR}/braker2_nano

# Resulting output hybrid file
RES_HYBD=${CHROM}_fix_hybrid.gff3

# Resulting output nano file
RES_NANO=${CHROM}_fix_nano.gff3

# Blast results directory
BLAST_DIR=rblast_${CHROM}

# File with gene, transcript and protein id mapping
IDMAP=${REFS}/accession_map_PF17X.txt

# Hybrid Braker annotation file
#BRAKER_HYBD=${BRAKER_HYBD_DIR}/braker2_ann_mod.gff3 
BRAKER_HYBD=${BRAKER_HYBD_DIR}/braker2_ann.gff3 

# Nano Braker annotation file
#BRAKER_NANO=${BRAKER_NANO_DIR}/braker2_ann_mod.gff3
BRAKER_NANO=${BRAKER_NANO_DIR}/braker2_ann.gff3

# Create directory
echo
echo mkdir -p ${OUT_DIR}

#Blast annotations
BLAST_ANN=${BLAST_DIR}/annotated_gene_names.txt

# Header of the chromosome to extract
HEAD="Py17XNL_${CHROM}"

#17X chromosome
REF_HEAD="PY17X_${CHROM}"

# Extract the fasta sequence
echo
echo "samtools faidx ${ASM_DIR}/${ASM} ${HEAD} >${OUT_DIR}/${CHROM}.fa"

# Run prokka
echo
echo prokka --force --addgenes --addmrna ${OUT_DIR}/${CHROM}.fa --outdir ${OUT_DIR}

# PROKKA annotations
echo
PROKKA_ANN=$(ls ${OUT_DIR}/PROKKA*gff)
echo "PROKKA_ANN=$(ls ${OUT_DIR}/PROKKA*gff)"

# Get 17X proteins
echo
echo "cat ${REF_PROT} | grep ${REF_HEAD} | sed 's/ | transcript.*$//' |sed 's/>//' >${OUT_DIR}/17x_${CHROM}_list.txt"

echo
echo "seqtk subseq ${REF_PROT} ${OUT_DIR}/17x_${CHROM}_list.txt >${OUT_DIR}/17x_${CHROM}.fa"

# Link protein files with better names
echo
echo "ls ${OUT_DIR}/*.faa | parallel ln -s {1} py17xnl.fa"
echo ln -s ${OUT_DIR}/17x_${CHROM}.fa py17x.fa

# Make reciprocal blast
echo
echo make reciprocal_blast SOURCE=py17xnl.fa TARGET=py17x.fa IDMAP=${IDMAP} BLAST_OUT=${OUT_DIR}/${BLAST_DIR}

# Remove links
echo 
echo rm -f py17xnl.fa py17x.fa

#
# Fix hybrid annotation
#

# Extract the hybrid braker gff3 for the chromosome to fix
echo
echo "cat ${BRAKER_HYBD} | grep ${CHROM} >${OUT_DIR}/${CHROM}_hybrid.gff3"

# Create a combined gff3 for the chromosome for hybrid annotation
echo
echo "python ${SCRIPTS}/combine_prokka_braker_gff.py ${OUT_DIR}/${BLAST_ANN} ${OUT_DIR}/${CHROM}_hybrid.gff3 ${PROKKA_ANN} >${OUT_DIR}/${RES_HYBD}" 

# Clean up
echo
echo rm -f braker.db prokka.db

#
# Fix nanopore-only annotation
#
# Extract the Nanopore-only braker gff3 for the chromosome to fix
echo
echo "cat ${BRAKER_NANO} | grep ${CHROM} >${OUT_DIR}/${CHROM}_nano.gff3"

# Create a combined gff3 for the chromosome for hybrid annotation
echo
echo "python ${SCRIPTS}/combine_prokka_braker_gff.py ${OUT_DIR}/${BLAST_ANN} ${OUT_DIR}/${CHROM}_nano.gff3 ${PROKKA_ANN} >${OUT_DIR}/${RES_NANO}"

# Clean up
echo
echo rm -f braker.db prokka.db
```
