## Supplemental File 1 for "Long-Read Genome Assembly and Gene Model Annotations for the Rodent Malaria Parasite *Plasmodium yoelii* 17XNL": get_hybrid_v2_fasta.sh

```
set -uex

NANO_PTNS=results/17XNL_PSU_nano_proteins.fa
NANO_CDS=results/17XNL_PSU_nano_cds.fa
NANO_TRANS=results/17XNL_PSU_nano_transcripts.fa
WORK=${ANN}/hybrid_ann_v2


SCRIPTS=scripts
# Create transcript/protein/cds fasta files for hybrid_v2 annotation

# Get transcript/protein ids of added genes from nanopore
cat ${WORK}/added_genes.txt | cut -f 2 >${WORK}/nano.txt

rm -f ${WORK}/nano_added_proteins.txt ${WORK}/nano_added_cds.txt ${WORK}nano_added_transcripts.txt
for i in $(cat nano.txt | grep -v nano_id)
do
echo $i
cat $NANO_PTNS | grep $i |sed -e 's/>//' >>nano_added_proteins.txt
cat $NANO_CDS | grep $i |sed -e 's/>//' >>nano_added_cds.txt
cat $NANO_TRANS | grep $i |sed -e 's/>//' >>nano_added_transcripts.txt

done

rm -f nano_added_proteins.fa nano_added_cds.fa nano_added_transcripts.fa

seqtk subseq -l 60 $NANO_PTNS nano_added_proteins.txt >> nano_added_proteins.fa

seqtk subseq -l 60 $NANO_CDS nano_added_cds.txt >> nano_added_cds.fa

seqtk subseq -l 60 $NANO_TRANS nano_added_transcripts.txt >> nano_added_transcripts.fa

# change the header from nanopore ids to new ids
cat added_genes.txt | cut -f 1,2  >id_map.txt
sed -i 's/nano_id/org_id/' id_map.txt
python ${SCRIPTS}/replace_header.py  nano_added_proteins.fa id_map.txt >added_proteins.fa
python ../../scripts/replace_header.py  nano_added_cds.fa id_map.txt >added_cds.fa
python ../../scripts/replace_header.py  nano_added_transcripts.fa id_map.txt >added_transcripts.fa

# Obtain the sequences of unchanged transcripts/cds/proteins from original hybrid annotation
rm -f hybrid_v2_mRNAs.txt
cat 17XNL_PSU_hybrid_v2.gff3 |awk '$3=="mRNA" {print $9}'| sed -e 's/;.*$//'  -e 's/ID=//'|sort >hybrid_v2_mRNAs.txt

cat added_proteins.fa| grep ">" | sed 's/>//' |sort >added
comm -23 hybrid_v2_mRNAs.txt added >hybrid_uniq_mRNAs.txt

seqtk subseq -l 60 ../../results/17XNL_PSU_hybrid_proteins_v1.fa hybrid_uniq_mRNAs.txt >uniq_proteins.fa

seqtk subseq -l 60 ../../results/17XNL_PSU_hybrid_cds_v1.fa hybrid_uniq_mRNAs.txt >uniq_cds.fa

seqtk subseq -l 60 ../../results/17XNL_PSU_hybrid_transcripts_v1.fa hybrid_uniq_mRNAs.txt >uniq_transcripts.fa

# Combine to create the final fasta files
cat added_proteins.fa uniq_proteins.fa >17XNL_PSU_hybrid_proteins_v2.fa
cat added_cds.fa uniq_cds.fa >17XNL_PSU_hybrid_cds_v2.fa
cat added_transcripts.fa uniq_transcripts.fa >17XNL_PSU_hybrid_transcripts_v2.fa
```
