## Supplemental File 1 for "Long-Read Genome Assembly and Gene Model Annotations for the Rodent Malaria Parasite *Plasmodium yoelii* 17XNL": get_ncRNA.sh

```
ANN=ann
WORK=${ANN}/ncRNA
SRC=scripts/ncRNA
ASSEMBLY=results/17XNL_PSU.fa
REF_GENOME=refs/17X_PLD.fa
REF_GFF=refs/17X_PLD.gff
HYBD_GFF=results/17XNL_PSU_hybrid_v2.gff3
HYBD_NCRNA=${WORK}/17XNL_PSU_hybrid_ncRNA.gff
HYBD_OUT=${WORK}/17XNL_PSU_hybrid_all.gff3

NANO_GFF=results/17XNL_PSU_nano.gff3
NANO_NCRNA=${WORK}/17XNL_PSU_nano_ncRNA.gff
NANO_OUT=${WORK}/17XNL_PSU_nano_all.gff3

# Create Output folder
echo
echo mkdir -p ${WORK}
echo

PREFIX=$(basename ${ASSEMBLY%.fa})
#echo $PREFIX

HYBD_GID=${WORK}/hybrid_last_gids.txt
NANO_GID=${WORK}/nano_last_gids.txt

tRNA_OUT=${WORK}/tRNA
rRNA_OUT=${WORK}/rRNA

echo
echo mkdir -p $tRNA_OUT $rRNA_OUT
echo

# Output files
tRNA_BED=${tRNA_OUT}/${PREFIX}"_tRNA.bed"
rRNA_BED=${rRNA_OUT}/${PREFIX}"_rRNA.bed"
ncRNA_BED=${WORK}/${PREFIX}"_ncRNA.bed"
ncRNA_GFF=${WORK}/${PREFIX}"_ncRNA.gff"

HYBD_OUT=${WORK}/${PREFIX}"_hybrid_all.gff3"
NANO_OUT=${WORK}/${PREFIX}"_nano_all.gff3"

# Find tRNAs
echo
echo bash ${SRC}/get_tRNAs.sh $ASSEMBLY ${tRNA_OUT}
echo

# Find rRNAs
echo
echo bash ${SRC}/get_rRNAs.sh ${REF_GENOME} ${REF_GFF} ${ASSEMBLY} ${rRNA_OUT}
echo
echo "#See notes in ${SRC}/get_rRNAs.sh  for manual modifications"
echo

# Merge tRNA and rRNA bed files.
echo
echo "#Merging bed files and sorting."
echo "cat ${tRNA_BED} ${rRNA_BED}  | sort -k 1,1 -k 2n,2 >${ncRNA_BED}"

#
# Steps for Hybrid model
#


# Get last gene ids per chromosome for hybrid annotation
echo
echo "#Get last gene ids per chromosome for hybrid annotation"
echo "python ${SRC}/get_last_gid.py $HYBD_GFF >$HYBD_GID"
echo

# Create ncRNA  gff for hybrid annotation
echo
echo "#Creating ncRNA gff file."
echo "python ${SRC}/ncRNA_gff.py $ncRNA_BED $HYBD_GID >${ncRNA_GFF}"
echo
echo mv $ncRNA_GFF $HYBD_NCRNA
echo
echo "#Combinig files"
echo "cat $HYBD_GFF $HYBD_NCRNA >${HYBD_OUT}"
echo

# clean up
echo rm -f ann.db


#
# Nanopore model
#

# Get last gene ids per chromosome for hybrid annotation
echo "#Get last gene ids per chromosome for nanopore-only annotation"
echo "python ${SRC}/get_last_gid.py $NANO_GFF >$NANO_GID"
echo

# Create ncRNA  gff for nanopore-only annotation
echo "#Creating ncRNA gff file."
echo "python ${SRC}/ncRNA_gff.py $ncRNA_BED $NANO_GID >${ncRNA_GFF}"
echo
echo mv $ncRNA_GFF $NANO_NCRNA
echo
echo "#Combinig files"
echo "cat $NANO_GFF $NANO_NCRNA >${NANO_OUT}"
echo

# clean up
echo rm -f ann.db
```
