## Supplemental File 1 for "Long-Read Genome Assembly and Gene Model Annotations for the Rodent Malaria Parasite *Plasmodium yoelii* 17XNL": hybrid_v2.sh

```
ANN=ann
WORK=${ANN}/hybrid_ann_v2
TRANS_17X=${WORK}/17X_transcripts.fa
GENOME=refs/17X_PLD.fa
REF_GFF=refs/17X_PLD.gff
ASSEMBLY_IDX=asm/index/17XNL_PSU
TRANS_17X_BAM=${WORK}/17X_transcripts.bam
TRANS_17X_BED=${WORK}/17X_transcripts.bed
SRC=scripts/ann_fix
HYBD_ANN=results/17XNL_PSU_hybrid_v1.gff3
NANO_ANN=results/17XNL_PSU_nano.gff3

OUT=${WORK}/17XNL_PSU_hybrid_v2.gff3

echo mkdir -p ${WORK}

# Map 17X gene-models to 17XNL assembly.
# For this first, create 17X transcripts from the gff3 file
echo
echo make transcripts TRANSCRIPTS=${TRANS_17X} ASSEMBLY=${GENOME} GFF3=${REF_GFF}

# Map 17X transcripts to 17XNL assembly.
echo
echo "hisat2 -x ${ASSEMBLY_IDX} -f ${TRANS_17X}  --sensitive |samtools sort >${TRANS_17X_BAM}"
echo samtools index ${TRANS_17X_BAM}

# Make transcripts bed file
echo
echo "bedtools bamtobed -i ${TRANS_17X_BAM} >${TRANS_17X_BED}"

# Add genes to the hybrid model that do not have the same start and stop-codon as in 17X reference gene from the nanopore model
echo
echo "python ${SRC}/lift_nano_genes.py $HYBD_ANN $NANO_ANN $TRANS_17X_BED >$OUT"

#clean up
echo 
echo rm -f  nano.db  hybrid.db

echo
echo mv hybrid_org_uniq_gids.txt ${WORK}
echo mv duplicated_genes_hybrid_gids.txt ${WORK}
echo mv added_genes.txt  ${WORK}
```
