## Supplemental File 1 for "Long-Read Genome Assembly and Gene Model Annotations for the Rodent Malaria Parasite *Plasmodium yoelii* 17XNL": map.sh

```
REFS=refs
ASM=asm
DATA=data
BAM=bam

PY17XNL=${ASM}/17XNL_PSU.fa
PY17X=${REFS}/PlasmoDB-56_Pyoeliiyoelii17X_Genome.fasta

PY17X_IDX_DIR=${REFS}/index
PY17XNL_IDX_DIR=${ASM}/index

PY17X_IDX=${PY17X_IDX_DIR}/17X_PLD
PY17XNL_IDX=${PY17XNL_IDX_DIR}/17XNL_PSU

CCS_DNA=${DATA}/ccs.fq
ONT_DNA=${DATA}/nano_dna.fq.gz
ILLUM_RNA=${DATA}/WT_all.fq.gz
ONT_RNA=${DATA}/nano_rna.fq.gz

CCS_17X_BAM=ccs_17X_PLD.bam
CCS_17XNL_BAM=ccs_17XNL_PSU.bam

CCS_17X_BGRPH=ccs_17X_PLD.bedgraph
CCS_17XNL_BGRPH=ccs_17XNL_PSU.bedgraph

CCS_17X_BW=ccs_17X_PLD.bw
CCS_17XNL_BW=ccs_17XNL_PSU.bw

ONT_DNA_17X_BAM=ont_dna_17X_PLD.bam
ONT_DNA_17XNL_BAM=ont_dna_17XNL_PSU.bam

ONT_DNA_17X_BGRPH=ont_dna_17X_PLD.bedgraph
ONT_DNA_17XNL_BGRPH=ont_dna_17XNL_PSU.bedgraph

ONT_DNA_17X_BW=ont_dna_17X_PLD.bw
ONT_DNA_17XNL_BW=ont_dna_17XNL_PSU.bw

ONT_RNA_17X_BAM=ont_rna_17X_PLD.bam
ONT_RNA_17XNL_BAM=ont_rna_17XNL_PSU.bam

ILLUM_DNA_17X_BAM=illum_dna_17X_PLD.bam
ILLUM_DNA_17XNL_BAM=illum_dna_17XNL_PSU.bam

ILLUM_RNA_17X_BAM=illum_rna_17X_PLD.bam
ILLUM_RNA_17XNL_BAM=illum_rna_17XNL_PSU.bam

CCS_TAG=map-pb
ONT_TAG=map-ont

# Create directory
echo
echo mkdir -p ${BAM}

#  Map ccs DNA to 17X reference
echo
echo "#Map ccs DNA to 17X reference"
echo
echo make minimap2_dna GENOME=${PY17X} TAG=${CCS_TAG} INPUT=${CCS_DNA} OUTPUT=${BAM}/${CCS_17X_BAM} SAMPLE=17XNL_PSU_ccs
echo
echo make coverage GENOME=${PY17X} INBAM=${BAM}/${CCS_17X_BAM} BEDGRAPH=${BAM}/${CCS_17X_BGRPH} BIGWIG=${BAM}/${CCS_17X_BW}

#  Map ccs DNA to 17XNL assembly
echo
echo "#  Map ccs DNA to 17XNL assembly"
echo
echo make minimap2_dna GENOME=${PY17XNL} TAG=${CCS_TAG} INPUT=${CCS_DNA} OUTPUT=${BAM}/${CCS_17XNL_BAM} SAMPLE=17XNL_PSU_ccs
echo
echo make coverage GENOME=${PY17XNL} INBAM=${BAM}/${CCS_17XNL_BAM} BEDGRAPH=${BAM}/${CCS_17XNL_BGRPH} BIGWIG=${BAM}/${CCS_17XNL_BW}

# Map ont DNA to 17X reference
echo
echo "# Map ont DNA to 17X reference"
echo
echo make minimap2_dna GENOME=${PY17X} TAG=${ONT_TAG} INPUT=${ONT_DNA} OUTPUT=${BAM}/${ONT_DNA_17X_BAM} SAMPLE=17XNL_PSU_ont
echo
echo make coverage GENOME=${PY17X} INBAM=${BAM}/${ONT_DNA_17X_BAM} BEDGRAPH=${BAM}/${ONT_DNA_17X_BGRPH} BIGWIG=${BAM}/${ONT_DNA_17X_BW}

# Map ont DNA to 17XNL_PSU
echo
echo "# Map ont DNA to 17XNL_PSU"
echo
echo make minimap2_dna GENOME=${PY17XNL} TAG=${ONT_TAG} INPUT=${ONT_DNA} OUTPUT=${BAM}/${ONT_DNA_17XNL_BAM}  SAMPLE=17XNL_PSU_ont
echo
echo make coverage GENOME=${PY17XNL} INBAM=${BAM}/${ONT_DNA_17XNL_BAM} BEDGRAPH=${BAM}/${ONT_DNA_17XNL_BGRPH} BIGWIG=${BAM}/${ONT_DNA_17XNL_BW}

# Map illum_dna to 17X reference
echo
echo "# Map illum_dna to 17X reference"
echo
echo make bwa_index INDEX_DIR=${PY17X_IDX_DIR} INDEX=${PY17X_IDX} ASSEMBLY=${PY17X}
echo make map_illum_dna SAMPLE=17XNL_illum_dna INDEX=${PY17X_IDX} OUTPUT=${BAM}/${ILLUM_DNA_17X_BAM}

# Map illum_dna to 17XNL assembly
echo
echo "# Map illum_dna to 17XNL assembly"
echo
echo make bwa_index INDEX_DIR=${PY17XNL_IDX_DIR} INDEX=${PY17XNL_IDX} ASSEMBLY=${PY17XNL}
echo make map_illum_dna SAMPLE=17XNL_illum_dna INDEX=${PY17XNL_IDX} OUTPUT=${BAM}/${ILLUM_DNA_17XNL_BAM}

# Map ont RNA to 17X reference
echo
echo "# Map ont RNA to 17X reference"
echo
echo make minimap2_rna ASSEMBLY=${PY17X} NANO_RNA=${ONT_RNA} OUTPUT=${BAM}/${ONT_RNA_17X_BAM} SAMPLE=17XNL_PSU_ont

# Map ont RNA to 17XNL assembly
echo
echo "# Map ont RNA to 17XNL assembly"
echo
echo make minimap2_rna ASSEMBLY=${PY17XNL} NANO_RNA=${ONT_RNA} OUTPUT=${BAM}/${ONT_RNA_17XNL_BAM} SAMPLE=17XNL_PSU_ont

# Map illum_RNA to 17X reference
# Make hisat2 indices
echo
echo "# Map illum_RNA to 17X reference"
echo
echo make hisat2_index ASSEMBLY=${PY17X} INDEX=${PY17X_IDX} INDEX_DIR=${PY17X_IDX_DIR}
echo
echo make hisat2 INDEX=${PY17X_IDX} ILLUM_RNA=$ILLUM_RNA OUTPUT=${BAM}/${ILLUM_RNA_17X_BAM}

# Map illum_RNA to 17XNL assembly
# Make hisat2 indices
echo
echo "# Map illum_RNA to 17XNL assembly"
echo
echo make hisat2_index ASSEMBLY=${PY17XNL} INDEX=${PY17XNL_IDX} INDEX_DIR=${PY17XNL_IDX_DIR}
echo
echo make hisat2 INDEX=${PY17XNL_IDX} ILLUM_RNA=$ILLUM_RNA OUTPUT=${BAM}/${ILLUM_RNA_17XNL_BAM}
```
