## Supplemental File 1 for "Long-Read Genome Assembly and Gene Model Annotations for the Rodent Malaria Parasite *Plasmodium yoelii* 17XNL": prep_cons.sh

```
# 
# Modify the headers and repeatmask cons2.fa 
#

# Rename header
ASM=asm
sed -i 's/Py17X/Py17XNL/' ${ASM}/cons2.fa
sed -i 's/_v3//' ${ASM}/cons2.fa

# Mask the genome
tantan -m scripts/atMask.mat  -r 0.01 ${ASM}/cons2.fa >${ASM}/17XNL_PSU.fa
samtools faidx ${ASM}/17XNL_PSU.fa
```
