## Supplemental File 1 for "Long-Read Genome Assembly and Gene Model Annotations for the Rodent Malaria Parasite *Plasmodium yoelii* 17XNL": rblast_ann.sh

```
REFS=refs
ASM_DIR=asm
ANN_DIR=ann
RES_DIR=results

ASM=${ASM_DIR}/17XNL_PSU.fa

BRAKER_DIR=${ANN_DIR}/braker2_hybrid
BLAST_DIR=${ANN_DIR}/rblast_res_hybrid

TRANS=${RES_DIR}/17XNL_PSU_hybrid_transcripts.fa
CDS=${RES_DIR}/17XNL_PSU_hybrid_cds.fa
PROT=${RES_DIR}/17XNL_PSU_hybrid_proteins.fa

#BRAKER_DIR=${ANN_DIR}/braker2_nano
#BLAST_DIR=${ANN_DIR}/rblast_res_nano

#TRANS=${RES_DIR}/17XNL_PSU_nano_transcripts.fa
#CDS=${RES_DIR}/17XNL_PSU_nano_cds.fa
#PROT=${RES_DIR}/17XNL_PSU_nano_proteins.fa

BRAKER_GFF3=${BRAKER_DIR}/augustus.hints_utr.gff3
BRAKER_PROT=${BRAKER_DIR}/augustus.hints_utr.aa
BRAKER_CDS=${BRAKER_DIR}/augustus.hints_utr.codingseq
BRAKER_ANN=${BRAKER_DIR}/braker2_ann.gff3
BRAKER_TMAP=${BLAST_DIR}/transcript_map.txt

# Blast Database file #Py17X proteins fasta
PY17X_PROT=${REFS}/PlasmoDB-56_Pyoeliiyoelii17X_AnnotatedProteins.fasta 

# File with gene, transcript and protein id mapping
IDMAP=${REFS}/accession_map_PY17X.txt

SOURCE=py17xnl.fa 
TARGET=py17x.fa


# Prepare blast files with short names
echo
echo ln -s ${BRAKER_PROT} ${SOURCE}
echo ln -s ${PY17X_PROT} ${TARGET}

# Run rblast
echo
echo make reciprocal_blast SOURCE=${SOURCE} TARGET=${TARGET} IDMAP=${IDMAP} BLAST_OUT=${BLAST_DIR} 

# Annotate gff3 with rblast results
echo 
echo make annotate_GFF3 BLAST_OUT=${BLAST_DIR} BRAKER_GFF3=${BRAKER_GFF3} BRAKER_ANN=${BRAKER_ANN}

#Clean up
echo
echo rm -f ${SOURCE} ${TARGET}

# Create transcript fasta file
echo
echo make transcripts TRANSCRIPTS=${TRANS} ASSEMBLY=${ASM} GFF3=${BRAKER_ANN}

# Create CDS fasta file
echo
echo make cds BRAKER_CDS=${BRAKER_CDS} CDS=${CDS} BRAKER_TMAP=${BRAKER_TMAP}

# Create protein fasta file
echo
echo make proteins BRAKER_PROT=${BRAKER_PROT} PROTEINS=${PROT} BRAKER_TMAP=${BRAKER_TMAP}
```
