## Supplemental File 1 for "Long-Read Genome Assembly and Gene Model Annotations for the Rodent Malaria Parasite *Plasmodium yoelii* 17XNL": submission.sh

```
set -ue

#foramt GFF3 file for submission
WORK=ann/submission
HYBD_GFF3=results/17XNL_PSU_hybrid_v3.gff3
NANO_GFF3=results/17XNL_PSU_nano_v2.gff3
SORTED=sorted.gff3
SCRIPTS=scripts/misc
ACC_MAP=ann/accession_map_PY17X_desc.txt
CHROMS=ann/chroms.txt
HYBD_OUT=${WORK}/17XNL_PSU_hybrid_v4.gff3
NANO_OUT=${WORK}/17XNL_PSU_nano_v3.gff3

# Change these variables

#GFF3=$HYBD_GFF3
#OUT=$HYBD_OUT

GFF3=$NANO_GFF3
OUT=$NANO_OUT


echo
echo mkdir -p ${WORK}


#Separate the rows with locus tag from those without the locus-tag
echo
echo "cat ${GFF3} |grep "locus_tag" >${WORK}/locus_tag.gff3"
echo "cat ${GFF3} |grep -v "locus_tag" >${WORK}/no_locus_tag.gff3"

#
echo
echo conda activate gt_env

# Sort GFF3
echo 
echo make sort_gff3 GFF3=${WORK}/no_locus_tag.gff3 SORTED_GFF3=${WORK}/${SORTED}

#
# Reactivate bioinfo env
echo
echo conda activate bioinfo

# Format GFF3
echo
echo "python ${SCRIPTS}/format_gff.py ${WORK}/${SORTED} $ACC_MAP $CHROMS >${WORK}/formatted.gff3"
echo "cat ${WORK}/formatted.gff3 ${WORK}/locus_tag.gff3 >${OUT}"

# Change ncRNA_gene to 'gene' feature
echo
echo sed -i \'s/ncRNA_gene/gene/\' ${OUT}
echo

exit
#
# Reactivate gt env
echo
echo conda activate gt_env

# Sort GFF3 again
echo 
echo make sort_gff3 GFF3=${WORK}/all.gff3 SORTED_GFF3=${OUT}

#
# Reactivate bioinfo env
echo
echo conda activate bioinfo

#Cleanup
echo
echo rm -f ${WORK}/locus_tag.gff3 ${WORK}/no_locus_tag.gff3 ${WORK}/${SORTED}
```
