## Supplemental File 1 for "Long-Read Genome Assembly and Gene Model Annotations for the Rodent Malaria Parasite *Plasmodium yoelii* 17XNL": utr_len.sh

```
# set -uex
# Generate UT lengths

HYBD_GFF3=results/17XNL_PSU_hybrid_v4.gff3
NANO_GFF3=results/17XNL_PSU_nano_v3.gff3
GENE_MATCH_HYBRID=ann/gene_match_hybrid.txt
GENE_MATCH_NANO=ann/gene_match_nano.txt


# Change these variables for nano
INPUT=$HYBD_GFF3
TAG=hybrid
GMATCH=$GENE_MATCH_HYBRID

#INPUT=$NANO_GFF3
#TAG=nano
#GMATCH=$GENE_MATCH_NANO

ANN=ann
WORK=ann/UTRS
SRC=scripts/misc

# CReate directory
echo
echo mkdir -p $WORK

## Sort the input gff3 file and add locus tag

#echo "# genome tools require python 2.7"
#echo "# conda create -n gt_env -y python=2.7"
#echo " # conda install -c bioconda genometools-genometools"
#echo "conda activate gt_env"
#echo "gt gff3 -sort -retainids ${INPUT} >${WORK}/sorted.gff3"

##Reactivate bioinfo environment
#echo 
#echo "conda activate bioinfo"

## Add locus tag
#echo
#echo "python scripts/format_gff.py  ${WORK}/sorted.gff3 ${ANN}/accession_map_PY17X_desc.txt ${ANN}/chroms.txt >${WORK}/input.gff3"

echo 
echo cp ${INPUT} ${WORK}/input.gff3

# Extract hybrid 5p UTRs
echo
echo "cat ${WORK}/input.gff3 | awk '\$3==\"five_prime_UTR\" {print}' >${WORK}/${TAG}_5P_UTR.gff3"

# Extract hybrid 3p UTRs
echo
echo "cat ${WORK}/input.gff3 | awk '\$3==\"three_prime_UTR\" {print}' >${WORK}/${TAG}_3P_UTR.gff3"

#Make a table  of 5p UTRs with length
echo
echo "python ${SRC}/attributes_gff.py ${WORK}/${TAG}_5P_UTR.gff3 five_prime_UTR transcript_id:gene_name:locus_tag | sed -e 's/transcript_id=//' -e 's/gene_name=//' -e 's/locus_tag=//' |awk '{ulen=\$3-\$2+1; print \$0\"\t\"ulen}' >${WORK}/${TAG}_5P_UTR_len.txt"

#Make a table  of 3p UTRs with length
echo
echo "python ${SRC}/attributes_gff.py ${WORK}/${TAG}_3P_UTR.gff3 three_prime_UTR transcript_id:gene_name:locus_tag | sed -e 's/transcript_id=//' -e 's/gene_name=//' -e 's/locus_tag=//' | awk '{ulen=\$3-\$2+1; print \$0\"\t\"ulen}' >${WORK}/${TAG}_3P_UTR_len.txt"

# Add 17X gene ids to the table
echo
echo "python ${SRC}/add_gids.py $GMATCH ${WORK}/${TAG}_5P_UTR_len.txt 5 >${WORK}/${TAG}_5P_UTR_len_17xgids.txt"

echo
echo "python ${SRC}/add_gids.py $GMATCH ${WORK}/${TAG}_3P_UTR_len.txt 5 >${WORK}/${TAG}_3P_UTR_len_17xgids.txt"
```
